# Multiplexing cell-cell communication

**DOI:** 10.1101/584664

**Authors:** John T. Sexton, Jeffrey J. Tabor

## Abstract

The engineering of advanced multicellular behaviors, such as the programmed growth of biofilms or tissues, requires cells to communicate multiple aspects of physiological information. Unfortunately, few cell-cell communication systems have been developed for synthetic biology. Here, we engineer a genetically-encoded channel selector device that enables a single communication system to transmit two separate intercellular conversations. Our design comprises multiplexer and demultiplexer sub-circuits constructed from a total of 12 CRISPRi-based transcriptional logic gates, an acyl homoserine lactone-based communication module, and three inducible promoters that enable small molecule control over the conversations. Experimentally-parameterized mathematical models of the sub-components predict the steady state and dynamical performance of the full system. Multiplexed cell-cell communication has applications in synthetic development, metabolic engineering, and other areas requiring the coordination of multiple pathways amongst a community of cells.

**One Sentence Summary:** We have engineered a synthetic genetic system that enables bacteria to have two separate conversations over a single chemical “wire” by separating the conversations in time.

## Main Text

Synthetic biologists have long aimed to engineer cells to cooperate to perform complex tasks. For example, cell communities have been programmed to undergo synchronized gene expression dynamics (*1*–*5*), act as distributed computers (*6*–*8*), and differentiate into simple patterns (*9*–*13*). However, due to challenges such as molecular cross-talk, no more than two communication systems have been combined in a single design (*14*–*16*). In stark contrast, evolution utilizes dozens of communication signals to orchestrate complex behaviors such as embryo patterning (*17*), the precise wiring of the nervous (*18*), circulatory and lymphatic systems (*19*), and innate and adaptive immunity.

In electronic engineering, channel selectors (CSs) enable a single communication resource such as a wire to transmit multiple conversations. In a CS, a multiplexer circuit (MUX) is linked to a demultiplexer circuit (DEMUX), and both are controlled by a common SELECT signal (Fig. 1, A **and** B). The MUX receives *n* input signals, but propagates only one to the DEMUX, depending on the SELECT value. The DEMUX relays this signal to one of *n* possible outputs, again depending on SELECT. A conversation occurs because a given input (IN_*i*_) is routed exclusively to the corresponding output (OUT_*i*_). Multiple conversations can occur sequentially if the SELECT value is changed (Fig. 1, A **and** B).

**Figure 1.**
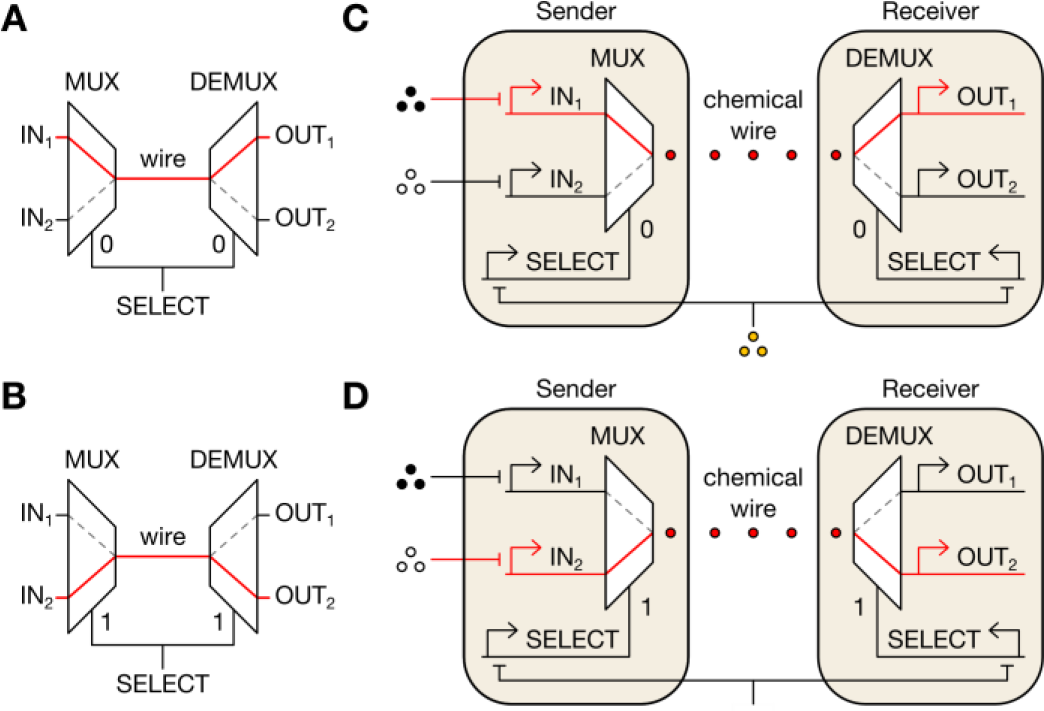
Electronic and biological channel selectors. (**A** and **B**) In an electrical CS, the SELECT signal directs the MUX to choose an input to transmit across the wire and the DEMUX to route the signal to the corresponding output. The CS can thus transmit two separate conversations over the same wire by changing the value of SELECT. (**C** and **D**) Design of a genetically-encoded CS that enables transmission of multiple conversations via a single cell-cell communication system (red circles). Here, IN_1_, IN_2_, and SELECT are transcriptional signals repressed by chemically-inducible promoters and their ligands (black, white, and orange circles).

To multiplex cell-cell communication (Fig. 1C **and** D), we implemented the simplest CS, where *n* = 2, genetically. First, we used formal logic synthesis to design the required 2-input MUX and 2-output DEMUX from the smallest possible number of Boolean NOT and NOR gates (**Fig. S1A**). We restricted our design to NOT and NOR because these gates can be combined to achieve any digital logic operation and readily constructed in live cells using transcriptional repressors and repressible promoters (*8*, *20*–*22*). Our design process yielded minimized MUX and DEMUX circuits composed of one NOT and three NOR gates assembled in three layers (**Fig. S1, B and C**), and two NOT and two NOR gates assembled in two layers (**Fig. S1, D and E**), respectively.

To implement these circuits genetically, we turned to the CRISPR interference (CRISPRi) technology, wherein a nuclease-dead Cas9 (dCas9):small guide RNA (sgRNA) complex sequence-specifically binds and represses transcription from a target promoter (*23*). We designed nine putatively orthogonal sgRNA:promoter pairs by encoding randomized and divergent operator sequences lacking homology to the *E. coli* genome (**Methods**) between the −35 and −10 sites of otherwise constitutive promoters (Figs. 2A **and** B). Indeed, all sgRNAs in our set (S1-S9) strongly repress their cognate promoters (P1-P9), without cross-repressing any of the eight non-cognate promoters (**Fig. 2C**).

**Figure 2.**
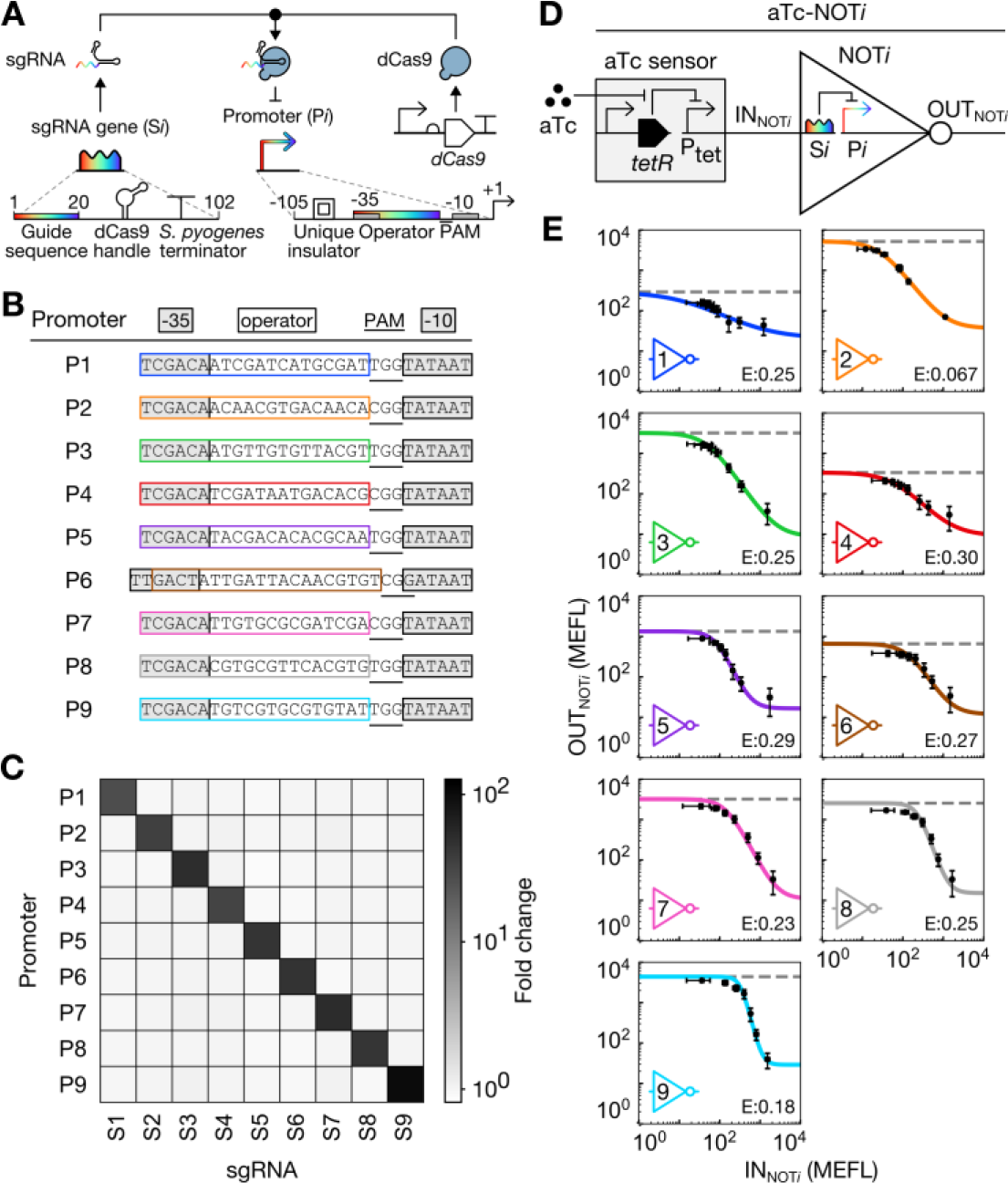
CRISPRi-based NOT gate library. (**A**) Schematic of our engineered sgRNA:promoter pairs. Rainbows indicate specificity-determining regions with variable sequences. (**B**) Core sequences of P1-P9. (**C**) Orthogonality of S*i*:P*i* pairs. Each sgRNA was strongly induced and the resulting fold change in sfGFP fluorescence from each promoter was measured. (**D**) Schematic of aTc-NOT*i* circuits used to measure NOT gate transfer functions. (**E**) Transfer functions of NOT1-NOT9. Bacteria were grown in different aTc concentrations. To calculate IN_NOT*i*_ and OUT_NOT*i*_ values, the mean sfGFP fluorescence of input and output probe strains were calculated and averaged across three replicates measured on different days. Error bars represent the standard error of the mean (s.e.m.) of mean sfGFP fluorescence. Dashed lines indicate the maximum gate output in the absence of the sgRNA gene. Colored lines represent model fits. Fit parameters are shown in **Table S1**. Error (E) represents the RMSE between the fit and data and is expressed in MEFL decades.

Each S*i*:P*i* pair constitutes a transcriptional NOT gate (NOT*i*), and inverts a high or low transcriptional input signal (IN_NOT*i*_) into low or high transcriptional output signal (OUT_NOT*i*_), respectively (Fig. 2, D **and** E). To characterize each sub-component in our system, we designed a set of standard probe plasmids, wherein a promoter of interest drives transcription of an insulated superfolder green fluorescent protein (*sfgfp*) reporter gene (**Fig. S2**). To generate a wide range of IN_NOT*i*_ signals, we placed transcription of S*i* under control of an anhydrotetracycline (aTc)-inducible promoter system (i.e. an aTc sensor; **Fig. 2D**) in the context of NOT*i*. Then, we co-transformed each aTc-NOT*i* gate into *E. coli* pairwise with the aTc sensor probe plasmid and the corresponding P*i* probe plasmid. We then exposed these 18 strains to different aTc concentrations, and quantified the resulting IN_NOT*i*_ and OUT_NOT*i*_ signals via sfGFP fluorescence in calibrated Molecules of Equivalent Fluorescein (MEFL) units (**Fig. S2A, S2B**). Most of the resulting IN_NOT*i*_/OUT_NOT*i*_ relationships (i.e. NOT*i* transfer functions) are well fit by a constrained Hill model (*n*=1), consistent with a recent study (*24*) (**Fig. S3**). However, an unconstrained Hill model better describes several of the transfer functions (**Fig. S3**). Thus, we chose to use unconstrained Hill models to describe the behaviors of all NOT*i* gates (Fig. 2E, **Table S1**). The root mean square error (RMSE) between the data and fits range between 0.067-0.30 MEFL decades (**Methods**).

Each NOT*i* gate can be converted into a corresponding NOR gate (i.e. NOR*i*) by adding a second instance of the sgRNA transcribed from a second, independent input promoter (**Fig. S4A**). NOR gates produce high output only when both inputs are low. We simulated the 2-input/1-output transfer functions of NOR*i* by adding a second transcriptional input term to each NOT*i* model (**Supplementary Materials**). Indeed, all nine NOR*i* gates are predicted to exhibit NOR-like behavior (**Fig. S4B**). We hypothesized that our library of NOT and NOR gates could be utilized to construct the MUX and DEMUX.

We selected NOR5, NOR3, NOT2, and NOR6 to implement the MUX (**Fig. 3A**). Considering only low (0) and high (1) signal values, a 2-input MUX can receive eight combinations of IN_1_, IN_2_, and SELECT. To generate these eight input combinations, we constructed eight MUX test circuits wherein constitutive promoters are absent (resulting in low signal) or present (resulting in high signal) in the appropriate circuit locations (**Fig. 3A**). Then, we probed the output of each gate and the overall MUX behavior in all eight test circuits. First, all gates propagate digital-like signals, generating only the minimum or maximum output value in all conditions (**Fig. 3A**). Second, all gates perform the proper computation in all test circuits (**Fig. 3A**). As a result, the MUX correctly chooses to relay the IN_1_ signal (to OUT_MUX_) when SELECT = 0 and IN_2_ when SELECT = 1 (**Fig. 3A**). Finally, all gate outputs are predicted accurately by a MUX model constructed from the individual gate models (RMSE between model predictions and data = 0.33 MEFL decades) (Fig. 3A, **Supplementary Materials**).

**Figure 3.**
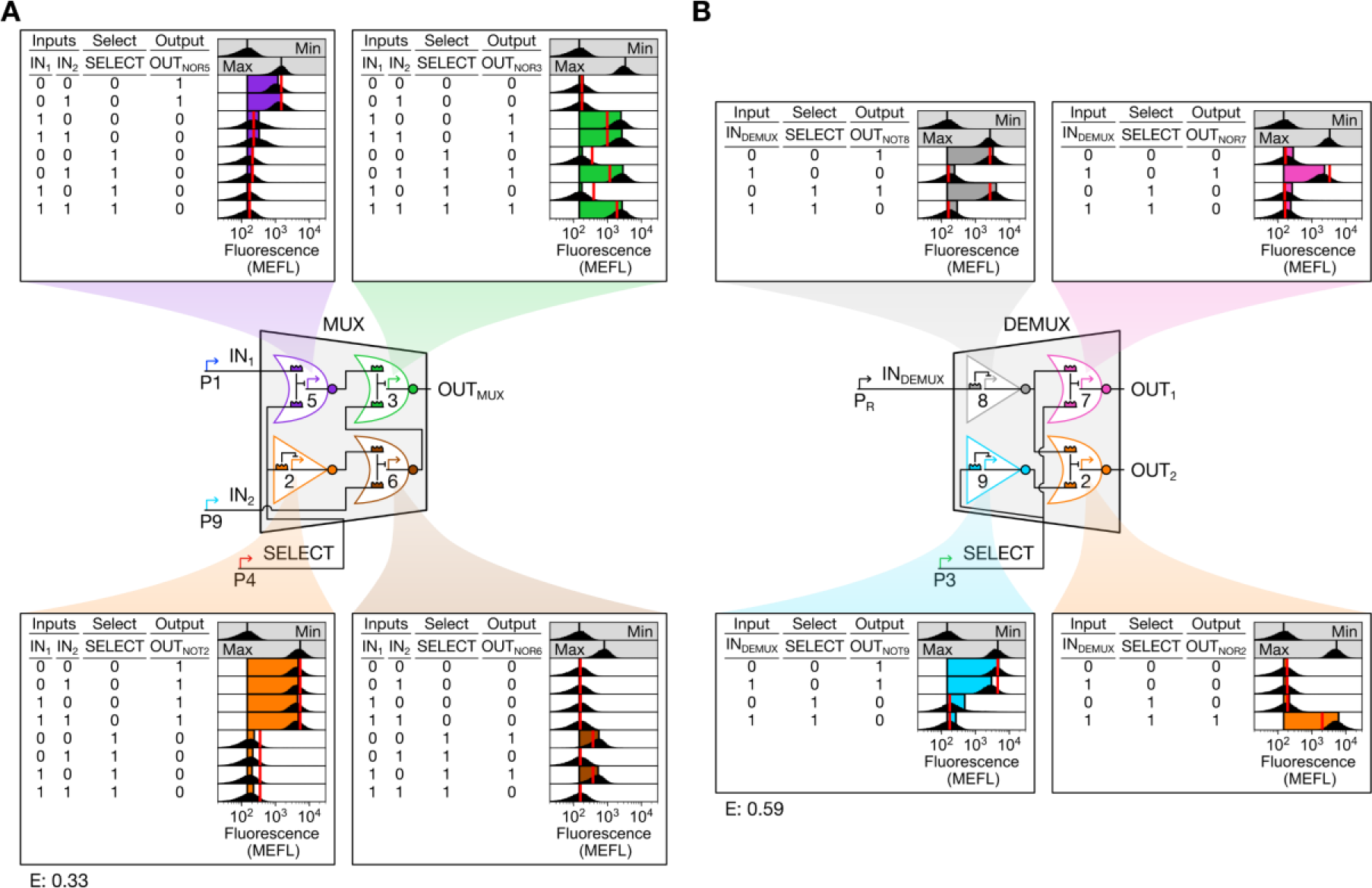
Biological MUX and DEMUX circuits. (**A**) MUX design and characterization. (**B**) DEMUX design and characterization. The strong promoter P_R_ was selected to generate IN_DEMUX_ after the weaker promoter P1 failed to generate correct DEMUX behavior. Flow cytometry fluorescence distributions (black), model-simulated mean fluorescence (red lines), and experimental mean sfGFP fluorescence (colored bars) are shown. Mean sfGFP fluorescence bars are bounded at left by mean autofluorescence and at right by mean fluorescence. RMSE between simulated and measured mean sfGFP fluorescence (E) is expressed in MEFL decades. Min and max indicate cellular autofluorescence and gate output in the absence of sgRNA, respectively (vertical lines indicate mean). Min was measured in triplicate on three separate days, max was measured once on a fourth day, and all other measurements were performed on a fifth day.

We similarly constructed the DEMUX from NOT8, NOR7, NOT9, and NOR2 (**Fig. 3B**). A 2-output DEMUX can receive four combinations of input (IN_DEMUX_) and SELECT signal values. We tested our DEMUX design using four test circuits, as before. All gates behave digitally, perform the proper computations, and are well predicted by a model (RMSE = 0.59 MEFL decades) (Fig. 3B, **Supplementary Materials**). Indeed, the DEMUX correctly relays IN_DEMUX_ to OUT_1_ when SELECT = 0, and to OUT_2_ when SELECT = 1 (**Fig. 3B**). Thus, our MUX and DEMUX both function as designed.

Ideally, a biological CS should interface with regulated promoters – those whose activities vary in response to perturbations. To demonstrate this capability, we first constructed the SENSOR-MUX, wherein the aTc sensor controls IN_1_, an isopropyl β-D-1-thiogalactopyranoside (IPTG) sensor controls IN_2_, and a 2,4-diacetylphloroglucinol (DAPG) sensor controls SELECT each via an additional sgRNA (**Figs. S5, S6**). When DAPG is present (SELECT = 0), the behavior of the SENSOR-MUX recapitulates that of the MUX (**Fig. S6**). However, two faults arise when DAPG is absent (SELECT = 1). First, when aTc is present (IN_1_ = 0) and IPTG is absent (IN_2_ = 1), only ~40% of cells propagate IN_2_ to OUT_MUX_ (**Fig. S6**). This fault is caused by leaky transcription from the DAPG sensor in ~60% of cells (**Fig. S6**). We chose not to debug this fault because it is a failure of the DAPG sensor rather than the MUX, and we hypothesized that production of the cell-cell communication signal from ~40% of cells would be sufficient to transmit information to the DEMUX. In the second fault, when IPTG is present (IN_2_ = 0) OUT_MUX_ reaches only intermediate, rather than low, levels (**Fig. S6**). This result indicates that NOR6 produces too low an output signal to fully repress NOR3 in the context of SENSOR-MUX. We corrected this fault by increasing the strength of P6, yielding NOR6* and SENSOR-MUX* (**Figs. S7 and S8**). The performance of SENSOR-MUX* matches a model based on the performance of the individual sensors and gates (RMSE = 0.40 MEFL decades) (**Fig. S8**).

For cell-cell communication, we chose the widely-used 3-oxohexanoyl acylhomoserine lactone (AHL) system (**Fig. S9**). First, we constructed AHL-DEMUX wherein an AHL sensor controls IN_DEMUX_, and DAPG again controls SELECT (**Fig. S10**). Probe experiments confirm that AHL-DEMUX computes proper outputs for all four possible AHL and DAPG input combinations, and the behavior of every sensor and gate agrees with model predictions (RMSE = 0.68 MEFL decades) (**Fig. S10, Supplementary Materials**). Finally, we built SENSOR-MUX*-AHL wherein OUT_MUX_ controls production of the AHL biosynthetic enzyme LuxI (**Fig. 4**).

**Figure 4.**
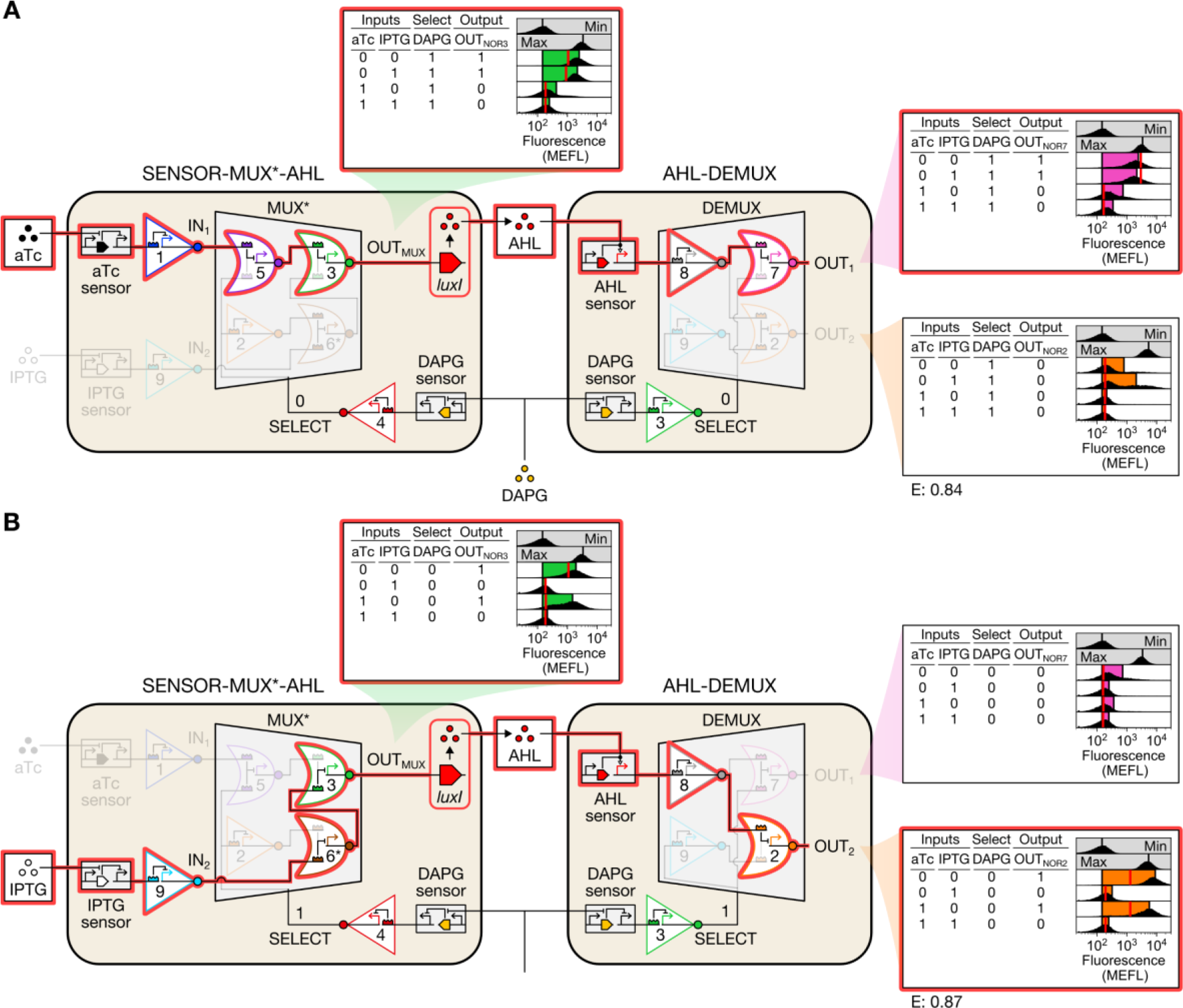
Multiplexing cell-cell communication. (**A**) Conversation 1. In the presence of DAPG, low aTc results in high IN_1_, which is routed to OUT_1_ via AHL. (**B**) Conversation 2. In the absence of DAPG, low IPTG results in high IN_2_, which is routed to OUT_2_ via AHL. Conversations are highlighted in red. Min was measured in triplicate on three separate days, max was measured once on a fourth day, and all other measurements were performed on a fifth day.

We implemented the full CS by growing SENSOR-MUX*-AHL cells with the eight combinations of aTc, IPTG, and DAPG and then diluting them into new cultures with AHL-DEMUX cells and fresh inducers (**Fig. S11**). In each case, we monitored the conversation by probing OUT_MUX_, OUT_1_, and OUT_2_. Indeed, when DAPG is present, only conversation 1 occurs: the presence of aTc is relayed through both circuits and cell strains, ultimately controlling OUT_1_ (**Fig. 4A**). Furthermore, when DAPG is absent, conversation 2 occurs instead: IPTG is relayed through both circuits and strains, and controls OUT_2_ (**Fig. 4B**). Thus, our CS switches between two different conversations in response to DAPG.

While the performance of the CS largely recapitulates the individual behaviors of SENSOR-MUX* and AHL-DEMUX (Figs. 4, **S8, and S10**), we observed that aTc erroneously activates OUT_2_ in ~40% of cells in the presence of DAPG (**Fig. 4A**). We determined that AHL concentrations of 5 nM or greater induce this fault in AHL-DEMUX cells (**Fig. S12, A and B**). We hypothesized this fault may arise due to high total sgRNA levels in AHL-DEMUX cells resulting in dCas9 saturation. To investigate this possibility, we calculated the total sgRNA expressed by SENSOR-MUX* and AHL-DEMUX for all input conditions. The AHL-DEMUX produces >27,000 MEFL total sgRNA under fault-inducing conditions, whereas both circuits produce <17,000 MEFL total sgRNA otherwise (**Fig. S12C**). AHL-induced S8 overexpression may therefore outcompete S2 for dCas9, causing NOR2 to fail. We were unable to increase dCas9 expression further due to toxicity. However, a recent dCas9 variant that can be expressed to higher levels may alleviate this issue (*24*).

The longest computation path through the CS comprises eight sequential layers: the DAPG sensor, NOT4, NOT2, NOR6*, NOR3, the AHL system, NOT8, and NOR7 (**Fig. S13, A and B**). We characterized the dynamics of signal propagation through this path by adding DAPG to SENSOR-MUX*-AHL cells, AHL to AHL-DEMUX cells, and probing the outputs of each layer over time. Sensors activated rapidly, with their reporters approaching steady state within ~1 h. Gate reporters responded in order, requiring an average of 1.3 h to propagate through each layer (**Fig. S13, E and F**). The SENSOR-MUX*-AHL and AHL-DEMUX cells completed their responses in 5 and 7 h, respectively (**Fig. S13, C and D**). A gene expression dynamics model assuming stable dCas9:sgRNA complexes closely predicts these dynamics (RMSE = 0.45 MEFL decades) (**Fig. S13, C and D, Supplementary Materials**).

The next generation of engineered multicellular behaviors will require more than two cell-cell communication channels (*25*). Genetically-encoded CSs can enable such applications by reusing communication systems. As the number of transcriptional logic gates that can be implemented in a single cell continues to increase, CSs that can switch between three or more conversations can be envisioned (**Fig. S14**). Computation times could be substantially accelerated using de-stabilized (e.g. proteolysis-tagged) repressors. Eventually, CSs could transmit information on the seconds timescale using logic gates based upon post-translational, rather than transcriptional regulation (*26*, *27*). Such advances will enable major progress toward long-standing goals of synthetic biology.

## Acknowledgements

We thank Evan Olson and Sebastián Castillo-Hair for helpful discussions on genetic circuit design, Joel Moake for use of his flow cytometer, Matthew Bennett for gifting us a plasmid harboring *cas9*, and Sebastián Castillo-Hair for constructing the dCas9-expressing pSC31_1 and pSC31_3 plasmids.

## Author contributions

JTS and JJT conceived of the project and designed the experiments. JTS performed the experiments and analyzed the data. JTS and JJT wrote the paper.

## Supplementary Materials

Materials and Methods

Supplementary Text

Figs. S1 to S16

Tables S1 to S17

References (28 - 52)

