## Supplementary Materials for "Multiplexing cell-cell communication"

### Materials and Methods

Design of sgRNA:promoter pairs

We designed our sgRNAs to specifically repress their cognate promoters without cross-repressing essential *E. coli* MG1655 genes. To this end, we first generated random pre-sequences encoding the 3'-most 14 bp of a CRISPRi operator, which includes the specificity-conveying seed region, followed by 1 bp encoding the first (degenerate) nucleotide of the adjacent PAM site (NGG). We biased these pre-sequences for ~40% GC content. We then appended two guanine residues to the 3' end of each pre-sequence to complete the PAM site. Then, we designed two different promoter sequences from each pre-sequence:GG by flanking it with -35 and -10 sites from apFAB126 (*28*) and λ $P_R$. We designed one sgRNA for each promoter by encoding six nucleotides matching the -35 site, 14 nucleotides matching the 5'-end of each pre-sequence, and 82 nucleotides encoding the *S. pyogenes* Cas9 handle and terminator (**Fig. 2A**). Designed promoters containing more than one additional primary and any additional secondary (NAG) PAM sites relative to the -35 and -10 sequences from which they were made were discarded, as were other promoters and sgRNAs derived from the same pre-sequence. Promoters containing BsaI restriction sites, which interfere with Golden Gate cloning, were also discarded along with related promoters and sgRNAs.

We then computationally screened each sgRNA for off-target binding to the *E. coli* MG1655 genome (GenBank U00096.3). First, we used custom Python scripts to search for genome sequences overlapping essential genes and matching the 3'-most ≥ 10 nucleotides of the operator:PAM sequence from each promoter. Essential genes were identified by querying all MG1655 gene features against the EcoCyc *E. coli* database (https://ecocyc.org/). If cell growth was reported to be arrested when the gene was knocked out in any growth medium, the gene was considered essential. Genome features of type 'rRNA', 'tRNA', and 'rep_origin' were also considered essential. If overlap of an essential feature was detected, the sgRNA:promoter pair was discarded, along with other pairs sharing a pre-sequence. Next, we searched for genome sequences matching the 3'-most ≥ 12 nucleotides of the operator:PAM sequence and again discarded any matches and pairs sharing a pre-sequence. Finally, we used the CasValue algorithm to identify sgRNAs likely to bind the genome. CasValue scores sgRNA:genome binding based on adjacency to PAM sites and sequence mismatches inside and outside the sgRNA seed region (defined as the 3'-most 13 bp of the operator sequence) (*29*). We used default scoring and *S. pyogenes* Cas9 settings and specified that no sgRNA operators should be permitted in the genome ('accept' parameter = 0). Given these parameters, CasValue identified sgRNAs with seed sequences that matched the genome exactly or contained one mismatch and were adjacent to a primary or secondary PAM site. Any identified sgRNAs were discarded along with other sgRNAs and promoters sharing a pre-sequence. To use CasValue, we first compiled the *E. coli* genome into Bowtie index files via the `bowtie-build` command from Bowtie version 1.0.1 (*30*). Ultimately, we designed 3,000 sgRNA:promoter pairs derived from $P_R$ and 3,000 derived from apFAB126 meeting these criteria.

Next, we used a graph theoretic approach to select two subsets of 10 dissimilar sgRNA:promoter pairs (one set derived from $P_R$ and the other from apFAB126). To achieve this, we enriched the 3,000 pre-sequences encoded in the 6,000 designed promoters for mismatches. We represented sets of pre-sequences as complete graphs wherein vertices represented pre-sequences and edges were weighted by number of matching nucleotides occurring along two different pre-sequences. Framed this way, we sought a 10-vertex subgraph that minimized the sum of all edge weights. The global optimum for this problem cannot be found quickly, so we developed several heuristics-based approaches to expedite our search. To begin, we sorted the ~4.5 million edges of our 3,000-vertex pre-sequence graph and then collected 100 vertices starting from edges with the lowest weights (fewest matches). Then, we iteratively reduced the total number of pairwise nucleotide matches (i.e. sum of all edge weights) among the 100 selected pre-sequences by looping four times over the 2,900 excluded pre-sequences, incorporating one pre-sequence at a time (resulting in a 101-vertex graph), and then discarding the pre-sequence contributing the most matches to the total. The 100-pre-sequence subset resulting from this procedure contained 20,096 total matches with 4.1 matches per edge on average and maximum 13 matches (of 15 possible). Next, we reduced this set to multiple 10-pre-sequence subsets by starting with all sets of two pre-sequences containing 5 or fewer sequence matches (i.e. edges with weight $\leq 5$) and then randomly selecting a set to expand. Sets were expanded by looping over all excluded pre-sequences, incorporating one pre-sequence at a time, and retaining sets that maintained 5 or fewer matches between all pairs of sequences. 8,154 sets of size 10 or greater were generated using this approach (with the largest set containing 14 pre-sequences). To select a final set, we developed a simple linear scoring algorithm that punished sequence matches at the 3'-end of the pre-sequence. We then selected the set with the lowest average score per edge. From this set, we created 10 sgRNA:promoter pairs based on $P_R$ and 10 based on apFAB126.

The strength of all twenty corresponding promoters was then characterized using sfGFP and flow cytometry. We discovered that most of the $P_R$-derived promoters were significantly weaker than $P_R$. We therefore selected the apFAB126-derived promoters for all but one pre-sequence (corresponding to P6), where the $P_R$-derived promoter was closer in strength to the other nine in the set than the apFAB126-derived alternative. We observed that one of the 10 sgRNAs slowed growth upon induction (guide sequence = `TCGACACAACGTGACGTATG`), so we discarded it to arrive at nine orthogonal sgRNA:promoter pairs.

Promoter insulation

To avoid potential interference from upstream DNA sequences (*31*), we added insulator sequences to P1-P9, $P_{tet}$ (output promoter from the aTc sensor), $P_{tac}$ (output promoter from the IPTG sensor), and J23114 and J23115 (which constitutively drive transcription of *phlF* in the DAPG sensor and *luxR* in the AHL sensor, respectively). To design insulators, we generated random 70 bp sequences and screened them against a list of forbidden *E. coli* sigma factor binding sites, RBS features, transposon insertion sites, common restriction enzyme recognition sites, and unwieldy sequence repeats (*32*) using custom Python scripts. We also forbade the TetR O2 operator sequence, a sequence encoding the LacI operator site and a small portion of the LacI promoter, the native *S. pyogenes* Cas9 handle sequence, and the sequence of a structurally optimized handle (the "F+E" handle from (*33*)). Insulators were then paired with P1-P9 using promoter prediction tools (*34, 35*) to ensure additional unwanted promoters were not introduced. The $P_{tet}$, $P_{tac}$, J23114, and J23115 insulators were later chosen randomly from the remaining available insulator sequences. If necessary, insulators were truncated to achieve a total promoter length of 105 bp.

Strains, media, antibiotics, and chemical inducers

*E. coli* NEB 10-*β* was used for cloning and MG1655 was used for experiments. LB Miller medium (MilliporeSigma 1.10285.5000 or Fisher BioReagents BP9723-5) was used to maintain cloning strains, and M9 medium (M9 salts (6.78 g/L $Na_2HPO_4$, 3 g/L $KH_2PO_4$, 0.5 g/L NaCl, 1 g/L $NH_4Cl$ via BD 248510 or 5.962 g/L $Na_2HPO_4$, 3.266 g/L $KH_2PO_4$, 0.526 g/L NaCl, 1.016 g/L $NH_4Cl$ via Teknova M1902), 0.1 mM calcium chloride (Fisher Chemical C79-500), 2 mM magnesium sulfate (VWR BDH9246-500G), 0.4% glucose (Avantor Performance Materials 4908-06), and 0.2% casamino acids (MilliporeSigma 2240-500GM)) buffered to pH 6.6 with 100 mM HEPES (MilliporeSigma 5320-500GM) was used for all experiments but the orthogonality assay, where unbuffered M9 lacking HEPES was used. Plasmids with a ~15-copy ColE1, ~5-copy p15A, or ~3-copy pSC101* origin of replication (*36*) were used. ColE1 plasmids bear a chloramphenicol resistance marker and were maintained with 35 µg/mL chloramphenicol (Alfa Aesar B20841). p15A plasmids bear a spectinomycin antibiotic resistance marker and were maintained with 100 µg/mL spectinomycin (Gold Biotechnology S-140-25). pSC101* plasmids bear an ampicillin resistance marker and were maintained with 50 µg/mL ampicillin (Dot Scientific DS102040). aTc (Clontech 631310), IPTG (IBI Scientific IB02125), DAPG (Santa Cruz Biotechnology 2161-86-6), and AHL (Sigma-Aldrich, K3007-10MG) were used for chemical induction. aTc was dissolved in ethanol, IPTG was dissolved in filtered water and then filter-sterilized with a 0.22 µm syringe filter, DAPG was dissolved in 200 proof ethanol and then filter-sterilized with a 0.22 µm syringe filter, and AHL was dissolved in filtered water.

Plasmid assembly

Plasmids were constructed using Golden Gate assembly (*37*). DNA fragments were amplified using primers (Integrated DNA Technologies, Coralville, IA) and size-separated via gel electrophoresis. Correctly-sized bands were purified (Promega Wizard Gel Clean-Up kit, A9282) and assembled into bacterial plasmids via digestion-ligation reactions with BsaI and T4 DNA ligase (New England Biolabs; BsaI=R3535 or R3733, T4=M0202). All sequences were designed to lack BsaI sites. Hierarchical assembly, where smaller parts were assembled into larger linear intermediates, amplified, and purified for use in a subsequent round of assembly, was frequently used. The regions of plasmids containing engineered DNA fragments were verified via Sanger sequencing (GeneWiz or Lone Star Labs).

*cas9* was amplified from pMJ806 (*38*). *dcas9* was created by introducing the nuclease-deactivating mutations D10A and H840A (*23*) during plasmid construction. *dcas9* was expressed weakly from pSC31_1, which was used for the orthogonality assay, and strongly from pSC31_3, which was used for all other assays (**Fig. S15**). The aTc and IPTG sensors were sourced from pSR11-2 (*39*) (**Figs. S5, S16**). The DAPG sensor is based on a previously published version (*40*) (**Fig. S5, S16**). Terminator sequences within the IPTG and DAPG sensors were replaced with strong synthetic versions (*41*). Initially, the DAPG sensor output was low in absence and presence of DAPG, so we reduced the strength of the *phlF* promoter, which recovered DAPG-dependent activation of the $P_{PhlF}$ output promoter (**Table S12**). *luxI* and *luxR* were amplified from pJT105 (*6*). NOT and NOR gates were designed from sgRNA:promoter pairs and strong, sequence-distinct terminators (*41*) (**Figs. 2D and S4A**; terminators omitted for clarity). Circuit sequences were designed by fusing gate transcription units. Promoter prediction tools (*34*, *35*) were used to eliminate unwanted promoters from circuit sequences. Putatively inert DNA mimicking sgRNAs in length (102 bp) were used as outputs of DEMUX circuits and created by randomly joining and truncating two 70 bp insulator sequences. Gate transcription units were synthesized as gBlocks (IDT) and incorporated into TOPO TA plasmids (Life Technologies, 450030) for subsequent assembly steps. All plasmids used in this study are listed in **Table S16**.

Freezer aliquots and experimental pre-cultures

All experimental cultures were started from -80℃ freezer aliquots to decrease day-to-day variability (*44*). To prepare the aliquots, transformed strains were grown to exponential phase in LB, diluted 100-fold into buffered M9, and grown for an additional ~4 h to $OD_{600}$ ~0.1. Cultures were then placed on ice, and glycerol was added to 18% final concentration. The solution was gently mixed, and 110 µL was aliquoted into PCR tubes, which were stored at −80°C. To initiate an experiment, a freezer aliquot was removed from −80°C storage, thawed, and gently mixed before transferring a calculated volume ranging from <1 µL to 110 µL into 3 mL fresh buffered M9. This inoculated pre-culture was then grown shaking at 37°C to $OD_{600}$ ~0.1 and then used to inoculate new experiment cultures.

Orthogonality assay

S1-S9 were expressed under control of the aTc sensor on separate ColE1 plasmids. P1-P9:*sfgfp* fusions were encoded on p15A plasmids. Pairwise combinations of these plasmids were co-transformed with pSC31_1 to create 81 strains (**Table S17**). To begin the assay, 3 mL LB cultures containing appropriate antibiotics were inoculated from LB-glycerol freezer stocks. Cultures were then incubated at 37°C shaking overnight. The following morning, 14 mL culture tubes were prepared with 3 mL unbuffered M9, appropriate antibiotics, and 0 or 20 ng/mL aTc. Overnight cultures were then diluted into this media to $OD_{600} = 5 \times 10^{-4}$. The inoculated M9 cultures were then placed in the 37°C shaking incubator and grown for ~6 h to $OD_{600}$ of 0.3-0.6. Cultures were then removed from the 37°C shaking incubator and placed on ice. Flow cytometry tubes were also prepared with 1mL of phosphate-buffered saline (PBS) and placed on ice. 15 µL of culture was then transferred to a cytometry tube for measurement.

Probe plasmids

Probe plasmids were designed for P1-P9, P6*, $P_{tet}$, $P_{tac}$, $P_{PhlF}$, $P_{lux}$, and $P_R$. A standardized, insulated *sfgfp* gene was developed by fusing the self-cleaving ribozyme insulator RiboJ (*42*) to a synthetic RBS and the *sfgfp* coding sequence. This gene was then fused to promoters of interest and carried on plasmids with a p15A origin of replication (**Fig. S15**). To probe a circuit promoter, we co-transformed the appropriate probe plasmid with a circuit plasmid and pSC31_3 (**Fig. S2**). We probed entire circuits by probing individual promoters in separate strains.

NOT gate transfer functions

*E. coli* were transformed with one of the nine aTc-NOT*i* plasmids, one input ($P_{tet}$) or output (P1-P9) probe plasmid, and pSC31_3. Pre-cultures were prepared from freezer aliquots and diluted to $OD_{600} = 8.84 \times 10^{-5}$ into fresh buffered M9 containing 0, 1, 1.5, 2, 3, 4, or 20 ng/mL aTc. These cultures were then placed in the 37°C shaking incubator and grown for 5.75-6.5 h prior to fluorophore maturation and flow cytometry. An autofluorescence control strain lacking *sfgfp* and positive control strains lacking aTc-NOT*i* plasmids were similarly characterized at 0 ng/mL aTc.

MUX and DEMUX characterization

Each MUX test circuit plasmid was separately co-transformed with P5, P3, P2, and P6 probe plasmids, resulting in 32 strains. Each DEMUX test circuit plasmid was co-transformed with P8, P7, P9, and P2 probe plasmids to create 16 strains. All 48 strains also contained pSC31_3. Pre-cultures were prepared from freezer aliquots and then diluted into fresh buffered M9 to OD600 = $8.84 \times 10^{-5}$ and grown for 5.75 h shaking at 37°C prior to fluorophore maturation and flow cytometry.

Sensor transfer functions

*E. coli* were transformed with one of four sensor circuit plasmids (**Figs. S5, S9, S16**), a probe plasmid containing $P_{tet}$, $P_{tac}$, $P_{PhlF}$, or $P_{lux}$, and pSC31_3. Pre-cultures were prepared from freezer aliquots and then diluted to $OD_{600} = 8.84 \times 10^{-5}$ into fresh buffered M9 containing inducer and grown shaking at 37°C for 5.75-6.5 h prior to fluorophore maturation and flow cytometry. aTc was added to 0, 0.5, 0.793, 1.26, 1.99, 3.16, 5.01, 7.95, 12.6, and 20 ng/mL, IPTG was induced to 0, 0.0090, 0.0216, 0.0520, 0.125, 0.3, 0.721, 1.73, 4.16, and 10 mM, DAPG was induced to 0, 2.33, 4.36, 8.16, 15.3, 28.6, 53.5, 100, 187., and 350 µM, and AHL was induced to 0, 0.1, 0.237, 0.562, 1.33, 3.16, 7.50, 17.8, 42.2, and 100 nM.

*luxI* and *luxR* expression optimization

Pre-cultures were prepared from freezer aliquots and then diluted into fresh buffered M9 to $OD_{600} = 0.001$ for sender cells and $OD_{600} = 8.84 \times 10^{-5}$ for receiver cells. This difference (amounting to ~3.5 doublings) accounts for the slower growth of sender cells due to *luxI* expression. Co-cultures crossing all four sender strains with all four receiver strains were then grown shaking at 37°C for 5.75 h prior to fluorophore maturation and flow cytometry. We found cells well separated by their FL3 (mCherry) fluorescence and used a threshold of 1,000 MECY to classify senders (above) and receivers (below).

SENSOR-MUX, SENSOR-MUX* and AHL-DEMUX characterization

The SENSOR-MUX and SENSOR-MUX* circuit plasmids were co-transformed with $P_{tet}$, $P_{tac}$, $P_{Phlf}$, P1, P2, P3, P4, P5, P6 or P6*, and P9 probe plasmids, resulting in ten SENSOR-MUX and ten SENSOR-MUX* characterization strains. The AHL-DEMUX circuit plasmid was co-transformed with $P_{lux}$, $P_{PhlF}$, P2, P3, P7, P8, and P9 probe plasmids, resulting in seven AHL-DEMUX characterization strains. All strains carried pSC31_3.

Pre-cultures were prepared from freezer aliquots and then diluted into fresh buffered M9 containing inducers to $OD_{600} = 5.52 \times 10^{-6}$ for SENSOR-MUX and SENSOR-MUX* cells and $OD_{600} = 8.84 \times 10^{-5}$ for AHL-DEMUX cells. SENSOR-MUX and SENSOR-MUX* cells were induced with all eight combinations of aTc (0 and 20 ng/mL), IPTG (0 and 0.3 mM), and DAPG (0 and 100 µM), and AHL-DEMUX cells were induced with all four combinations of AHL (0 and 3.5 nM) and DAPG (0 and 100 µM). Induced cultures were grown shaking at 37°C for 9 h (SENSOR-MUX and SENSOR-MUX*) or 5.75 h (AHL-DEMUX) prior to fluorophore maturation and flow cytometry. SENSOR-MUX and SENSOR-MUX* incubation times were extended to accommodate circuit computation time. Prior experiments had also revealed that aTc was unable to induce cells for long periods of time, so we added a second dose (60 ng into 3 mL of culture) to +aTc SENSOR-MUX and SENSOR-MUX* cultures 5.5-6 h into their 9 h incubation.

AHL-DEMUX response to higher AHL concentrations (**Fig. S12**) was characterized using a similar protocol, wherein AHL-DEMUX strains probing $P_{lux}$, P7, and P2 were inoculated to $OD_{600} = 9.77 \times 10^{-5}$, induced with 0, 5, 10, and 50 nM AHL and 0 and 100 µM DAPG, and grown for 8 h. The 3.5 nM AHL data from the previous experiment was included in the visualization.

Co-culture experiments

We used the P3 probe plasmid to characterize SENSOR-MUX*-AHL output ($OUT_{MUX}$), and the P7 and P2 probe plasmids to characterize the two AHL-DEMUX outputs ($OUT_1$ and $OUT_2$) (**Fig. S11**). All three strains carried pSC31_3. To begin the experiment, a SENSOR-MUX*-AHL pre-culture was prepared from a freezer aliquot and then induced with aTc, IPTG, and DAPG. +IPTG cultures were diluted to $OD_{600} = 1.25 \times 10^{-6}$, and −IPTG cultures were diluted to $OD_{600} = 5 \times 10^{-6}$. These dilutions accounted for the increased duration of this experiment as well as a minor increase in cell growth rate we observed for +IPTG conditions. Induced SENSOR-MUX*-AHL cultures were then grown shaking at 37°C overnight for ~13 h. AHL-DEMUX pre-cultures were also prepared from freezer aliquots and grown overnight. Initial cell densities were designed to maintain cells in exponential phase overnight and to synchronize all cultures at sub-saturating densities ($OD_{600}$ <1.0) the following day. As before, we added a second dose of aTc (60 ng into 3mL of culture) to +aTc SENSOR-MUX*-AHL cultures ~8 h into the ~13 h incubation.

The next day, preconditioned SENSOR-MUX*-AHL cells and AHL-DEMUX pre-culture cells were diluted together into fresh buffered M9 and inducers. SENSOR-MUX*-AHL cells were diluted to $OD_{600} = 0.001$, and AHL-DEMUX cells were diluted to $OD_{600} = 8.84 \times 10^{-5}$. SENSOR-MUX*-AHL cells were diluted into cultures containing the same inducer combination. All eight combinations of aTc, IPTG, DAPG, and SENSOR-MUX*-AHL cells were prepared in duplicate, one for each AHL-DEMUX output strain. All 16 cocultures were then grown shaking at 37°C for 5.75 h prior to fluorophore maturation and flow cytometry. We found cells well separated by their FL3 (mCherry) fluorescence and used a threshold of 600 MECY to classify SENSOR-MUX*-AHL (below) and AHL-DEMUX (above) cells.

Dynamics experiments

$P_{PhlF}$, P4, P2, P6*, and P3 probe plasmids were used to characterize SENSOR-MUX*-AHL, and $P_{lux}$, P8, and P7 probe plasmids were used to characterize the AHL-DEMUX. All eight strains carried pSC31_3. Pre-cultures were prepared from freezer aliquots and then diluted into fresh buffered M9 to $OD_{600}$ = 6.6 x $10^{-6}$ and preconditioned overnight in 0.3 mM IPTG (SENSOR-MUX*-AHL) or 100 µM DAPG (AHL-DEMUX) for ~10 h. Initial cell densities were designed to maintain cells in exponential phase overnight. The next day, SENSOR-MUX*-AHL cultures were diluted 2:1 in fresh buffered M9 +IPTG to synchronize them with AHL-DEMUX cultures. All cultures were then diluted and additionally induced to 100 µM DAPG (SENSOR-MUX*-AHL) or 3.5 nM AHL (AHL-DEMUX). The original IPTG (SENSOR-MUX*-AHL) and DAPG (AHL-DEMUX) concentrations used to precondition cells were maintained. These temporal inducer profiles were designed to ensure circuits completed necessary pre-conditioning computations before toggling their respective portions of the longest CS computation path. After the second induction, cultures were sampled and diluted with fresh M9 and inducers every hour for 10 h. Induced M9 was pre-heated for 5-10 min. prior to dilution to avoid temperature-derived cell growth effects. Samples were also collected at half-hour time points for the first 3 h. 14 total time points were collected (T1-T14; T2, T4, and T6 being half-hour time points). All samples were immediately subjected to fluorophore maturation and then stored on ice until flow cytometry was performed.

Fluorophore maturation

Prior to flow cytometry, 30 or 50 µL of liquid culture was added to 500 µL PBS containing 1 mg/mL chloramphenicol, which inhibits protein synthesis in bacteria, and then incubated in a 37°C water bath for 1 h. This allows nascent, non-fluorescent sfGFP and mCherry molecules to mature and contribute to the cellular fluorescence signal, even during dynamics experiments (*45*, *44*). Samples were then placed on ice until flow cytometry was performed.

Flow cytometry

Cellular fluorescence was measured using a modified BD FACScan flow cytometer. Samples were collinearly excited by blue (488nm, 30mW) and yellow (561nm, 50mW) solid-state lasers (Cytek), and sample fluorescence was acquired through an optical bandpass filter centered at 510nm with a bandwidth of 21 nm (FL1) and separately through a 650 nm longpass filter (FL3). Forward scatter (FSC) and side scatter (SSC) fluorescence was also measured. Samples were acquired using either FlowJo Collectors' Edition 7.5.110.7 on a Windows 7 computer or BD CellQuest Pro 5.1.1 on a Macintosh computer. For *E. coli* samples, the voltage settings of the FSC, SSC, FL1, and FL3 detectors were FSC=10X, SSC=600, FL1=650, 750, or 850, and FL3=650, 750, or 850. FL1 and FL3 settings were adjusted to ensure fluorescence generated by cells did not saturate the detectors. An SSC threshold was specified in the acquisition software to eliminate low-SSC debris (below either 60% or 65% SSC). The fluorescence values of all events overcoming this threshold were recorded. Additionally, a polygon gate was manually drawn using the acquisition software to ensure enough *E. coli* cell-sized events were recorded. For monoculture experiments, 20,000-30,000 *E. coli* events were recorded. For coculture experiments, >10,000 events were recorded for both cell populations. For the orthogonality assay, 50,000-70,000 events were recorded.

SpheroTech rainbow calibration particles (RCP-30-5A) were used together with the FlowCal Python package (*43*) to calibrate FL1 and FL3 fluorescence measurements from arbitrary units (a.u.) to molecules of equivalent fluorescein (MEFL) and molecules of equivalent Cy5 (MECY), respectively. Calibrated fluorescence units enable comparison of measurements across instrument detector voltage and gain settings and account for long-term instrument drift (*43*). For calibration particle samples, the FSC and SSC detector voltage settings were 10X and 460, respectively. The SSC threshold setting was 10%. FL1 and FL3 detector voltage settings were specified to match the settings used to measure *E. coli* samples during the same flow cytometry session. One or two drops of calibration particles were aliquoted from the manufacturer-issued dropper bottle into 500 µL or 1 mL phosphate-buffered saline (PBS), which was then gently mixed and stored on ice until acquisition. Between 20,000 and 30,000 calibration particles were always acquired immediately prior to acquiring *E. coli* samples. After acquisition, FlowCal was used within custom Python scripts to generate MEFL and MECY calibration curves from calibration particle data. The calibration curves were then used to calibrate each FL1 and FL3 fluorescence measurement for all *E. coli* samples. FlowCal was also used to isolate *E. coli* cells from other particles. Specifically, a density gate was applied to all samples to retain events in the densest region of their FSC vs. SSC plots. The fraction of events retained was 0.85. Calibration particle samples were additionally gated to discard the first 250 and last 100 events and any events saturating the FSC or SSC channels, per default FlowCal behavior. Data from the orthogonality assay, which was taken prior to a major cytometer reconfiguration, was also gated to remove saturating events occurring in the lowest bin of the FL1 channel.

Visualization of cytometry data

We visualize population-level cellular fluorescence using frequency distributions and violin plots. To create a frequency distribution, the fluorescence measurements of a sample, which initially span 1,024 possible MEFL values, are first down-sampled into 64 bins logarithmically spanning the same range of values. This assigns 1 bin per 16 adjacent MEFL values. The frequency of each bin is then plotted and normalized such that the most frequent bin (i.e. the peak) spans half the y-axis of a frequency distribution plot. To create a violin plot, cellular fluorescence is first binned into 100 bins logarithmically spaced from 10 MEFL to 100,000 MEFL. The frequency of each bin is then displayed vertically using a normalized symmetrical histogram. The width of the histogram indicates the relative frequency of values at the associated y-axis position. The top and bottom 1% of measurements were trimmed (discarded) for aesthetic purposes. The center vertical axis of each violin indicates its associated x-axis value. Mean cellular fluorescence, calculated prior to trimming, is displayed as a solid line atop each violin.

Error calculations

To compare circuit behavior to model fits or simulations, we calculated the RMSE between predicted and measured $\log_{10}$-transformed sfGFP fluorescence values:

$$\mathrm{RMSE} = \sqrt{\frac{\sum_{i=1}^{N}\left[\left(\log_{10}\left(F_i^{\mathrm{pred.}}\right) - \log_{10}\left(F_i^{\mathrm{meas.}}\right)\right)^2\right]}{N}} \tag{1}$$

where $F$ is sfGFP fluorescence and $N$ is the total number of measurements. RMSE summarizes the difference between predicted and measured sfGFP fluorescence in $\log_{10}$ space in units of MEFL decades. If a sfGFP fluorescence value was equal to zero (e.g. for some predicted sensor outputs), that comparison was omitted from the RMSE calculation.

CS scaling laws

Laws describing how different features of a CS scale with the number of conversations it can transmit were determined (**Fig. S14**). Features include number of gates required, number of gate (i.e.

promoter) layers through which an input signal would have to propagate to reach its output, and maximum number of incoming (fan-in) and outgoing (fan-out) gate connections. The CS was assumed to be constructed from NOT and NOR gates, and its features were calculated assuming the MUX and DEMUX composing it were implemented using the same approach (i.e. 2-layer or Recursive). Laws were deduced by comparing different 2-, 4-, and 8-channel CS implementations. *c* is number of channels and $s$ is number of bits required for the SELECT signal, which is $\lceil \log_2 c \rceil$. For gates:

$$\text{gates}_{\text{MUX}}^{\text{2-layer}}(c) = c + s + 1 \tag{2}$$
$$\text{gates}_{\text{DEMUX}}^{\text{2-layer}}(c) = c + s + 1 \tag{3}$$
$$\text{gates}_{\text{CS}}^{\text{2-layer}}(c) = c + s + 1 \tag{4}$$
$$\text{gates}_{\text{MUX}}^{\text{Recursive}}(c) = 4(c - 1) \tag{5}$$
$$\text{gates}_{\text{DEMUX}}^{\text{Recursive}}(c) = 4(c - 1) \tag{6}$$
$$\text{gates}_{\text{CS}}^{\text{Recursive}}(c) = 4(c - 1) \tag{7}$$

For layers:

$$\text{layers}_{\text{MUX}}^{\text{2-layer}}(c) = 3 \tag{8}$$
$$\text{layers}_{\text{DEMUX}}^{\text{2-layer}}(c) = 2 \tag{9}$$
$$\text{layers}_{\text{CS}}^{\text{2-layer}}(c) = 6 \tag{10}$$
$$\text{layers}_{\text{MUX}}^{\text{Recursive}}(c) = 3s \tag{11}$$
$$\text{layers}_{\text{DEMUX}}^{\text{Recursive}}(c) = 2s \tag{12}$$
$$\text{layers}_{\text{CS}}^{\text{Recursive}}(c) = 5s + 1 \tag{13}$$

For maximum gate fan-in:

$$\text{max fan-in}_{\text{MUX}}^{\text{2-layer}}(c) = c \tag{14}$$
$$\text{max fan-in}_{\text{DEMUX}}^{\text{2-layer}}(c) = s + 1 \tag{15}$$
$$\text{max fan-in}_{\text{CS}}^{\text{2-layer}}(c) = c \tag{16}$$
$$\text{max fan-in}_{\text{MUX}}^{\text{Recursive}}(c) = 2 \tag{17}$$
$$\text{max fan-in}_{\text{DEMUX}}^{\text{Recursive}}(c) = 2 \tag{18}$$
$$\text{max fan-in}_{\text{CS}}^{\text{Recursive}}(c) = 2 \tag{19}$$

For maximum gate fan-out:

$$\text{max fan-out}_{\text{MUX}}^{\text{2-layer}}(c) = 2^{s-1} \tag{20}$$
$$\text{max fan-out}_{\text{DEMUX}}^{\text{2-layer}}(c) = c \tag{21}$$
$$\text{max fan-out}_{\text{CS}}^{\text{2-layer}}(c) = c \tag{22}$$
$$\text{max fan-out}_{\text{MUX}}^{\text{Recursive}}(c) = 1 \tag{23}$$
$$\text{max fan-out}_{\text{DEMUX}}^{\text{Recursive}}(c) = 2 \tag{24}$$
$$\text{max fan-out}_{\text{CS}}^{\text{Recursive}}(c) = 2 \tag{25}$$

### Supplementary Text

Relating transcriptional signals to sfGFP fluorescence

Throughout this study, we used probe plasmids and sfGFP fluorescence to report transcriptional signals of promoters of interest. To describe this relationship, we considered the following probe plasmid model:

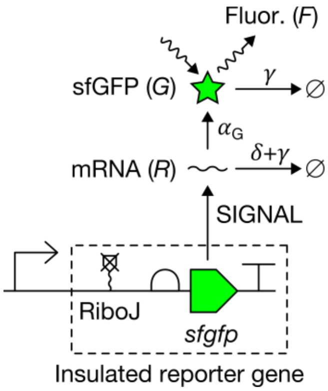

where SIGNAL is the promoter transcription rate (mRNA time$^{-1}$), $\delta$ is mRNA degradation rate (time$^{-1}$), $\alpha_G$ is sfGFP translation rate (sfGFP mRNA$^{-1}$ time$^{-1}$), $\gamma$ is dilution rate due to cell growth (time$^{-1}$), and $F$ is measured sfGFP fluorescence (MEFL). We assume that RiboJ renders $\alpha_G$ independent of the promoter sequence by removing any promoter-specific portion of the mRNA (*42*). Our fluorophore maturation protocol allows us to assume all sfGFP proteins mature instantaneously to the fluorescent state. sfGFP expression dynamics can therefore be described by the following ordinary differential equations:

$$\frac{dR}{dt} = \text{SIGNAL} - (\delta + \gamma)R \tag{26}$$

$$\frac{dG}{dt} = \alpha_G R - \gamma G \tag{27}$$

We also expect sfGFP fluorescence to be proportional to number of sfGFP molecules:

$$F = \eta G \tag{28}$$

where $\eta$ is fluorescence per molecule (MEFL sfGFP$^{-1}$). Steady-state sfGFP fluorescence ($F_{\text{SIGNAL}}^{SS}$) is therefore:

$$F_{\text{SIGNAL}}^{SS} = \text{SIGNAL}^{SS} \cdot \frac{\alpha_G}{\gamma^{SS}(\delta + \gamma^{SS})} \cdot \eta \tag{29}$$

$$= \text{SIGNAL}^{SS} \cdot \left(k_{\text{sfgfp}} \cdot \eta\right) \tag{30}$$

where $k_{\text{sfgfp}}$ is a promoter-independent steady-state expression constant (sfGFP (mRNA time$^{-1}$)$^{-1}$). In the text, the ($k_{\text{sfgfp}} \cdot \eta$) proportionality constant (MEFL (mRNA time$^{-1}$)$^{-1}$) is omitted for simplicity.

NOT gate models

We fit the following Hill model to the measured NOT1-NOT9 and NOT6* transfer functions:

$$\mathrm{OUT}_{\mathrm{NOT}i} = \mathrm{NOT}i(\mathrm{IN}_{\mathrm{NOT}i}) = \mathrm{GATE}i_{\min} + \frac{(\mathrm{P}i_{\max} - \mathrm{GATE}i_{\min})}{1 + \left(\frac{\mathrm{IN}_{\mathrm{NOT}i}}{K}\right)^{n}} \quad (31)$$

where $\mathrm{P}i_{\max}$ is mean sfGFP fluorescence produced by P$i$ in the absence of S$i$ (MEFL), $\mathrm{GATE}i_{\min}$ is minimum gate output (MEFL), $K$ is the $\mathrm{IN}_{\mathrm{NOT}i}$ value at which $\mathrm{OUT}_{\mathrm{NOT}i}$ is half repressed (MEFL), and $n$ is the Hill coefficient (dimensionless), which describes the steepness of the transfer function. Here, $\mathrm{IN}_{\mathrm{NOT}i}$ and $\mathrm{OUT}_{\mathrm{NOT}i}$ represent steady-state sfGFP fluorescence produced by the underlying $\mathrm{IN}_{\mathrm{NOT}i}$ and $\mathrm{OUT}_{\mathrm{NOT}i}$ transcriptional signals (i.e. $\mathrm{IN}_{\mathrm{NOT}i} = F^{SS}_{\mathrm{IN}_{\mathrm{NOT}i}}$ and $\mathrm{OUT}_{\mathrm{NOT}i} = F^{SS}_{\mathrm{OUT}_{\mathrm{NOT}i}}$). It can be shown, however, that $n$ is unaffected by the ($k_{\mathrm{sfgfp}} \cdot \eta$) proportionality constant and the underlying $\mathrm{GATE}i_{\min}$, $\mathrm{P}i_{\max}$, and $K$ parameters can be calculated by dividing by ($k_{\mathrm{sfgfp}} \cdot \eta$). Thus, our transfer functions still capture fundamental gate behavior and can be expected to accurately predict gate output given gate input.

To fit these models, we used the Lmfit Python package to perform a constrained least-squares fit of $\mathrm{GATE}i_{\min}$, $K$, and $n$ to pairs of $\mathrm{IN}_{\mathrm{NOT}i}$ and $\mathrm{OUT}_{\mathrm{NOT}i}$ fluorescence values using the default Levenberg-Marquardt fitting algorithm (*46*). $\mathrm{P}i_{\max}$ was fixed as mean sfGFP fluorescence produced by P$i$ in the absence of the gate sgRNA. $\mathrm{GATE}i_{\min}$ was constrained to [0,∞), $K$ was constrained to [1e−5,∞), and $n$ was constrained to [1e−3,∞). Initial parameter values were $\mathrm{GATE}i_{\min}$=50 MEFL, $K$=350 MEFL, and $n$=3.0. As with the RMSE calculation, the residual of each fit was calculated in $\log_{10}$ MEFL space (i.e. residual = $\log_{10}(\mathrm{OUT}^{\mathrm{measured}}_{\mathrm{NOT}i}) - \log_{10}(\mathrm{OUT}^{\mathrm{predicted}}_{\mathrm{NOT}i})$). Plots are shown in **Fig. 2E** and **Fig. S7B** and the resulting fit parameters are listed in **Table S1**. Two of three replicates for NOT2 measured at 0 ng/mL aTc yielded negative $\mathrm{IN}_{\mathrm{NOT}i}$ values and were discarded from the fit.

NOR gate models

We created NOR1-NOR9 and NOR6* models by extending each corresponding NOT model with a second transcriptional input term ($\mathrm{IN}_{\mathrm{NOR}i}2$):

$$\mathrm{OUT}_{\mathrm{NOR}i} = \mathrm{NOR}i(\mathrm{IN}_{\mathrm{NOR}i}1, \mathrm{IN}_{\mathrm{NOR}i}2) = \mathrm{GATE}i_{\min} + \frac{(\mathrm{P}i_{\max} - \mathrm{GATE}i_{\min})}{1 + \left(\frac{\mathrm{IN}_{\mathrm{NOR}i}1 + \mathrm{IN}_{\mathrm{NOR}i}2}{K}\right)^{n}} \quad (32)$$

where $\mathrm{IN}_{\mathrm{NOR}i}1$ and $\mathrm{IN}_{\mathrm{NOR}i}2$ are mean sfGFP fluorescence (MEFL) reporting two independent transcriptional inputs. Plots are shown in **Fig. S4** and fit parameters are listed in **Table S1**.

MUX model

MUX output is described by composing the models of the component gates:

$$\mathrm{OUT}_{\mathrm{MUX}} = \mathrm{NOR3}(\mathrm{NOR5}(\mathrm{IN}_1, \mathrm{SELECT}), \mathrm{NOR6}(\mathrm{NOT2}(\mathrm{SELECT}), \mathrm{IN}_2)) \quad (33)$$

$\mathrm{IN}_1$, $\mathrm{IN}_2$, and SELECT were modeled as {0, $\mathrm{P1}_{\max}$}, {0, $\mathrm{P9}_{\max}$}, and {0, $\mathrm{P4}_{\max}$} MEFL, respectively, where $\mathrm{P}i_{\max}$ is maximum mean sfGFP fluorescence produced by P$i$. $\mathrm{P}i_{\max}$ values are listed in **Table S1**. Total cellular fluorescence was simulated by summing sfGFP fluorescence with mean autofluorescence (145 MEFL). Simulations are listed in **Table S2**, and cellular fluorescence simulations are plotted in **Fig. 3A**.

DEMUX model

DEMUX output is similarly described by:

$$\mathrm{OUT_{DEMUX}1 = NOR7(NOT8(IN_{DEMUX}), SELECT)} \tag{34}$$
$$\mathrm{OUT_{DEMUX}2 = NOR2\big(NOT8(IN_{DEMUX}), NOT9(SELECT)\big)} \tag{35}$$

$IN_{DEMUX}$ and SELECT were modeled as {0, $P_{R,max}$} and {0, $P3_{max}$} MEFL, respectively. $P3_{max}$ is listed in **Table S1**, and $P_{R,max}$ was measured to be 5864 MEFL using the same protocol used to measure other $Pi_{max}$ values (**Methods**). Total cellular fluorescence was simulated as with the MUX model. Simulations are listed in **Table S3**, and cellular fluorescence simulations are plotted in **Fig. 3B**.

SENSOR-MUX model

The SENSOR-MUX model was constructed from MUX, NOT1, NOT9, NOT4, and all-or-none aTc, IPTG, and DAPG sensor models. Its output is described by:

$$\mathrm{OUT_{SENSOR\text{-}MUX} = OUT_{MUX}(IN_1, IN_2, SELECT)} \tag{36}$$

where

$$\mathrm{IN_1 = NOT1(OUT_{aTc\ sensor})} \tag{37}$$
$$\mathrm{IN_2 = NOT9(OUT_{IPTG\ sensor})} \tag{38}$$
$$\mathrm{SELECT = NOT4(OUT_{DAPG\ sensor})} \tag{39}$$

$OUT_{aTc\ sensor}$, $OUT_{IPTG\ sensor}$, and $OUT_{DAPG\ sensor}$ were modeled as {0, $P_{tet,induced}$}, {0, $P_{tac,induced}$}, and {0, $P_{PhlF,induced}$} MEFL, respectively. $P_{tet,induced}$, $P_{tac,induced}$, and $P_{PhlF,induced}$ are mean sfGFP fluorescence (MEFL) produced by the aTc, IPTG, and DAPG sensors upon induction with 20 ng/mL aTc, 0.3 mM IPTG, and 100 µM DAPG, respectively, (**Fig. S5**) and are 1146 MEFL, 8176 MEFL, and 4932 MEFL, respectively. Total cellular fluorescence was simulated as before. Simulations are listed in **Table S5**, and cellular fluorescence simulations are plotted in **Fig. S6**.

SENSOR-MUX* model

The SENSOR-MUX* model was constructed by replacing NOR6 with NOR6* in the SENSOR-MUX model. Its output is described by:

$$\mathrm{OUT_{SENSOR\text{-}MUX^*} = OUT_{MUX^*}(IN_1, IN_2, SELECT)} \tag{40}$$

where

$$\mathrm{OUT_{MUX^*} = NOR3(NOR5(IN_1, SELECT), NOR6^*(NOT2(SELECT), IN_2))} \tag{41}$$

and $IN_1$, $IN_2$, and SELECT are as described for the SENSOR-MUX model. The DAPG sensor model was modified in response to SENSOR-MUX characterization to incorporate leakiness observed in the absence of IPTG:

$$\mathrm{OUT_{DAPG\ sensor}} = \begin{cases} \mathrm{P_{PhlF,\ leaky}}, & \mathrm{DAPG{=}0, IPTG{=}0} \\ 0, & \mathrm{DAPG{=}0, IPTG{=}1} \\ \mathrm{P_{PhlF,\ induced}}, & \mathrm{DAPG{=}1} \end{cases} \tag{42}$$

$P_{PhlF,leaky}$ was calculated by averaging the mean sfGFP fluorescence values from the two relevant DAPG sensor output measurements in the SENSOR-MUX (DAPG=0, IPTG=0; **Table S5**) and is 1316 MEFL. The aTc and IPTG sensors were modeled as they were in the SENSOR-MUX. Total cellular fluorescence was simulated as before. Simulations are listed in **Table S6**, and cellular fluorescence simulations are plotted in **Fig. S8**.

AHL-DEMUX model

The AHL-DEMUX model was constructed from DEMUX, NOT3, and all-or-none AHL and DAPG sensor models. Its output is described by:

$$\mathrm{OUT_{AHL\text{-}DEMUX}1} = \mathrm{OUT_{DEMUX}1(IN_{DEMUX}, SELECT)} \tag{43}$$
$$\mathrm{OUT_{AHL\text{-}DEMUX}2} = \mathrm{OUT_{DEMUX}2(IN_{DEMUX}, SELECT)} \tag{44}$$

where

$$\mathrm{IN_{DEMUX}} = \mathrm{OUT_{AHL\,sensor}} \tag{45}$$
$$\mathrm{SELECT} = \mathrm{NOT3(OUT_{DAPG\,sensor})} \tag{46}$$

Thus,

$$\mathrm{OUT_{AHL\text{-}DEMUX}1} = \mathrm{NOR7}\big(\mathrm{NOT8(OUT_{AHL\,sensor}), NOT3(OUT_{DAPG\,sensor})}\big) \tag{47}$$
$$\mathrm{OUT_{AHL\text{-}DEMUX}2} = \mathrm{NOR2}\Big(\mathrm{NOT8(OUT_{AHL\,sensor}), NOT9}\big(\mathrm{NOT3(OUT_{DAPG\,sensor})}\big)\Big) \tag{48}$$

$OUT_{AHL\ sensor}$ and $OUT_{DAPG\ sensor}$ were modeled as $\{0, P_{lux,induced}\}$ and $\{0, P_{PhlF,induced}\}$ MEFL, respectively. $P_{PhlF,induced}$ was as specified previously, and $P_{lux,induced}$ was calculated from the AHL sensor transfer function (**Fig. S9D, Table S4**) with an input of 3.5 nM and is 1041 MEFL. Total cellular fluorescence was simulated as before. Simulations are listed in **Table S7**, and cellular fluorescence simulations are plotted in **Fig. S10**.

Dynamical SENSOR-MUX*-AHL and AHL-DEMUX gene expression models

To model the gene expression dynamics of SENSOR-MUX*-AHL and AHL-DEMUX, we first devised dynamical gate models based on our steady-state models:

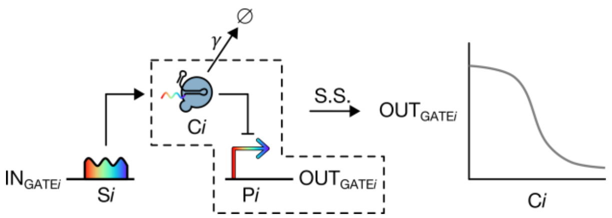

Each dynamical model describes how the gate dCas9:sgRNA complex (C*i*) changes over time:

$$\frac{dCi}{dt} = \mathrm{IN}_{\mathrm{GATE}i} - \gamma Ci \tag{49}$$

Here, $IN_{GATEi}$ is total transcriptional input to the gate (mRNA time$^{-1}$), which is $IN_{NOTi}$ for a NOT gate and $IN_{NORi}1$+$IN_{NORi}2$ for a NOR gate, $OUT_{GATEi}$ is transcriptional output from the gate (mRNA time$^{-1}$), and γ is dilution rate of C*i* due to cell growth (time$^{-1}$). We equated C*i* production rate to $IN_{GATEi}$ based on the following assumptions: (1) minimal sgRNA degradation occurs, due either to quick uptake and protection by dCas9 or low activate mRNA degradation, (2) dCas9:sgRNA complex formation is fast relative to sgRNA synthesis, and (3) dCas9 is in excess. We also assumed C*i* is stable and relies on dilution by cell growth for elimination.

To relate $\text{OUT}_{\text{GATE}i}$ to C*i*, we assumed C*i* rapidly equilibrates with P*i* (instantaneously achieving steady state), which allowed us to invoke our steady-state characterizations. We began by considering our gate transfer functions (generalized here via $\text{IN}_{\text{GATE}i}$):

$$\text{OUT}_{\text{GATE}i} = \text{NOT}i(\text{IN}_{\text{GATE}i}) \tag{50}$$

We then recalled that these transfer functions describe fluorescence values, not transcription rates (see "NOT models" section above). To recognize this, we reintroduced ($k_{\text{sfgfp}} \cdot \eta$) and then isolated $\text{OUT}_{\text{GATE}i}$:

$$\text{OUT}_{\text{GATE}i} = \frac{1}{k_{\text{sfgfp}} \cdot \eta} \cdot \text{NOT}i\left(\text{IN}_{\text{GATE}i} \cdot k_{\text{sfgfp}} \cdot \eta\right) \tag{51}$$

Next, we assumed that C*i* was related to $\text{IN}_{\text{GATE}i}$ at steady state by some invertible function *f*:

$$\text{C}i = f(\text{IN}_{\text{GATE}i}) \tag{52}$$

In considering different mechanistic forms of *f*, we found that *f* consistently factored into the following form when dCas9 was in excess:

$$f_{\text{excess dCas9}}(\text{IN}_{\text{GATE}i}) = \text{IN}_{\text{GATE}i} \cdot k_{\text{C}i} \tag{53}$$

where $k_{\text{C}i}$ is a steady-state expression constant (C*i* (mRNA time$^{-1}$)$^{-1}$), much like $k_{\text{sfgfp}}$. Indeed, for the simple model proposed above, $k_{\text{C}i} = \frac{1}{\gamma^{SS}}$. More generally, though, this assumption allowed us to relate $\text{OUT}_{\text{GATE}i}$ to C*i* via *f*:

$$\text{OUT}_{\text{GATE}i} = \frac{1}{k_{\text{sfgfp}} \cdot \eta} \cdot \text{NOT}i\left(f^{-1}(C\text{i}) \cdot k_{\text{sfgfp}} \cdot \eta\right) \tag{54}$$

For practical reasons, we chose not to model our gates in this form, as doing so would require estimating several unknown parameters (e.g. $\alpha_{\text{G}}$ and η). Instead, we transformed C*i* and $\text{OUT}_{\text{GATE}i}$ into proxy sfGFP fluorescence signals and then modeled their dynamics. First, we transformed C*i* into the sfGFP fluorescence signal that would have accompanied it during the steady-state characterization (*Fi*, in MEFL):

$$Fi = f^{-1}(C\text{i}) \cdot \left(k_{\text{sfgfp}} \cdot \eta\right) \tag{55}$$

This transformation maps C*i* to $\text{IN}_{\text{NOT}i}^{SS}$ and then to *Fi*. Similarly, we transformed dynamical transcriptional signals ($\text{IN}_{\text{GATE}i}$ and $\text{OUT}_{\text{GATE}i}$) into the sfGFP fluorescence signals they would generate at steady-state ($F_{\text{IN}_{\text{GATE}i}}^{SS}$, $F_{\text{OUT}_{\text{GATE}i}}^{SS}$, in MEFL):

$$F^{SS}_{\mathrm{IN}_{\mathrm{GATE}i}} = \mathrm{IN}_{\mathrm{GATE}i} \cdot \left(k_{\mathrm{sfgfp}} \cdot \eta\right) \tag{56}$$

$$F^{SS}_{\mathrm{OUT}_{\mathrm{GATE}i}} = \mathrm{OUT}_{\mathrm{GATE}i} \cdot \left(k_{\mathrm{sfgfp}} \cdot \eta\right) \tag{57}$$

$$= \mathrm{NOT}i\left(f^{-1}(C\mathrm{i}) \cdot k_{\mathrm{sfgfp}} \cdot \eta\right) \tag{58}$$

$$= \mathrm{NOT}i(Fi) \tag{59}$$

Here, we noted that $F^{SS}_{\mathrm{OUT}_{\mathrm{GATE}i}}$ could now be easily calculated as NOT*i*(*Fi*). Next, we considered the dynamical behavior of *Fi*:

$$\frac{dFi}{dt} = \frac{d}{dt}\left[f^{-1}(C\mathrm{i}) \cdot \left(k_{\mathrm{sfgfp}} \cdot \eta\right)\right] \tag{60}$$

$$= \left(k_{\mathrm{sfgfp}} \cdot \eta\right) \cdot \frac{d}{dt}\left[f^{-1}(\mathrm{C}i)\right] \tag{61}$$

Here, we assumed $f^{-1}$ was differentiable (which, practically speaking, requires that $f$ is strictly increasing), applied the chain rule, incorporated the underlying dynamical behavior of C*i*, and related C*i* and $\mathrm{IN}_{\mathrm{GATE}i}$ to *Fi* and $F^{SS}_{\mathrm{IN}_{\mathrm{GATE}i}}$, respectively:

$$\frac{dFi}{dt} = \left(k_{\mathrm{sfgfp}} \cdot \eta\right) \cdot \left[\frac{df^{-1}}{d\mathrm{C}i} \cdot \frac{d\mathrm{C}i}{dt}\right] \tag{62}$$

$$= \left(k_{\mathrm{sfgfp}} \cdot \eta\right) \cdot \left[\frac{df^{-1}}{d\mathrm{C}i} \cdot \left[\mathrm{IN}_{\mathrm{GATE}i} - \gamma \mathrm{C}i\right]\right] \tag{63}$$

$$= \left(k_{\mathrm{sfgfp}} \cdot \eta\right) \cdot \left[\frac{df^{-1}}{d\mathrm{C}i} \cdot \left[\frac{F^{SS}_{\mathrm{IN}_{\mathrm{GATE}i}}}{k_{\mathrm{sfgfp}} \cdot \eta} - \gamma \cdot f\left(\frac{Fi}{k_{\mathrm{sfgfp}} \cdot \eta}\right)\right]\right] \tag{64}$$

To simplify this expression, we relied on the excess dCas9 assumption, which allowed us to populate $f$ as $f_{\mathrm{excess\ dCas9}}$. Interestingly, we found that the sfGFP proportionality constants were eliminated:

$$\frac{dFi}{dt} = \left(k_{\mathrm{sfgfp}} \cdot \eta\right) \cdot \left[\frac{1}{k_{\mathrm{C}i}} \cdot \left[\frac{F^{SS}_{\mathrm{IN}_{\mathrm{GATE}i}}}{k_{\mathrm{sfgfp}} \cdot \eta} - \gamma \cdot \left(\frac{Fi \cdot k_{\mathrm{C}i}}{k_{\mathrm{sfgfp}} \cdot \eta}\right)\right]\right] \tag{65}$$

$$= \frac{1}{k_{\mathrm{C}i}} F^{SS}_{\mathrm{IN}_{\mathrm{GATE}i}} - \gamma \cdot Fi \tag{66}$$

As an aside, we suspect that protein-based gates would abstractly conform to $f_{\mathrm{excess\ dCas9}}$ (i.e. the protein regulator would be proportional to $\mathrm{IN}_{\mathrm{GATE}i}$ at steady state) and could easily be substituted into this dynamical framework. Finally, we recognized that $k_{\mathrm{C}i} = \frac{1}{\gamma^{SS}}$ for the simple C*i* dynamics model proposed above (we also assumed $\gamma^{\mathrm{SS}} = \gamma$):

$$\frac{dFi}{dt} = \gamma\left(F^{SS}_{\mathrm{IN}_{\mathrm{GATE}i}} - Fi\right) \tag{67}$$

This final dynamics expression can alternatively be interpreted as driving $Fi$ to a set point ($F^{SS}_{\text{IN}_{\text{GATE}i}}$), which is often the output of a steady-state transfer function, with dynamics governed by γ. This interpretation comports with other models used previously (*45*, *39*, *22*). Together, $Fi$ and $F^{SS}_{\text{OUT}_{\text{GATE}i}}$ constitute a dynamical gate model. A time-varying $Fi(t)$ signal, which is a proxy for C$i$, can be calculated by numerically integrating $\frac{dFi}{dt}$, and a time-varying gate output signal ($F^{SS}_{\text{OUT}_{\text{GATE}i}}(t)$) can then be calculated as $\text{NOT}i\big(Fi(t)\big)$.

To realize circuit models from these gate models, we connected them together and with sensor output signals. To do so, we first collected all $Fi$ and $F^{SS}_{\text{OUT}_{\text{GATE}i}}$ terms into vectors:

$$\boldsymbol{c}(t) = [F1(t), \cdots, F9(t), F6^{*}(t)] \tag{68}$$

$$\boldsymbol{o}_{\textbf{GATES}}(t) = \left[F^{SS}_{\text{OUT}_{\text{GATE1}}}\big(F1(t)\big), \cdots, F^{SS}_{\text{OUT}_{\text{GATE9}}}\big(F9(t)\big), F^{SS}_{\text{OUT}_{\text{GATE6}^*}}\big(F6^{*}(t)\big)\right] \tag{69}$$

To incorporate sensor output signals, we augmented $\boldsymbol{o}_{\textbf{GATES}}$ with sensor transcription rates transformed to steady-state sfGFP fluorescence values, as done above:

$$\boldsymbol{o}(t) = \left[F^{SS}_{\text{OUT}_{\text{aTc sensor}}}(t), F^{SS}_{\text{OUT}_{\text{IPTG sensor}}}(t), F^{SS}_{\text{OUT}_{\text{AHL sensor}}}(t), F^{SS}_{\text{OUT}_{\text{DAPG sensor}}}(t) \mid \boldsymbol{o}_{\textbf{GATES}}(t)\right] \tag{70}$$

We then described the dynamical behavior of all $Fi$ using a circuit connectivity matrix **M**, which specifies which transcriptional signals transcribe which complexes:

$$\frac{d\boldsymbol{c}}{dt} = \boldsymbol{oM} - \gamma\boldsymbol{c} \tag{71}$$

Here, $m_{ij} \in \{0,1\}$ indicates whether signal $i$ transcribes C$j$, and the dot product of $\boldsymbol{o}$ and $\boldsymbol{M}$ sums all transcription rates (via their sfGFP proxy signals) for each C$i$ complex across the circuit specified by **M**.

Finally, we simulated the expression of sfGFP from every gate and sensor. Ideally, this would be done by decoding the transcription rate from $F^{SS}_{\text{OUT}_{\text{GATE}i}}$ or $F^{SS}_{\text{OUT}_{\text{SENSOR}}}$ and then using it to simulate a sfGFP expression model like the one proposed in the first section. However, this would again require estimating unknown parameters. Instead, we assumed a simplified sfGFP expression model, akin to that of C$i$:

$$\frac{dG_i}{dt} = \text{OUT}_i - \gamma G_i \tag{72}$$

where $i$ corresponds to the gate or sensor being probed. Then, using the same line of reasoning we used with C$i$, we transformed $G_i$ into $F_{Gi}$, the sfGFP fluorescence that $G_i$ would produce at steady state. Because we used the same insulated *sfgfp* gene and a fluorophore maturation protocol, $F_{Gi}$ should also match sfGFP fluorescence measured during our dynamics experiments. To model $F_{Gi}$ dynamics, we simply augmented **c** and the columns of **M** with $F_{Gi}$ terms. The final connectivity matrices for SENSOR-MUX*-AHL and AHL-DEMUX are shown in **Tables S13 and S14**.

To simulate SENSOR-MUX*-AHL and AHL-DEMUX, we specified sensor output signals and then numerically evaluated equation (71) parameterized with the SENSOR-MUX*-AHL or AHL-DEMUX connectivity matrix (**Tables S13 and S14**) using a custom Python script. Each circuit was simulated for 30 hours starting at time $t$ = −19 h with 1-minute time steps. Sensor output signals were designed to reflect induction by IPTG and DAPG at time $t$ = −10 h followed by induction with DAPG and

AHL at time $t$ = 0 h, respectively. The SENSOR-MUX*-AHL sensor output signals were therefore specified as follows:

$$F^{SS}_{\mathrm{OUT}_{\mathrm{aTc\ sensor}}}(t) = \quad 0, \quad -19.0\ \mathrm{h} \le t < 11.0\ \mathrm{h} \tag{73}$$

$$F^{SS}_{\mathrm{OUT}_{\mathrm{IPTG\ sensor}}}(t) = \begin{cases} 0, & -19.0\ \mathrm{h} \le t < -10.0\ \mathrm{h} \\ \mathrm{P}_{\mathrm{tac,\ induced}}, & -10.0\ \mathrm{h} \le t < 11.0\ \mathrm{h} \end{cases} \tag{74}$$

$$F^{SS}_{\mathrm{OUT}_{\mathrm{DAPG\ sensor}}}(t) = \begin{cases} 0, & -19.0\ \mathrm{h} \le t < 0.0\ \mathrm{h} \\ \mathrm{P}_{\mathrm{PhlF,\ induced}}, & 0.0\ \mathrm{h} \le t < 11.0\ \mathrm{h} \end{cases} \tag{75}$$

The AHL-DEMUX sensor output signals were specified as follows:

$$F^{SS}_{\mathrm{OUT}_{\mathrm{AHL\ sensor}}}(t) = \begin{cases} 0, & -19.0\ \mathrm{h} \le t < 0.0\ \mathrm{h} \\ \mathrm{P}_{\mathrm{lux,\ induced}}, & 0.0\ \mathrm{h} \le t < 11.0\ \mathrm{h} \end{cases} \tag{76}$$

$$F^{SS}_{\mathrm{OUT}_{\mathrm{DAPG\ sensor}}}(t) = \begin{cases} 0, & -19.0\ \mathrm{h} \le t < -10.0\ \mathrm{h} \\ \mathrm{P}_{\mathrm{PhlF,\ induced}}, & -10.0\ \mathrm{h} \le t < 11.0\ \mathrm{h} \end{cases} \tag{77}$$

$P_{tac,induced}$, $P_{lux,induced}$, and $P_{PhlF,induced}$ are as defined for the SENSOR-MUX and AHL-DEMUX steady-state models. C$i$ and $G_i$ were initialized to 0 (i.e. **c**($t$ = −19 h) = 0). γ was calculated from $OD_{600}$ measurements taken during the dynamics experiment and averaged across strains and was 0.8776 hour$^{-1}$ (doubling time = 47.4 min.) with a standard deviation of 0.017 hour$^{-1}$. Measured and simulated sfGFP fluorescence (i.e. $F_{Gi}$) for the SENSOR-MUX*-AHL and AHL-DEMUX dynamics experiments are listed in **Tables S10 and S11** and plotted for 0 ≤ $t$ ≤ 10.5 h in **Fig. S13**.

### Supplementary Figures

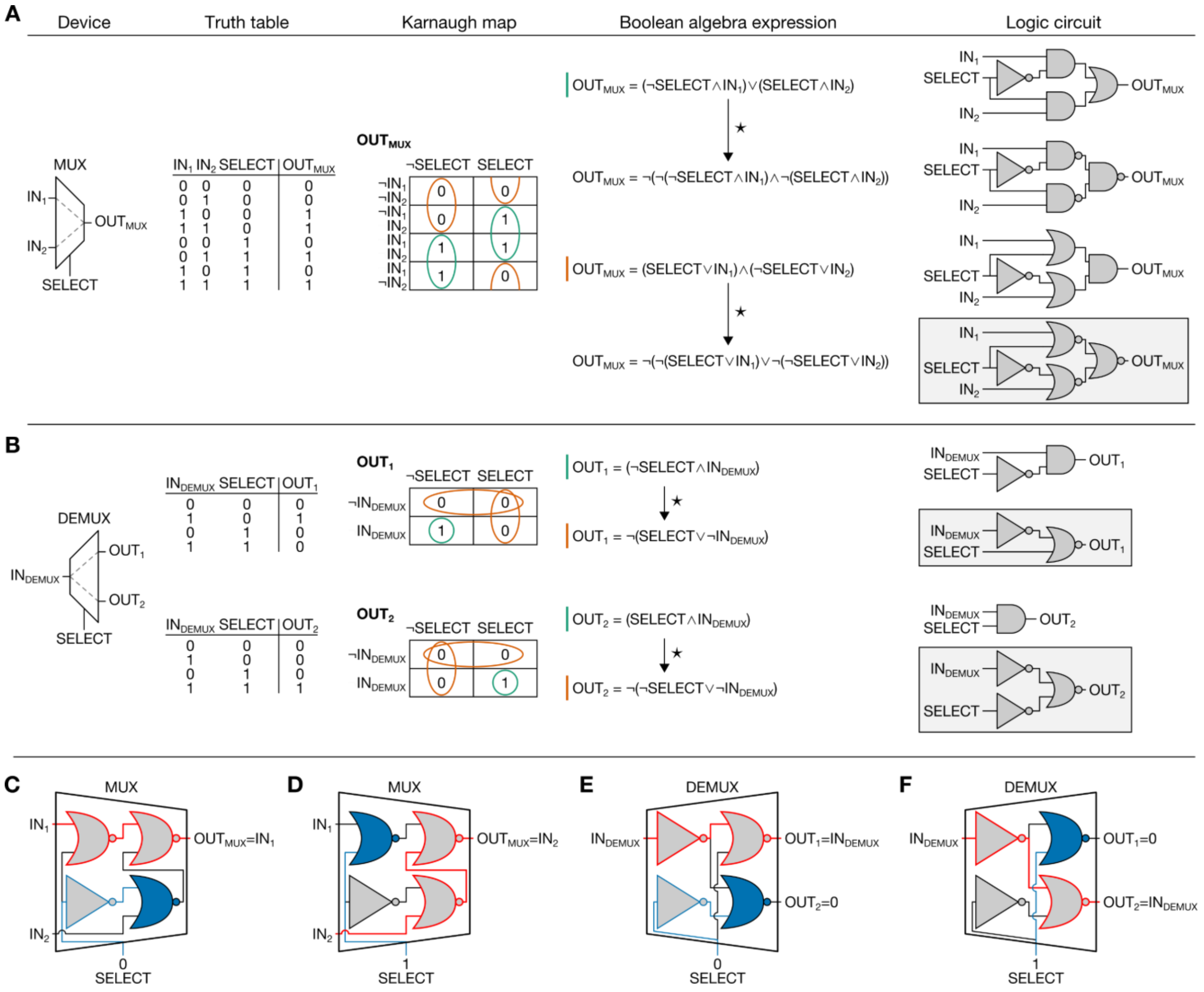

**Fig. S1. Design of MUX and DEMUX circuits.** Logic synthesis was used to design a (**A**) 2-input/1-output MUX and (**B**) 1-input/2-output DEMUX. First, the function of each circuit was described via a truth table, where all possible digital input combinations are listed alongside expected outputs. Karnaugh maps, which are graphical representations of truth tables organized to identify logic patterns, were then used to derive simplified Boolean algebra expressions for $OUT_{MUX}$, $OUT_1$, and $OUT_2$. Multiple synonymous minimized logic circuits were then created by reading each Boolean algebra expression. We selected circuits that only use NOR and NOT gates. The $OUT_1$ and $OUT_2$ circuits were combined to implement the DEMUX. ¬ specifies logical negation (NOT), ∧ specifies logical conjunction (AND), ∨ specifies logical disjunction (OR), and ⋆ specifies algebraic manipulation via De Morgan's law. (**C** - **F**) Gate-level illustrations of how the MUX and the DEMUX select one input for propagation to the appropriate output. If either input to a NOR gate is a logical 1, the output will be a logical 0 independent of the other input. The MUX and DEMUX SELECT signals use this mechanism to halt the propagation of $IN_2$ (**C**) or $IN_1$ (**D**) in the MUX and to deactivate $OUT_2$ (**E**) or $OUT_1$ (**F**) in the DEMUX. Solid blue gates have an input of 1 originating from SELECT (path highlighted in blue), and red highlights the path of the propagated input signal.

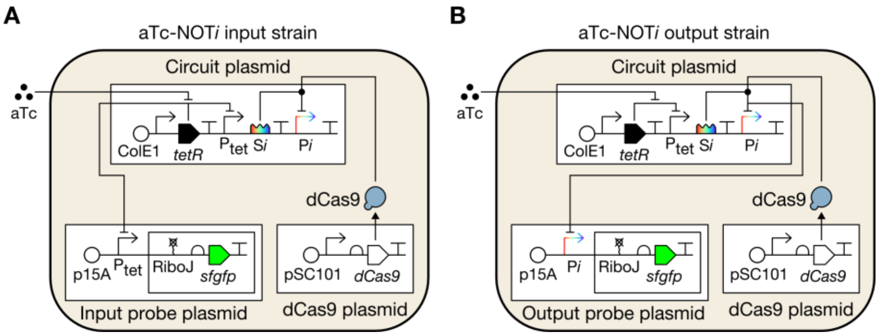

**Fig. S2. Probing NOT gate inputs and outputs.** Schematics of aTc-NOT*i* (**A**) input and (**B**) output probe strains.

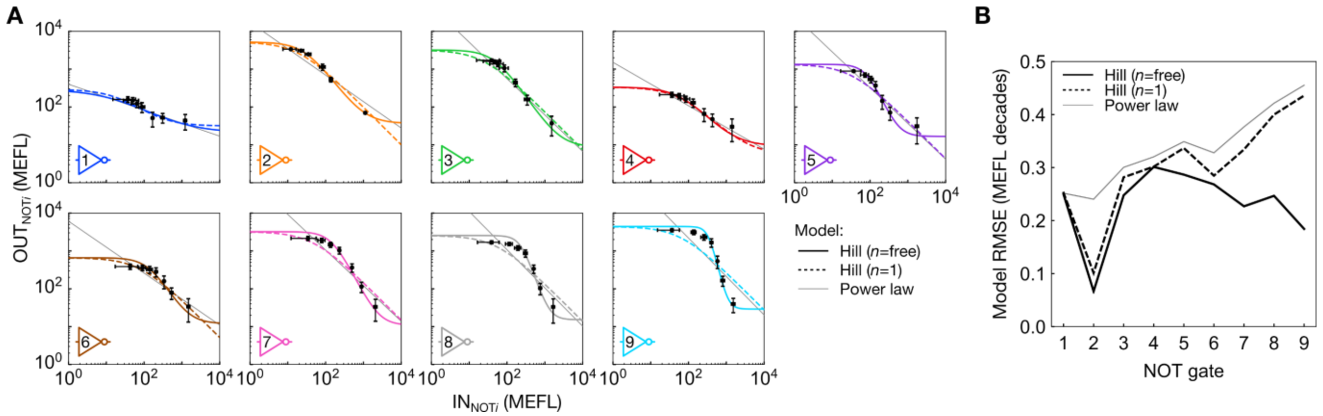

**Fig. S3. Comparisons of NOT gate models.** (**A**) Unconstrained Hill (*n* = free), constrained Hill (*n* = 1), and power law models fit to each gate transfer function. Fit parameters are listed in **Table S1**. (**B**) Performance of each model for each gate. The constrained and unconstrained Hill models perform similarly for NOT1-6. The unconstrained model performs better for NOT7-9 as their transfer functions are more sigmoidal. The power law models perform similarly to the constrained Hill models.

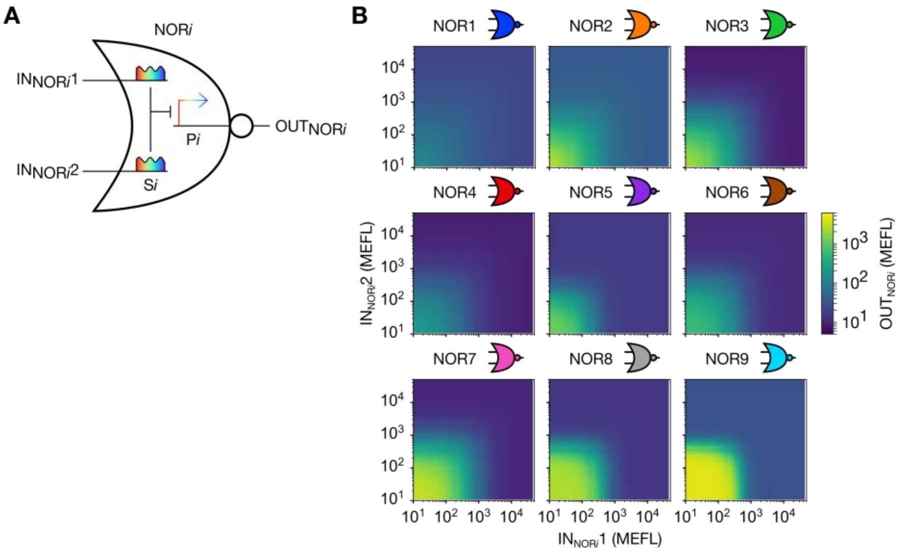

**Fig. S4. NOR gate models.** (**A**) General schematic for a CRISPRi-based NOR gate. (**B**) Transfer function model simulations for NOR1-NOR9.

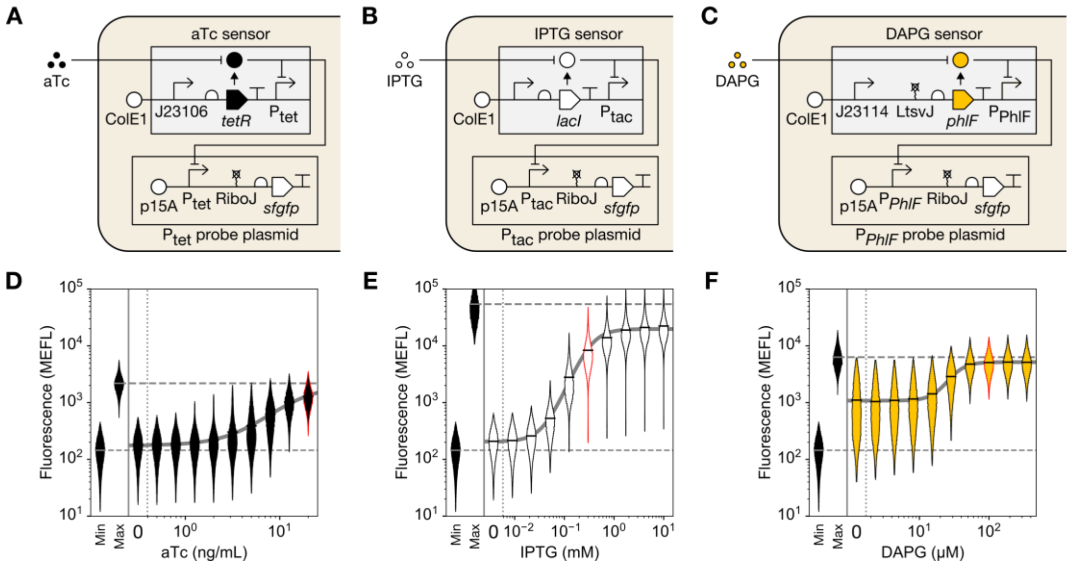

**Fig. S5. Sensor transfer functions.** (**A-C**) Schematics of the strains used to probe the transfer functions. pSC31_3 is present in all strains but not pictured. (**D-F**) Sensor transfer functions. Violin plots, which contain vertical histograms centered on corresponding x-axis values, are used to show mean and variation in sensor output over a range of input ligand concentrations. Min shows cellular autofluorescence and max shows sensor output when the repressor is absent (aTc and DAPG sensors) or expressed from the genome only (IPTG sensor). Solid gray lines show activating Hill model fits to mean sfGFP fluorescence (**Table S4**), which was summed with mean autofluorescence for these plots. The ligand concentrations chosen to activate each sensor in all main text experiments is highlighted in red (aTc = 20 ng/mL, IPTG = 0.3 mM, DAPG = 100 μM). The DAPG sensor suffers from leaky transcription in ~60% of cells at low DAPG concentrations (see protracted violins in panel F). We also witnessed this phenomenon in circuits if IPTG was absent (**Figs. S6 and S8**). All violins represent data combined from experiments on three separate days, except for the max violins, where only one replicate was measured.

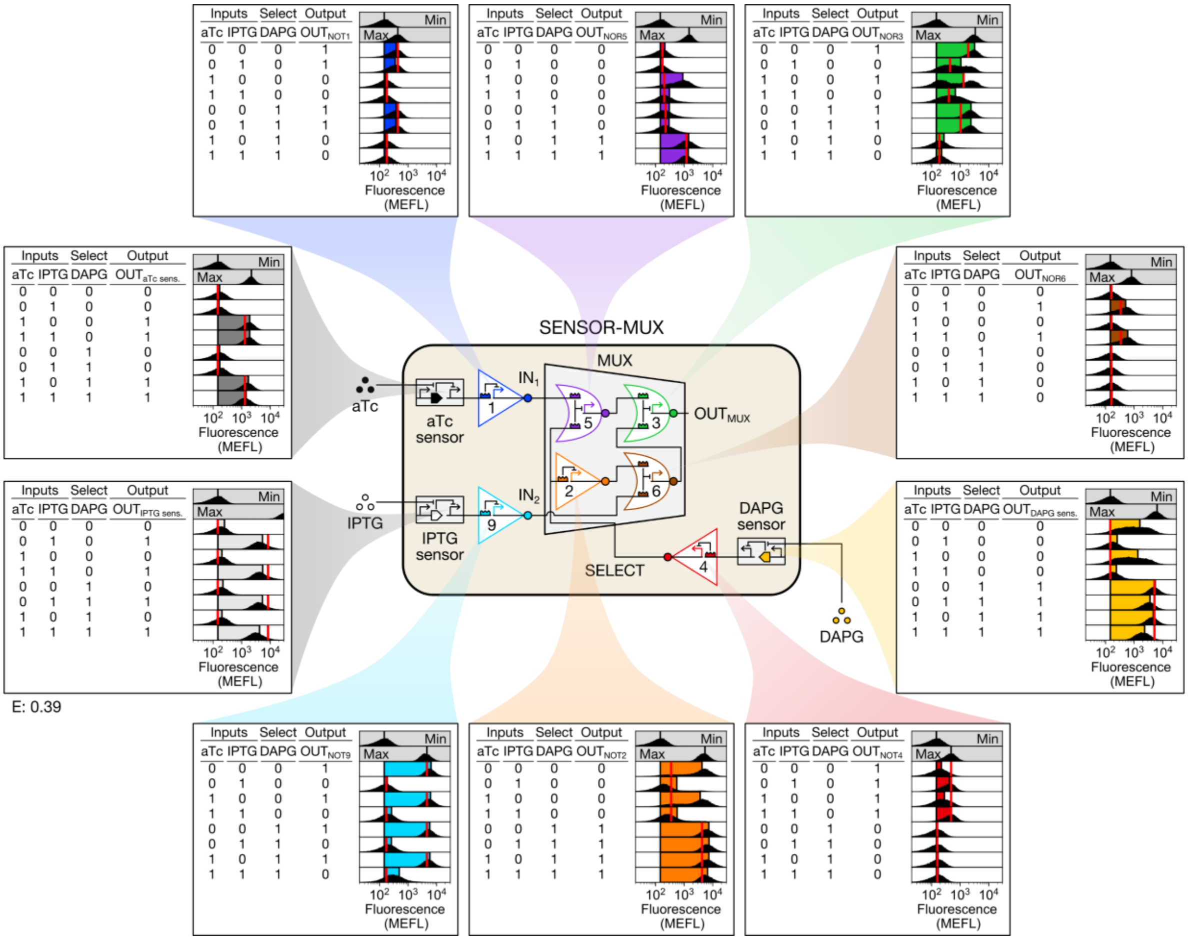

**Fig. S6. SENSOR-MUX characterization.** The SENSOR-MUX was constructed from the MUX by expressing S1, S9, and S4 from the aTc, IPTG, and DAPG sensors, respectively, therein placing $IN_1$, $IN_2$, and SELECT under the control of aTc, IPTG, and DAPG. The SENSOR-MUX circuit plasmid was then separately co-transformed with the $P_{tet}$, $P_{tac}$, $P_{PhlF}$, P1, P2, P3, P4, P5, P6, and P9 probe plasmids, and the resulting ten strains were incubated with the eight possible binary combinations of aTc, IPTG, and DAPG (**Methods**). Min was measured in triplicate on three separate days, max was measured once on a fourth day, and all other measurements were performed on a fifth day. For sensors, max shows sensor output when the repressor is absent (aTc and DAPG sensors) or expressed from the genome only (IPTG sensor). Inducer concentrations: 0 (0); 20 ng/mL aTc, 0.3 mM IPTG, and 100 μM DAPG (1).

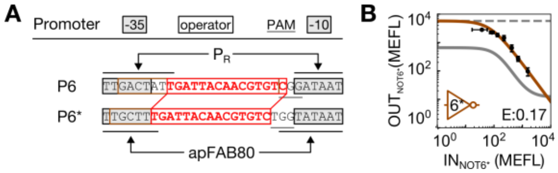

**Fig. S7. Design and characterization of NOT6*.** (**A**) The NOT6 output promoter P6 was originally designed by adding a PAM site to a randomly generated pre-sequence (red) and then flanking that sequence with -35 and -10 sequences from $P_R$. To design P6*, we moved the P6 pre-sequence into the strong promoter apFAB80 (*28*) and insulated it with the original P6 insulator sequence (not shown). The NOT6* gate was then created by designing a new cognate sgRNA (S6*). (**B**) NOT6* transfer function. NOT6* was characterized exactly as NOT1-9 were, and data are presented as presented in **Fig. 2E**. The solid gray line shows the original NOT6 transfer function for reference. NOT6* achieves almost 10-fold higher outputs relative to NOT6 but can still be repressed to low (< 100 MEFL) outputs, suggesting it should appropriately repress NOR3 in the SENSOR-MUX.

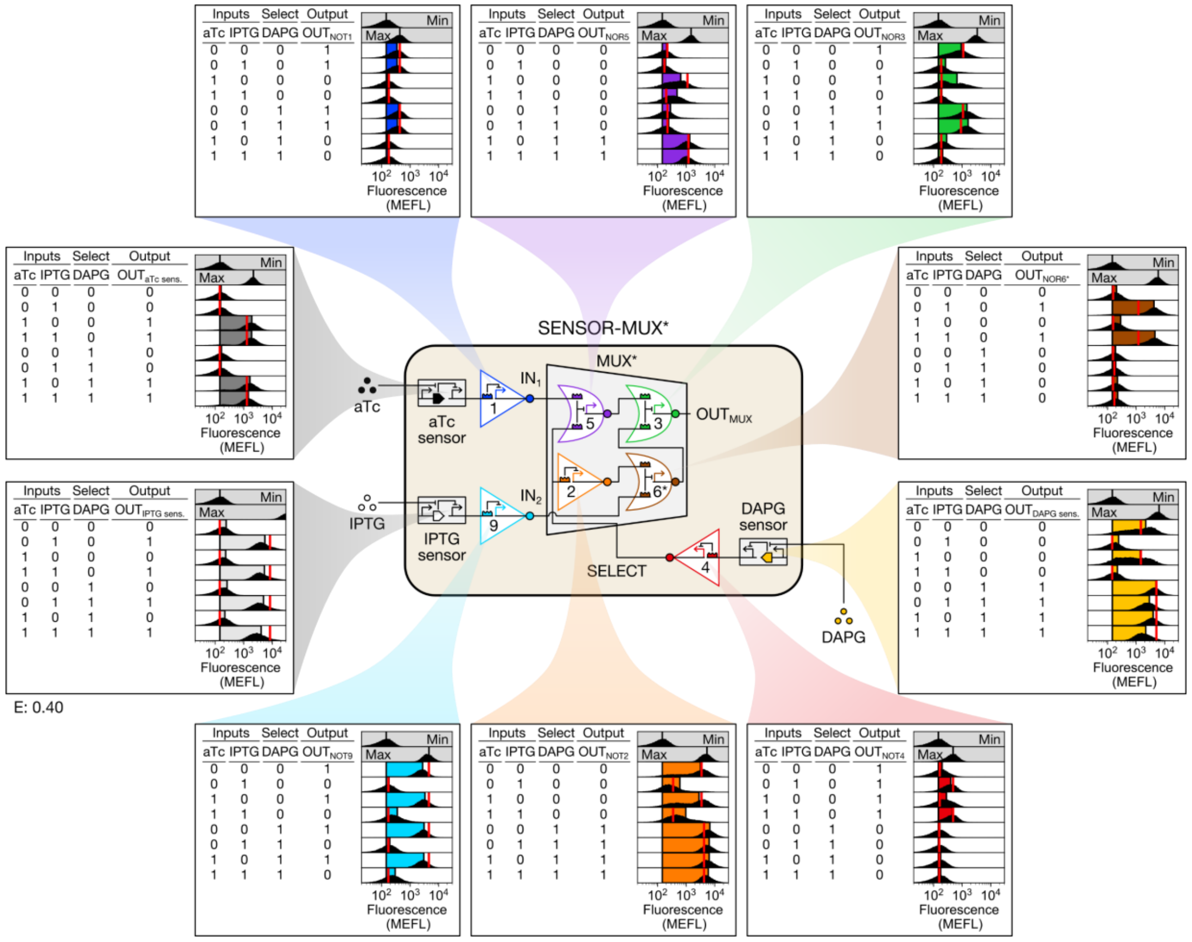

**Fig. S8. SENSOR-MUX* characterization.** SENSOR-MUX* was constructed from the SENSOR-MUX by replacing the weaker NOR6 with the stronger NOR6*. The SENSOR-MUX* circuit plasmid pJS0301 was then separately co-transformed with the $P_{tet}$, $P_{tac}$, $P_{PhlF}$, P1, P2, P3, P4, P5, P6*, and P9 probe plasmids, and the resulting ten strains were incubated with the eight possible binary combinations of aTc, IPTG, and DAPG. Min was measured in triplicate on three separate days, max was measured once on a fourth day, and all other measurements were performed on a fifth day. Inducer concentrations: 0 (0); 20ng/mL aTc, 0.3mM IPTG, and 100µM DAPG (1).

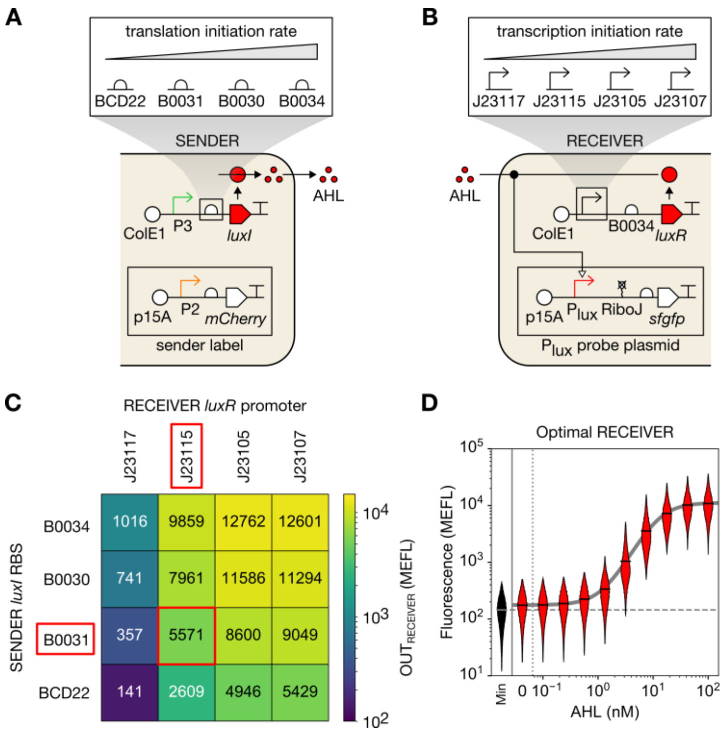

**Fig. S9. Optimization of the cell-cell communication system.** (**A**-**B**) Schematics of strains used to optimize the cell-cell communication system. Four SENDER strains were created that use different ribosome binding sites (RBSs) to initiate translation of the *luxI* AHL-biosynthetic gene, and four RECEIVER strains were created that use different constitutive promoters to express the *luxR* transcription factor gene. The SENSOR-MUX* output promoter P3 drives *luxI* to simulate its use in our CS, where it will transmit the output of SENSOR-MUX* to the DEMUX. RECEIVERs probe the AHL-responsive $P_{lux}$ promoter to report communication, and SENDERs contain a constitutive mCherry expression plasmid for identification by flow cytometry. pSC31_3 is present in all strains but not pictured. (**C**) Heatmap showing mean sfGFP fluorescence generated by $P_{lux}$ after co-culturing all combinations of SENDERs and RECEIVERs. In our CS, $P_{lux}$ will receive the AHL-mediated SENSOR-MUX* output signal in the DEMUX. As such, we selected a *luxI*-*luxR* expression combination that achieved $P_{lux}$ expression most closely matching that of $P_R$ (5,864 MEFL), which was previously used to drive $IN_{DEMUX}$ (**Fig. 3B**). This occurred with the B0031 *luxI* RBS and the BBa_J23115 *luxR* promoter (boxed in red). (**D**) Transfer function of the optimal RECEIVER strain. A violin plot shows cellular fluorescence as a function of exogenous AHL. All violins represent data combined from experiments on three separate days.

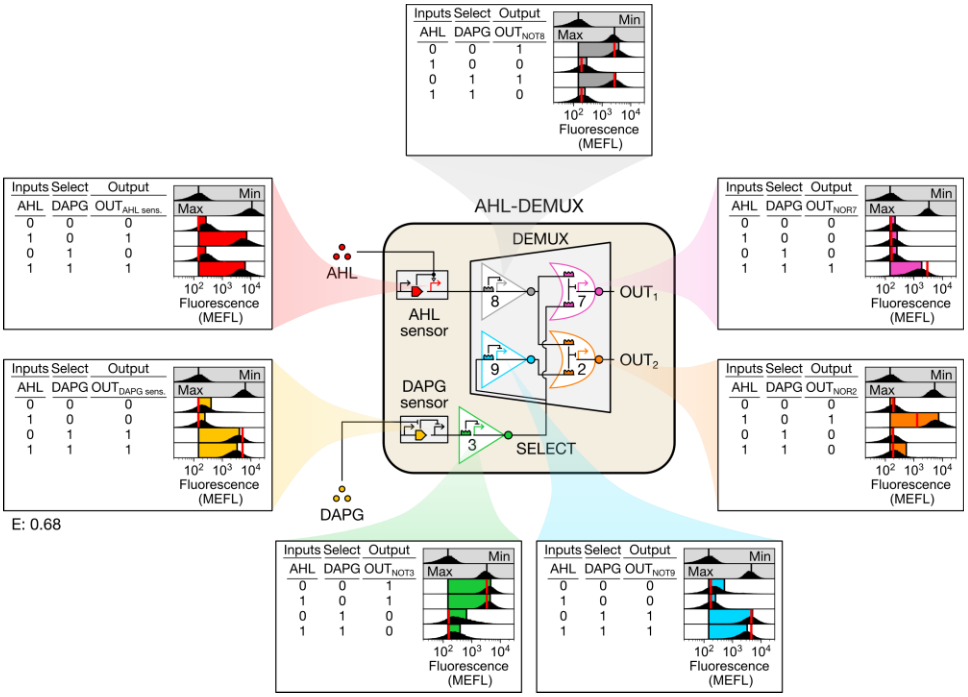

**Fig. S10. AHL-DEMUX characterization.** The AHL-DEMUX was constructed from the DEMUX by replacing the $P_R$ input promoter with the AHL sensor and expressing S3 from the DAPG sensor, therein placing $IN_{DEMUX}$ and SELECT under the control of AHL and DAPG, respectively. A constitutive mCherry expression cassette was also added (not pictured) to fluorescently distinguish AHL-DEMUX cells. As with the DEMUX, the AHL-DEMUX simulation error is noticeably larger than that of the MUX, SENSOR-MUX, and SENSOR-MUX*. This arises due to prediction errors near autofluorescence, which occur more frequently in the DEMUX circuits and appear less significant when autofluorescence is included, as above. Min was measured in triplicate on three separate days, max was measured once on a fourth day, and all other measurements were performed on a fifth day. For sensors, Max shows sensor output when the repressor is absent (DAPG sensors) or at maximum induction (AHL sensor). Inducer concentrations: 0 (0); 3.5nM AHL, 100µM DAPG (1).

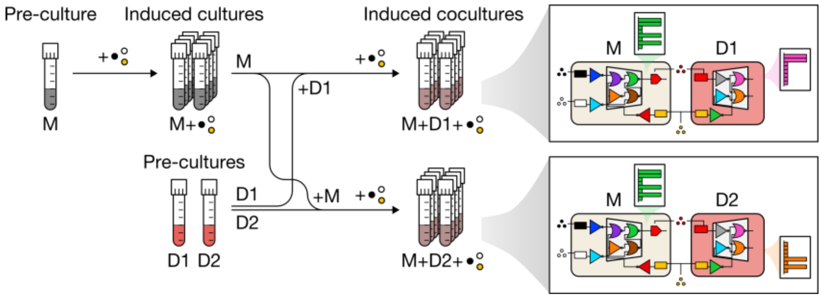

**Fig. S11. Design of the CS co-culture experiment.** SENSOR-MUX*-AHL cells (M) were first grown in an un-induced pre-culture and then diluted into fresh cultures containing the eight combinations of aTc, IPTG, and DAPG (inducers are shown here as colored circles). Two AHL-DEMUX strains probing $OUT_1$ and $OUT_2$, respectively, (D1, D2) were simultaneously grown in pre-cultures while SENSOR-MUX*-AHL cells were induced. Then, induced SENSOR-MUX*-AHL cells were diluted with one of the two AHL-DEMUX strains and fresh inducers to inoculate co-cultures. Co-cultures were grown and then cellular fluorescence was measured via flow cytometry to quantify $OUT_{MUX}$, $OUT_1$, and $OUT_2$ signals. A constitutively expressed mCherry fluorescent reporter was expressed in the AHL-DEMUX strains and used to unmix SENSOR-MUX*-AHL and AHL-DEMUX fluorescence signals. See **Methods** for full details.

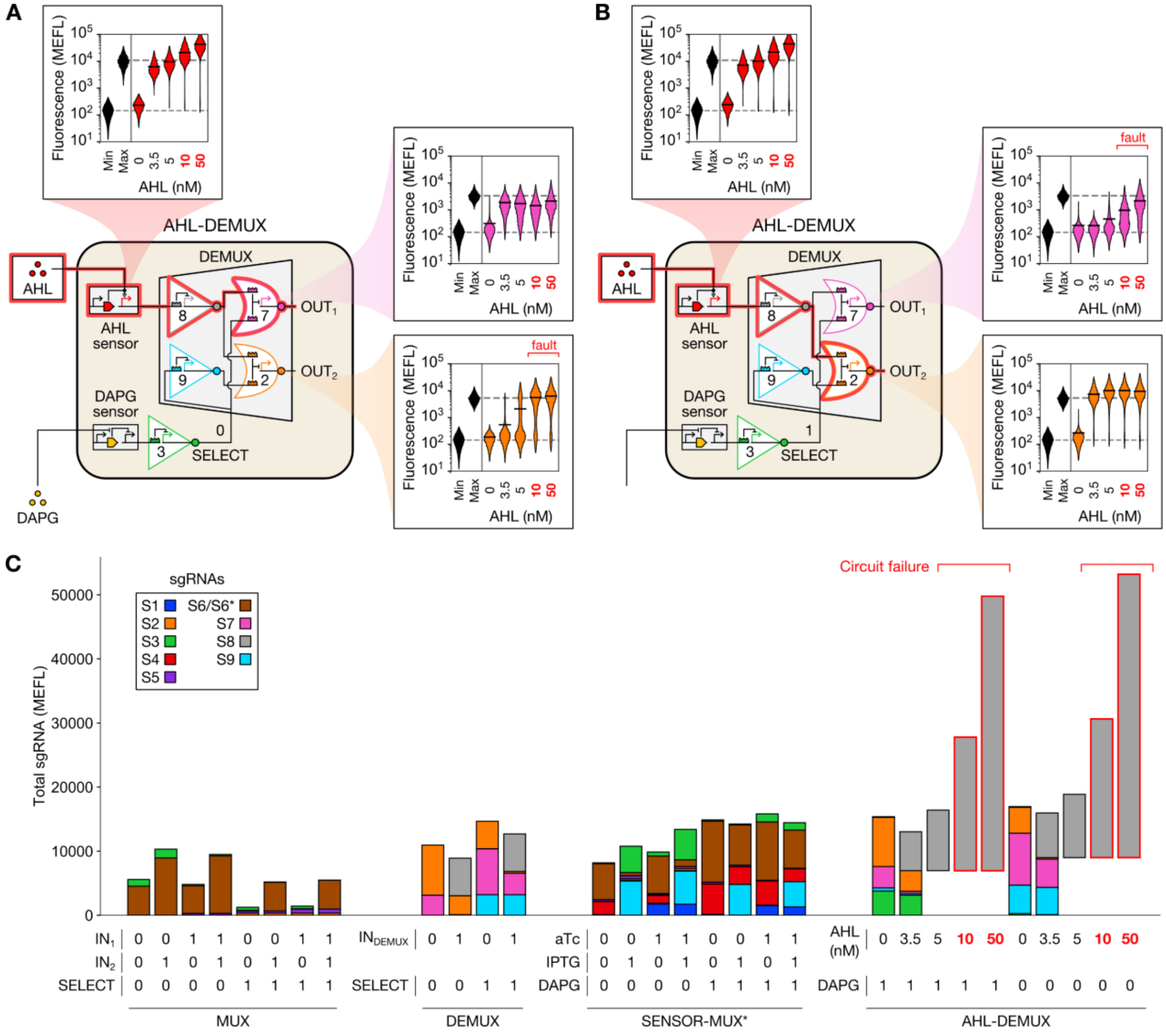

**Fig. S12. Characterization of the AHL-DEMUX at high AHL concentrations.** (**A**, **B**) Measurements of $OUT_{AHL\ sensor}$, $OUT_1$, and $OUT_2$ for AHL-DEMUX cells incubated at various AHL concentrations. At concentrations >5 nM, the output not selected ($OUT_2$ when SELECT is low and $OUT_1$ when SELECT is high) fails high in a sub-population of cells, and the relative size of that sub-population increases as more AHL is introduced. By contrast, correct outputs are achieved at 0 - 5 nM AHL. Max indicates gate output in the absence of sgRNA (NOR7 and NOR2) or maximum $P_{lux}$ activation measured during the communication channel optimization experiments (**Fig. S9D**). All violins represent data from one experiment, except for min violins, where data from three experiments performed on separate days was combined. (**C**) Total sgRNA produced by circuits characterized in this study. sgRNA was calculated by summing the probed mean sfGFP fluorescence (in MEFL) produced by each promoter expressing a given sgRNA (**Table S9**). Circuits behaved correctly if less than ~20,000 MEFL sgRNA was expressed. At higher total expression levels, though, circuit failures arose, as described in (A, B). The AHL-DEMUX sgRNA expression profile for 5, 10, and 50 nM AHL was assumed to match the corresponding profile at 3.5 nM.

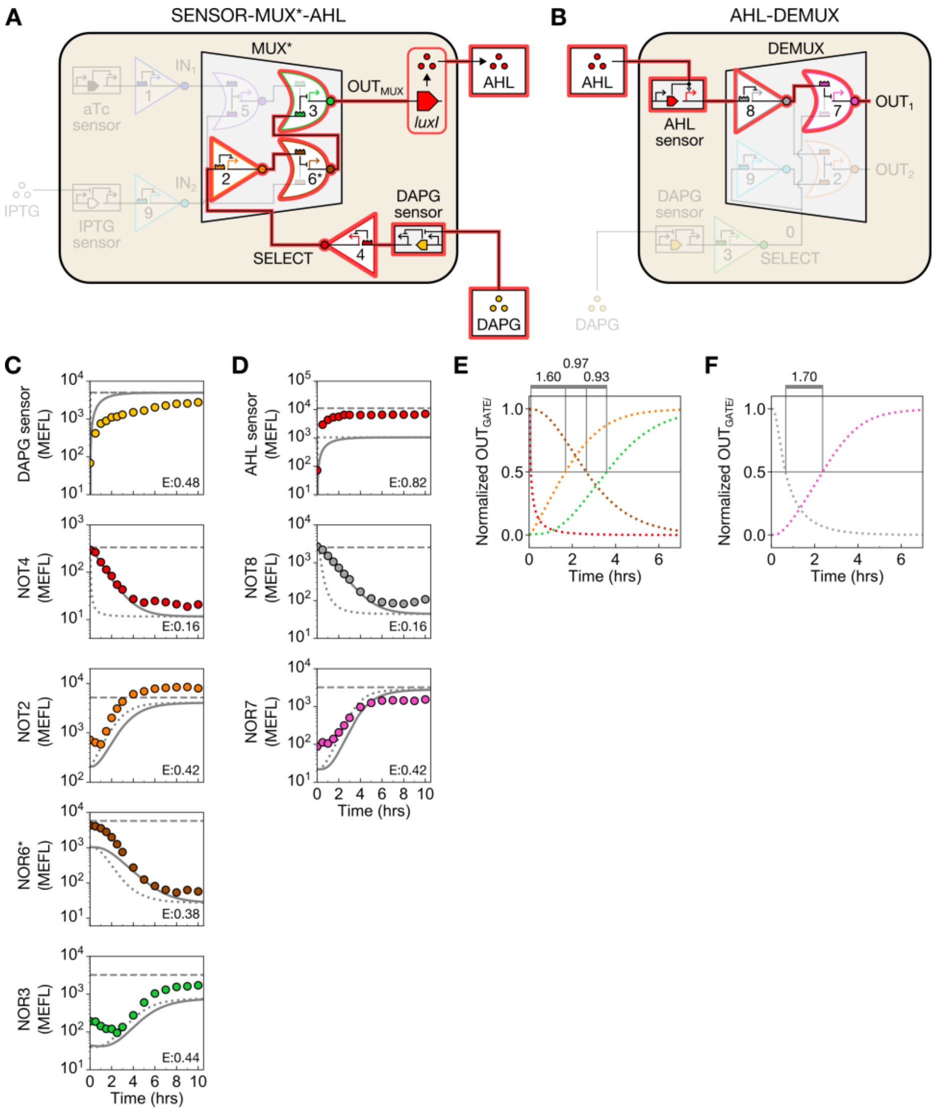

**Fig. S13. CS signal propagation dynamics.** (**A-B**) The longest computation path through the CS is shown in red. Dynamic responses of (**C**) SENSOR-MUX*-AHL cells to DAPG and (**D**) AHL-DEMUX cells to AHL. Colored circles indicate mean sfGFP fluorescence. Solid gray lines show simulated mean sfGFP fluorescence, and dotted gray lines show simulated transcriptional output via a sfGFP proxy signal that can be thought of as a set point (**Supplementary Text**). Dashed lines show the maximum gate output when the sgRNA is absent, sensor output when the sensor transcription factor is absent (DAPG sensor), or sensor output at max activation (AHL sensor). (**E-F**) Simulated gate transcriptional outputs overlaid to calculate signal propagation delay. Output signals were $\log_{10}$-transformed and then normalized before calculating when each output crosses 0.5 ($t_{1/2}$). The duration between $t_{1/2}$ times for consecutive gates is indicate above each plot in hours.

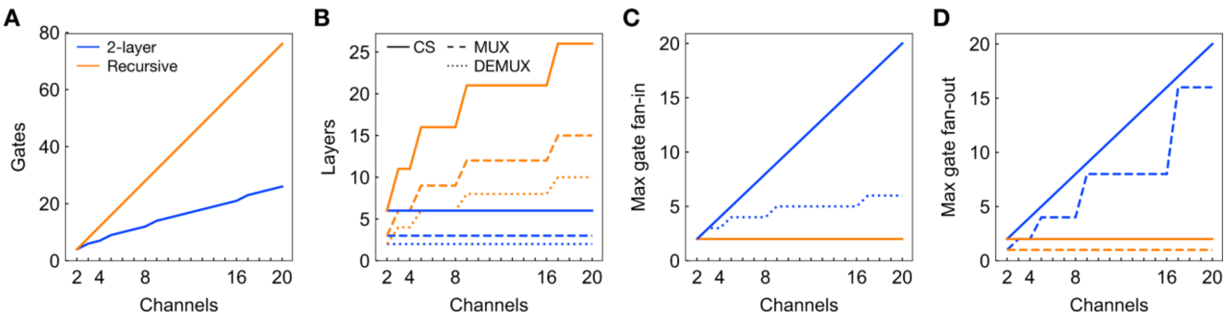

**Fig. S14. CS scaling laws.** Larger MUX and DEMUX circuits can be constructed in many ways, each subject to different engineering tradeoffs. Two general approaches emerge, however, wherein larger circuits are constructed by linking smaller circuits of the same type ("Recursive") or by connecting one layer of NOT gates to one (DEMUX) or two (MUX) layers of NOR gates ("2-layer"). While considerably fewer (**A**) gates (and therefore promoter:repressor pairs) and (**B**) layers are required for the 2-layer approach, the (**C-D**) maximum number of incoming (fan-in) and outgoing (fan-out) gate connections required quickly extends beyond those currently demonstrated for repressor-based gates in synthetic biology (which is three (*47*) and two (*21*, *22*), respectively). While nothing fundamentally precludes higher gate fan-in and fan-out, larger CSs would likely combine both approaches (e.g. an 8-channel MUX via two 2-layer 4-channel MUXs connected to one 2-channel MUX, requiring 18 gates, 6 layers, and max gate fan-in and fan-out of 4 and 2, respectively). To achieve large circuits, multiple orthogonal gate technologies could be combined to increase the number of gates available in a single cell (e.g. CRISPRi, TetR homologs, transcription activator-like effector repressors (TALERs), and zinc-finger nucleases (ZFNs)), with upwards of 30 gates theoretically possible using libraries already published (*20*, *24*). For plots, MUX and DEMUX relationships are only shown if they don't match the corresponding CS relationship.

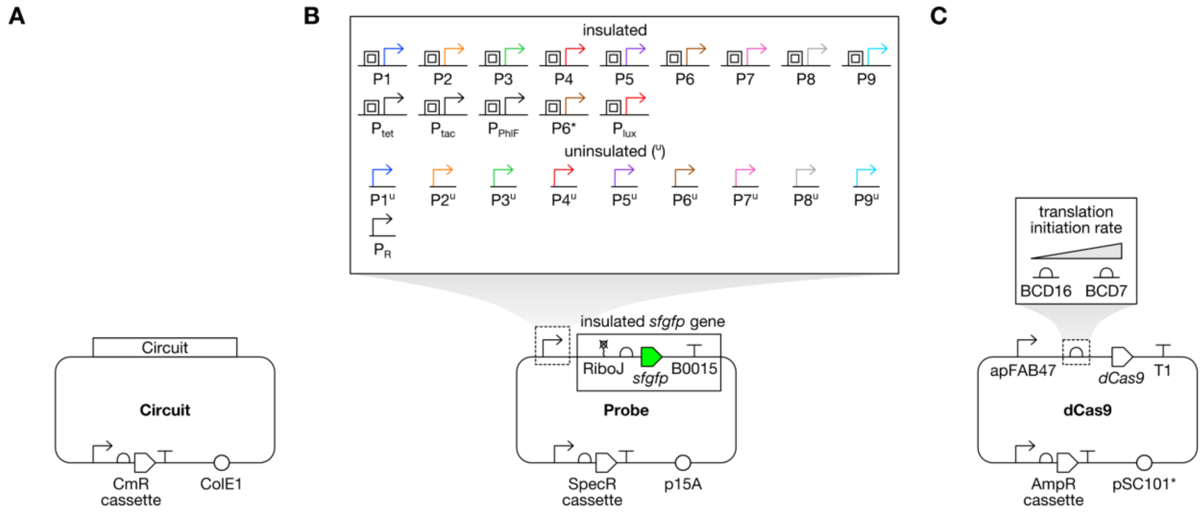

**Fig. S15. Plasmid maps.** Schematics of (**A**) circuit, (**B**) probe, and (**C**) dCas9 plasmids used in this study. Probe plasmids harboring uninsulated P1-P9 and the dCas9 plasmid pSC31_1 harboring the weak RBS BCD16 were only used in the orthogonality assay. Not pictured: empty circuit and probe plasmids, which only contain antibiotic resistance cassettes and origins of replication, and a constitutive mCherry expression plasmid (pJS0205), wherein mCherry was expressed by P2 and a synthetic RBS. Circuit maps are listed in **Fig. S16**.

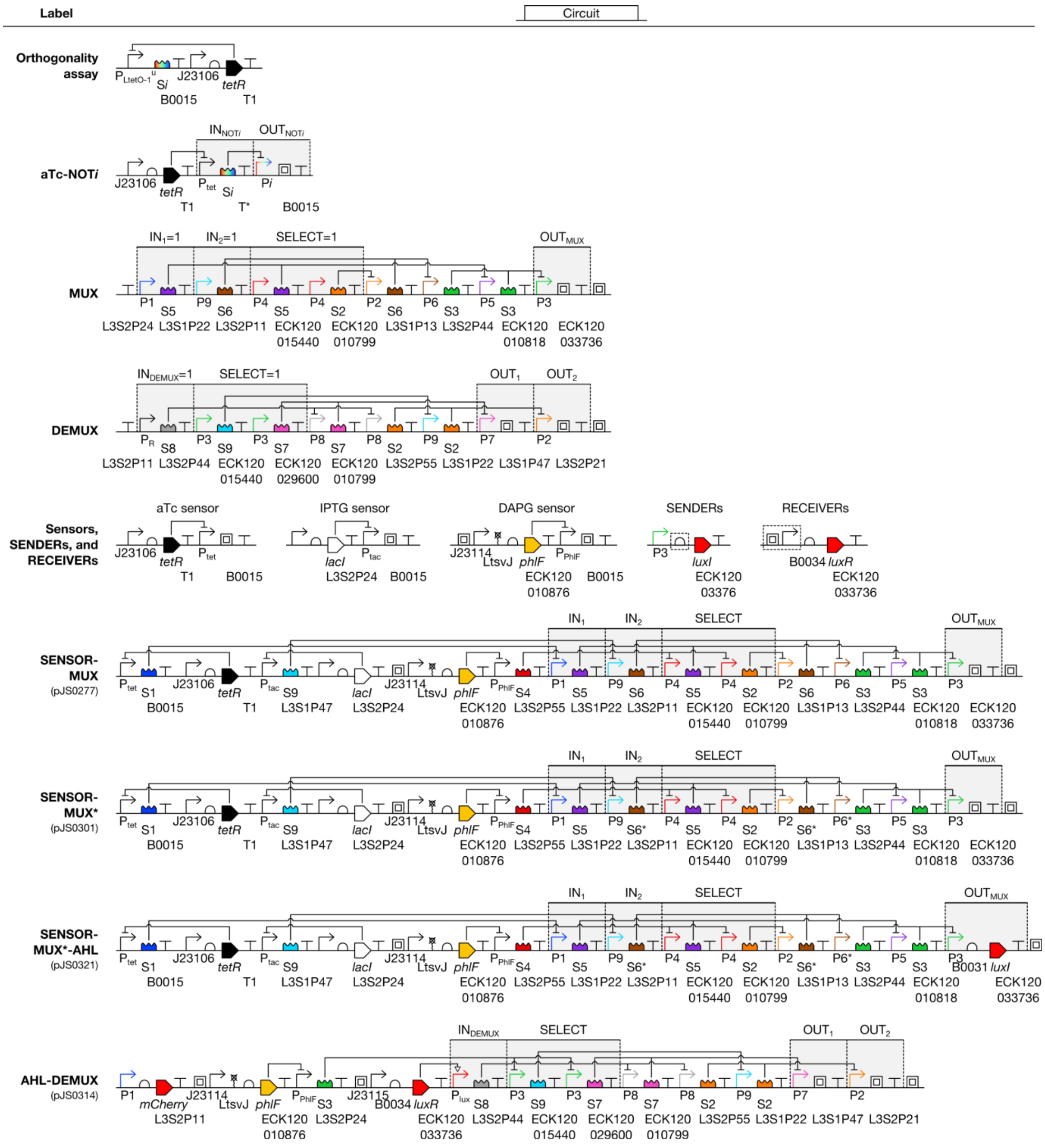

**Fig. S16. Genetic device schematics.** Circuits were carried on circuit plasmids (**Fig. S15**). Transcription units relevant to signals discussed in the text are highlighted in gray. Additional MUX and DEMUX plasmids were constructed by omitting transcription units corresponding to different input signals. SENDER RBSs and RECEIVER promoters (dashed boxes) were optimized as discussed in **Fig. S9**. $P_{tet}$ differs from $P_{LtetO\text{-}1}$ by an insulator.

### Supplementary Tables

**Table S1. NOT gate transfer function model parameters.** Parameter fits to unconstrained Hill functions, their standard errors (s.e.), and gate dynamic ranges are given.

| Gate | P$i_{max}$ (MEFL) | GATE$i_{min}$ (MEFL) | s.e. (MEFL) | $K$ (MEFL) | s.e. (MEFL) | $n$ (unitless) | s.e. (unitless) | Dynamic range (fold change) |
|---|---|---|---|---|---|---|---|---|
| 1 | 292 | 21 | 36 | 17 | 14 | 0.69 | 0.54 | 13.9 |
| 2 | 5194 | 37 | 13 | 29 | 2.5 | 1.39 | 0.11 | 141.2 |
| 3 | 3208 | 8.7 | 20 | 48 | 13 | 1.49 | 0.36 | 364.0 |
| 4 | 335 | 9.2 | 18 | 65 | 28 | 1.15 | 0.57 | 35.8 |
| 5 | 1339 | 17 | 7.8 | 82 | 18 | 2.31 | 0.60 | 80.2 |
| 6 | 659 | 12 | 18 | 136 | 36 | 1.70 | 0.63 | 55.0 |
| 7 | 3213 | 11 | 16 | 146 | 30 | 1.98 | 0.37 | 294.8 |
| 8 | 2549 | 15 | 11 | 234 | 39 | 2.99 | 0.67 | 167.5 |
| 9 | 4398 | 29 | 12 | 339 | 32 | 4.29 | 0.75 | 151.3 |
| 6* | 5756 | 0 | † | 77 | † | 1.34 | † | Inf † |

† A constraint was encountered fitting NOT6*'s data (GATE6*$_{min}$=0). As a result, no standard error estimates could be calculated.

**Table S2. Measured and simulated MUX behavior.** Measured mean sfGFP fluorescence was calculated by subtracting mean autofluorescence (145 MEFL) from mean cellular fluorescence, and simulated mean cellular fluorescence was similarly calculated by adding mean autofluorescence to simulated sfGFP fluorescence.

| $IN_1$ | $IN_2$ | SELECT | Output | Measured mean fluor. (MEFL) | Simulated mean fluor. (MEFL) | Measured mean sfGFP fluor. (MEFL) | Simulated mean sfGFP fluor. (MEFL) |
|---|---|---|---|---|---|---|---|
| 0 | 0 | 0 | $OUT_{NOR5}$ | 1178.96 | 1483.33 | 1034.33 | 1338.70 |
| 0 | 1 | 0 | $OUT_{NOR5}$ | 1513.46 | 1483.33 | 1368.83 | 1338.70 |
| 1 | 0 | 0 | $OUT_{NOR5}$ | 322.63 | 227.90 | 178.00 | 83.28 |
| 1 | 1 | 0 | $OUT_{NOR5}$ | 328.80 | 227.90 | 184.17 | 83.28 |
| 0 | 0 | 1 | $OUT_{NOR5}$ | 217.78 | 210.43 | 73.15 | 65.81 |
| 0 | 1 | 1 | $OUT_{NOR5}$ | 195.41 | 210.43 | 50.79 | 65.81 |
| 1 | 0 | 1 | $OUT_{NOR5}$ | 172.20 | 173.21 | 27.57 | 28.58 |
| 1 | 1 | 1 | $OUT_{NOR5}$ | 169.49 | 173.21 | 24.86 | 28.58 |
| 0 | 0 | 0 | $OUT_{NOR3}$ | 189.77 | 175.04 | 45.15 | 30.41 |
| 0 | 1 | 0 | $OUT_{NOR3}$ | 185.76 | 175.06 | 41.13 | 30.43 |
| 1 | 0 | 0 | $OUT_{NOR3}$ | 2491.02 | 978.05 | 2346.39 | 833.43 |
| 1 | 1 | 0 | $OUT_{NOR3}$ | 2655.27 | 986.28 | 2510.64 | 841.65 |
| 0 | 0 | 1 | $OUT_{NOR3}$ | 178.62 | 352.08 | 34.00 | 207.45 |
| 0 | 1 | 1 | $OUT_{NOR3}$ | 2848.42 | 1168.71 | 2703.80 | 1024.08 |
| 1 | 0 | 1 | $OUT_{NOR3}$ | 176.08 | 393.23 | 31.45 | 248.60 |
| 1 | 1 | 1 | $OUT_{NOR3}$ | 2689.96 | 1894.02 | 2545.33 | 1749.39 |
| 0 | 0 | 0 | $OUT_{NOT2}$ | 4672.02 | 5339.02 | 4527.40 | 5194.39 |
| 0 | 1 | 0 | $OUT_{NOT2}$ | 4688.78 | 5339.02 | 4544.16 | 5194.39 |
| 1 | 0 | 0 | $OUT_{NOT2}$ | 4463.47 | 5339.02 | 4318.85 | 5194.39 |
| 1 | 1 | 0 | $OUT_{NOT2}$ | 4726.87 | 5339.02 | 4582.24 | 5194.39 |
| 0 | 0 | 1 | $OUT_{NOT2}$ | 234.25 | 349.74 | 89.63 | 205.11 |
| 0 | 1 | 1 | $OUT_{NOT2}$ | 213.46 | 349.74 | 68.83 | 205.11 |
| 1 | 0 | 1 | $OUT_{NOT2}$ | 203.34 | 349.74 | 58.72 | 205.11 |
| 1 | 1 | 1 | $OUT_{NOT2}$ | 224.18 | 349.74 | 79.55 | 205.11 |
| 0 | 0 | 0 | $OUT_{NOR6}$ | 156.22 | 157.94 | 11.59 | 13.31 |
| 0 | 1 | 0 | $OUT_{NOR6}$ | 158.02 | 157.07 | 13.39 | 12.45 |
| 1 | 0 | 0 | $OUT_{NOR6}$ | 165.70 | 157.94 | 21.08 | 13.31 |
| 1 | 1 | 0 | $OUT_{NOR6}$ | 161.64 | 157.07 | 17.02 | 12.45 |
| 0 | 0 | 1 | $OUT_{NOR6}$ | 552.82 | 372.45 | 408.20 | 227.82 |
| 0 | 1 | 1 | $OUT_{NOR6}$ | 155.03 | 158.24 | 10.40 | 13.62 |
| 1 | 0 | 1 | $OUT_{NOR6}$ | 526.28 | 372.45 | 381.65 | 227.82 |
| 1 | 1 | 1 | $OUT_{NOR6}$ | 160.91 | 158.24 | 16.28 | 13.62 |

**Table S3. Measured and simulated DEMUX behavior.** Fluorescence signals were calculated as described in **Table S2**.

| $IN_{DEMUX}$ | SELECT | Output | Measured mean fluor. (MEFL) | Simulated mean fluor. (MEFL) | Measured mean sfGFP fluor. (MEFL) | Simulated mean sfGFP fluor. (MEFL) |
|---|---|---|---|---|---|---|
| 0 | 0 | $OUT_{NOT8}$ | 3283.33 | 2693.88 | 3138.70 | 2549.25 |
| 1 | 0 | $OUT_{NOT8}$ | 242.30 | 160.02 | 97.67 | 15.39 |
| 0 | 1 | $OUT_{NOT8}$ | 4105.89 | 2693.88 | 3961.26 | 2549.25 |
| 1 | 1 | $OUT_{NOT8}$ | 290.82 | 160.02 | 146.20 | 15.39 |
| 0 | 0 | $OUT_{NOR7}$ | 279.90 | 166.53 | 135.27 | 21.90 |
| 1 | 0 | $OUT_{NOR7}$ | 2352.52 | 3321.28 | 2207.89 | 3176.65 |
| 0 | 1 | $OUT_{NOR7}$ | 261.41 | 157.72 | 116.79 | 13.09 |
| 1 | 1 | $OUT_{NOR7}$ | 237.80 | 162.44 | 93.18 | 17.82 |
| 0 | 0 | $OUT_{NOT9}$ | 4815.73 | 4543.07 | 4671.10 | 4398.44 |
| 1 | 0 | $OUT_{NOT9}$ | 2989.01 | 4543.07 | 2844.38 | 4398.44 |
| 0 | 1 | $OUT_{NOT9}$ | 483.53 | 173.98 | 338.90 | 29.35 |
| 1 | 1 | $OUT_{NOT9}$ | 264.99 | 173.98 | 120.36 | 29.35 |
| 0 | 0 | $OUT_{NOR2}$ | 180.75 | 184.00 | 36.12 | 39.38 |
| 1 | 0 | $OUT_{NOR2}$ | 175.64 | 186.27 | 31.02 | 41.65 |
| 0 | 1 | $OUT_{NOR2}$ | 214.17 | 191.65 | 69.54 | 47.03 |
| 1 | 1 | $OUT_{NOR2}$ | 6090.02 | 2012.86 | 5945.39 | 1868.23 |

**Table S4. Sensor transfer function model parameters.** Activating Hill models ($y(x) = \min + ((\max - \min)/(1 + (K/x)^n))$) were fit to aTc, IPTG, DAPG, and AHL sensor transfer function data (**Figs. S5, S9D**). Max and min are maximum and minimum mean sfGFP fluorescence produced by the sensor (MEFL), $K$ is ligand concentration at which the sensor is half activated (units depend on the ligand), and $n$ is the Hill coefficient (dimensionless), which describes the steepness of the transfer function. Parameter fits, their standard errors (s.e.), and sensor dynamic ranges are listed here.

| Sensor | max (MEFL) | s.e. (MEFL) | min (MEFL) | s.e. (MEFL) | $K$ | s.e. | units | $n$ (unitless) | s.e. (unitless) | Dynamic range (fold change) |
|---|---|---|---|---|---|---|---|---|---|---|
| aTc | 1875 | 919 | 32 | 4.5 | 15 | 7.6 | ng/mL | 1.81 | 0.29 | 58.7 |
| IPTG | 19628 | 892 | 62 | 4.0 | 0.33 | 0.023 | mM | 2.14 | 0.081 | 317.1 |
| DAPG | 5022 | 301 | 937 | 52 | 32 | 3.1 | µM | 3.51 | 0.79 | 5.4 |
| AHL | 11039 | 510 | 32 | 1.3 | 12 | 0.80 | nM | 1.82 | 0.047 | 349.0 |

**Table S5. Measured and simulated SENSOR-MUX behavior.** Fluorescence signals were calculated as described in **Table S2**. Inducer concentrations: 0 (0); 20 ng/mL aTc, 0.3 mM IPTG, and 100 µM DAPG (1).

| aTc | IPTG | DAPG | Output | Measured mean fluor. (MEFL) | Simulated mean fluor. (MEFL) | Measured mean sfGFP fluor. (MEFL) | Simulated mean sfGFP fluor. (MEFL) |
|---|---|---|---|---|---|---|---|
| 0 | 0 | 0 | $OUT_{aTc\ sensor}$ | 168.44 | 144.63 | 23.82 | 0.00 |
| 0 | 1 | 0 | $OUT_{aTc\ sensor}$ | 160.60 | 144.63 | 15.97 | 0.00 |
| 1 | 0 | 0 | $OUT_{aTc\ sensor}$ | 1901.48 | 1290.97 | 1756.85 | 1146.34 |
| 1 | 1 | 0 | $OUT_{aTc\ sensor}$ | 1901.89 | 1290.97 | 1757.26 | 1146.34 |
| 0 | 0 | 1 | $OUT_{aTc\ sensor}$ | 167.68 | 144.63 | 23.05 | 0.00 |
| 0 | 1 | 1 | $OUT_{aTc\ sensor}$ | 170.14 | 144.63 | 25.51 | 0.00 |
| 1 | 0 | 1 | $OUT_{aTc\ sensor}$ | 1705.60 | 1290.97 | 1560.97 | 1146.34 |
| 1 | 1 | 1 | $OUT_{aTc\ sensor}$ | 1421.22 | 1290.97 | 1276.60 | 1146.34 |
| 0 | 0 | 0 | $OUT_{IPTG\ sensor}$ | 249.02 | 144.63 | 104.39 | 0.00 |
| 0 | 1 | 0 | $OUT_{IPTG\ sensor}$ | 5345.54 | 8321.02 | 5200.92 | 8176.39 |
| 1 | 0 | 0 | $OUT_{IPTG\ sensor}$ | 202.74 | 144.63 | 58.11 | 0.00 |
| 1 | 1 | 0 | $OUT_{IPTG\ sensor}$ | 5411.36 | 8321.02 | 5266.73 | 8176.39 |
| 0 | 0 | 1 | $OUT_{IPTG\ sensor}$ | 220.21 | 144.63 | 75.58 | 0.00 |
| 0 | 1 | 1 | $OUT_{IPTG\ sensor}$ | 5324.31 | 8321.02 | 5179.69 | 8176.39 |
| 1 | 0 | 1 | $OUT_{IPTG\ sensor}$ | 202.95 | 144.63 | 58.33 | 0.00 |
| 1 | 1 | 1 | $OUT_{IPTG\ sensor}$ | 4198.54 | 8321.02 | 4053.92 | 8176.39 |
| 0 | 0 | 0 | $OUT_{DAPG\ sensor}$ | 1580.28 | 144.63 | 1435.65 | 0.00 |
| 0 | 1 | 0 | $OUT_{DAPG\ sensor}$ | 253.88 | 144.63 | 109.25 | 0.00 |
| 1 | 0 | 0 | $OUT_{DAPG\ sensor}$ | 1340.90 | 144.63 | 1196.27 | 0.00 |
| 1 | 1 | 0 | $OUT_{DAPG\ sensor}$ | 237.97 | 144.63 | 93.34 | 0.00 |
| 0 | 0 | 1 | $OUT_{DAPG\ sensor}$ | 5483.54 | 5076.89 | 5338.91 | 4932.27 |
| 0 | 1 | 1 | $OUT_{DAPG\ sensor}$ | 3584.51 | 5076.89 | 3439.89 | 4932.27 |
| 1 | 0 | 1 | $OUT_{DAPG\ sensor}$ | 4645.54 | 5076.89 | 4500.91 | 4932.27 |
| 1 | 1 | 1 | $OUT_{DAPG\ sensor}$ | 2319.26 | 5076.89 | 2174.63 | 4932.27 |
| 0 | 0 | 0 | $OUT_{NOT1}$ | 386.07 | 436.58 | 241.45 | 291.95 |
| 0 | 1 | 0 | $OUT_{NOT1}$ | 354.95 | 436.58 | 210.32 | 291.95 |
| 1 | 0 | 0 | $OUT_{NOT1}$ | 162.93 | 179.96 | 18.30 | 35.34 |
| 1 | 1 | 0 | $OUT_{NOT1}$ | 159.23 | 179.96 | 14.61 | 35.34 |
| 0 | 0 | 1 | $OUT_{NOT1}$ | 382.75 | 436.58 | 238.12 | 291.95 |
| 0 | 1 | 1 | $OUT_{NOT1}$ | 374.88 | 436.58 | 230.26 | 291.95 |
| 1 | 0 | 1 | $OUT_{NOT1}$ | 165.61 | 179.96 | 20.98 | 35.34 |
| 1 | 1 | 1 | $OUT_{NOT1}$ | 170.32 | 179.96 | 25.69 | 35.34 |
| 0 | 0 | 0 | $OUT_{NOT9}$ | 6352.68 | 4543.07 | 6208.05 | 4398.44 |
| 0 | 1 | 0 | $OUT_{NOT9}$ | 185.38 | 173.71 | 40.75 | 29.08 |
| 1 | 0 | 0 | $OUT_{NOT9}$ | 6103.66 | 4543.07 | 5959.03 | 4398.44 |
| 1 | 1 | 0 | $OUT_{NOT9}$ | 261.93 | 173.71 | 117.30 | 29.08 |
| 0 | 0 | 1 | $OUT_{NOT9}$ | 5724.95 | 4543.07 | 5580.32 | 4398.44 |
| 0 | 1 | 1 | $OUT_{NOT9}$ | 259.20 | 173.71 | 114.57 | 29.08 |
| 1 | 0 | 1 | $OUT_{NOT9}$ | 5776.32 | 4543.07 | 5631.69 | 4398.44 |
| 1 | 1 | 1 | $OUT_{NOT9}$ | 489.62 | 173.71 | 345.00 | 29.08 |
| 0 | 0 | 0 | $OUT_{NOT4}$ | 218.71 | 479.70 | 74.08 | 335.07 |
| 0 | 1 | 0 | $OUT_{NOT4}$ | 413.88 | 479.70 | 269.25 | 335.07 |
| 1 | 0 | 0 | $OUT_{NOT4}$ | 279.89 | 479.70 | 135.26 | 335.07 |
| 1 | 1 | 0 | $OUT_{NOT4}$ | 480.57 | 479.70 | 335.95 | 335.07 |
| 0 | 0 | 1 | $OUT_{NOT4}$ | 152.05 | 156.23 | 7.43 | 11.60 |
| 0 | 1 | 1 | $OUT_{NOT4}$ | 157.02 | 156.23 | 12.40 | 11.60 |
| 1 | 0 | 1 | $OUT_{NOT4}$ | 167.86 | 156.23 | 23.24 | 11.60 |
| 1 | 1 | 1 | $OUT_{NOT4}$ | 170.27 | 156.23 | 25.65 | 11.60 |

| | | | | | | | |
|---|---|---|---|---|---|---|---|
| 0 | 0 | 0 | $OUT_{NOR5}$ | 210.74 | 173.21 | 66.11 | 28.58 |
| 0 | 1 | 0 | $OUT_{NOR5}$ | 169.62 | 173.21 | 25.00 | 28.58 |
| 1 | 0 | 0 | $OUT_{NOR5}$ | 894.97 | 200.58 | 750.35 | 55.96 |
| 1 | 1 | 0 | $OUT_{NOR5}$ | 306.40 | 200.58 | 161.77 | 55.96 |
| 0 | 0 | 1 | $OUT_{NOR5}$ | 301.55 | 222.44 | 156.92 | 77.81 |
| 0 | 1 | 1 | $OUT_{NOR5}$ | 295.12 | 222.44 | 150.50 | 77.81 |
| 1 | 0 | 1 | $OUT_{NOR5}$ | 1430.96 | 1196.67 | 1286.33 | 1052.05 |
| 1 | 1 | 1 | $OUT_{NOR5}$ | 1388.10 | 1196.67 | 1243.47 | 1052.05 |
| 0 | 0 | 0 | $OUT_{NOR3}$ | 3180.92 | 1894.02 | 3036.29 | 1749.39 |
| 0 | 1 | 0 | $OUT_{NOR3}$ | 1056.24 | 439.62 | 911.61 | 295.00 |
| 1 | 0 | 0 | $OUT_{NOR3}$ | 1264.32 | 1310.16 | 1119.69 | 1165.53 |
| 1 | 1 | 0 | $OUT_{NOR3}$ | 676.36 | 398.10 | 531.74 | 253.47 |
| 0 | 0 | 1 | $OUT_{NOR3}$ | 2383.14 | 1040.23 | 2238.52 | 895.60 |
| 0 | 1 | 1 | $OUT_{NOR3}$ | 2369.44 | 1025.47 | 2224.82 | 880.84 |
| 1 | 0 | 1 | $OUT_{NOR3}$ | 270.44 | 184.19 | 125.81 | 39.56 |
| 1 | 1 | 1 | $OUT_{NOR3}$ | 224.74 | 184.13 | 80.12 | 39.50 |
| 0 | 0 | 0 | $OUT_{NOT2}$ | 4217.31 | 349.74 | 4072.68 | 205.11 |
| 0 | 1 | 0 | $OUT_{NOT2}$ | 549.84 | 349.74 | 405.21 | 205.11 |
| 1 | 0 | 0 | $OUT_{NOT2}$ | 3618.87 | 349.74 | 3474.24 | 205.11 |
| 1 | 1 | 0 | $OUT_{NOT2}$ | 567.72 | 349.74 | 423.09 | 205.11 |
| 0 | 0 | 1 | $OUT_{NOT2}$ | 7278.48 | 4212.73 | 7133.86 | 4068.11 |
| 0 | 1 | 1 | $OUT_{NOT2}$ | 7414.30 | 4212.73 | 7269.67 | 4068.11 |
| 1 | 0 | 1 | $OUT_{NOT2}$ | 7308.02 | 4212.73 | 7163.40 | 4068.11 |
| 1 | 1 | 1 | $OUT_{NOT2}$ | 6560.61 | 4212.73 | 6415.99 | 4068.11 |
| 0 | 0 | 0 | $OUT_{NOR6}$ | 155.70 | 158.24 | 11.08 | 13.62 |
| 0 | 1 | 0 | $OUT_{NOR6}$ | 501.97 | 341.36 | 357.34 | 196.73 |
| 1 | 0 | 0 | $OUT_{NOR6}$ | 157.98 | 158.24 | 13.36 | 13.62 |
| 1 | 1 | 0 | $OUT_{NOR6}$ | 590.38 | 341.36 | 445.76 | 196.73 |
| 0 | 0 | 1 | $OUT_{NOR6}$ | 153.45 | 157.19 | 8.83 | 12.56 |
| 0 | 1 | 1 | $OUT_{NOR6}$ | 151.68 | 158.60 | 7.05 | 13.97 |
| 1 | 0 | 1 | $OUT_{NOR6}$ | 160.05 | 157.19 | 15.43 | 12.56 |
| 1 | 1 | 1 | $OUT_{NOR6}$ | 168.44 | 158.60 | 23.81 | 13.97 |

**Table S6. Measured and simulated SENSOR-MUX* behavior.** Fluorescence signals were calculated as described in **Table S2**. Inducer concentrations: 0 (0); 20 ng/mL aTc, 0.3 mM IPTG, and 100 µM DAPG (1).

| aTc | IPTG | DAPG | Output | Measured mean fluor. (MEFL) | Simulated mean fluor. (MEFL) | Measured mean sfGFP fluor. (MEFL) | Simulated mean sfGFP fluor. (MEFL) |
|---|---|---|---|---|---|---|---|
| 0 | 0 | 0 | $OUT_{aTc\ sensor}$ | 161.69 | 144.63 | 17.07 | 0.00 |
| 0 | 1 | 0 | $OUT_{aTc\ sensor}$ | 155.34 | 144.63 | 10.72 | 0.00 |
| 1 | 0 | 0 | $OUT_{aTc\ sensor}$ | 1922.85 | 1290.97 | 1778.22 | 1146.34 |
| 1 | 1 | 0 | $OUT_{aTc\ sensor}$ | 1889.83 | 1290.97 | 1745.20 | 1146.34 |
| 0 | 0 | 1 | $OUT_{aTc\ sensor}$ | 166.77 | 144.63 | 22.14 | 0.00 |
| 0 | 1 | 1 | $OUT_{aTc\ sensor}$ | 161.14 | 144.63 | 16.52 | 0.00 |
| 1 | 0 | 1 | $OUT_{aTc\ sensor}$ | 1650.96 | 1290.97 | 1506.33 | 1146.34 |
| 1 | 1 | 1 | $OUT_{aTc\ sensor}$ | 1452.75 | 1290.97 | 1308.13 | 1146.34 |
| 0 | 0 | 0 | $OUT_{IPTG\ sensor}$ | 243.28 | 144.63 | 98.66 | 0.00 |
| 0 | 1 | 0 | $OUT_{IPTG\ sensor}$ | 5488.74 | 8321.02 | 5344.11 | 8176.39 |
| 1 | 0 | 0 | $OUT_{IPTG\ sensor}$ | 233.58 | 144.63 | 88.95 | 0.00 |
| 1 | 1 | 0 | $OUT_{IPTG\ sensor}$ | 5310.42 | 8321.02 | 5165.79 | 8176.39 |
| 0 | 0 | 1 | $OUT_{IPTG\ sensor}$ | 261.67 | 144.63 | 117.04 | 0.00 |
| 0 | 1 | 1 | $OUT_{IPTG\ sensor}$ | 4938.10 | 8321.02 | 4793.47 | 8176.39 |
| 1 | 0 | 1 | $OUT_{IPTG\ sensor}$ | 218.61 | 144.63 | 73.98 | 0.00 |
| 1 | 1 | 1 | $OUT_{IPTG\ sensor}$ | 4075.19 | 8321.02 | 3930.57 | 8176.39 |
| 0 | 0 | 0 | $OUT_{DAPG\ sensor}$ | 2159.95 | 1460.59 | 2015.32 | 1315.96 |
| 0 | 1 | 0 | $OUT_{DAPG\ sensor}$ | 242.55 | 144.63 | 97.92 | 0.00 |
| 1 | 0 | 0 | $OUT_{DAPG\ sensor}$ | 1381.08 | 1460.59 | 1236.45 | 1315.96 |
| 1 | 1 | 0 | $OUT_{DAPG\ sensor}$ | 220.16 | 144.63 | 75.53 | 0.00 |
| 0 | 0 | 1 | $OUT_{DAPG\ sensor}$ | 4885.01 | 5076.89 | 4740.38 | 4932.27 |
| 0 | 1 | 1 | $OUT_{DAPG\ sensor}$ | 2896.49 | 5076.89 | 2751.87 | 4932.27 |
| 1 | 0 | 1 | $OUT_{DAPG\ sensor}$ | 3927.94 | 5076.89 | 3783.31 | 4932.27 |
| 1 | 1 | 1 | $OUT_{DAPG\ sensor}$ | 2157.77 | 5076.89 | 2013.14 | 4932.27 |
| 0 | 0 | 0 | $OUT_{NOT1}$ | 347.89 | 436.58 | 203.26 | 291.95 |
| 0 | 1 | 0 | $OUT_{NOT1}$ | 350.67 | 436.58 | 206.04 | 291.95 |
| 1 | 0 | 0 | $OUT_{NOT1}$ | 160.71 | 179.96 | 16.08 | 35.34 |
| 1 | 1 | 0 | $OUT_{NOT1}$ | 160.03 | 179.96 | 15.40 | 35.34 |
| 0 | 0 | 1 | $OUT_{NOT1}$ | 395.46 | 436.58 | 250.83 | 291.95 |
| 0 | 1 | 1 | $OUT_{NOT1}$ | 358.70 | 436.58 | 214.07 | 291.95 |
| 1 | 0 | 1 | $OUT_{NOT1}$ | 164.30 | 179.96 | 19.67 | 35.34 |
| 1 | 1 | 1 | $OUT_{NOT1}$ | 169.32 | 179.96 | 24.69 | 35.34 |
| 0 | 0 | 0 | $OUT_{NOT9}$ | 2751.51 | 4543.07 | 2606.88 | 4398.44 |
| 0 | 1 | 0 | $OUT_{NOT9}$ | 182.34 | 173.71 | 37.72 | 29.08 |
| 1 | 0 | 0 | $OUT_{NOT9}$ | 3318.15 | 4543.07 | 3173.52 | 4398.44 |
| 1 | 1 | 0 | $OUT_{NOT9}$ | 355.19 | 173.71 | 210.57 | 29.08 |
| 0 | 0 | 1 | $OUT_{NOT9}$ | 3080.04 | 4543.07 | 2935.42 | 4398.44 |
| 0 | 1 | 1 | $OUT_{NOT9}$ | 194.28 | 173.71 | 49.66 | 29.08 |
| 1 | 0 | 1 | $OUT_{NOT9}$ | 3023.99 | 4543.07 | 2879.37 | 4398.44 |
| 1 | 1 | 1 | $OUT_{NOT9}$ | 299.40 | 173.71 | 154.77 | 29.08 |
| 0 | 0 | 0 | $OUT_{NOT4}$ | 196.99 | 164.01 | 52.37 | 19.38 |
| 0 | 1 | 0 | $OUT_{NOT4}$ | 396.19 | 479.70 | 251.57 | 335.07 |
| 1 | 0 | 0 | $OUT_{NOT4}$ | 280.28 | 164.01 | 135.66 | 19.38 |
| 1 | 1 | 0 | $OUT_{NOT4}$ | 454.15 | 479.70 | 309.53 | 335.07 |
| 0 | 0 | 1 | $OUT_{NOT4}$ | 164.69 | 156.23 | 20.06 | 11.60 |
| 0 | 1 | 1 | $OUT_{NOT4}$ | 161.09 | 156.23 | 16.46 | 11.60 |
| 1 | 0 | 1 | $OUT_{NOT4}$ | 175.47 | 156.23 | 30.85 | 11.60 |
| 1 | 1 | 1 | $OUT_{NOT4}$ | 175.55 | 156.23 | 30.92 | 11.60 |

| | | | | | | | |
|---|---|---|---|---|---|---|---|
| 0 | 0 | 0 | $OUT_{NOR5}$ | 206.31 | 219.12 | 61.68 | 74.49 |
| 0 | 1 | 0 | $OUT_{NOR5}$ | 191.47 | 173.21 | 46.84 | 28.58 |
| 1 | 0 | 0 | $OUT_{NOR5}$ | 643.63 | 1109.34 | 499.00 | 964.71 |
| 1 | 1 | 0 | $OUT_{NOR5}$ | 480.34 | 200.58 | 335.71 | 55.96 |
| 0 | 0 | 1 | $OUT_{NOR5}$ | 280.94 | 222.44 | 136.31 | 77.81 |
| 0 | 1 | 1 | $OUT_{NOR5}$ | 242.97 | 222.44 | 98.34 | 77.81 |
| 1 | 0 | 1 | $OUT_{NOR5}$ | 1327.02 | 1196.67 | 1182.40 | 1052.05 |
| 1 | 1 | 1 | $OUT_{NOR5}$ | 1212.87 | 1196.67 | 1068.24 | 1052.05 |
| 0 | 0 | 0 | $OUT_{NOR3}$ | 921.22 | 1086.01 | 776.60 | 941.38 |
| 0 | 1 | 0 | $OUT_{NOR3}$ | 259.96 | 183.44 | 115.33 | 38.81 |
| 1 | 0 | 0 | $OUT_{NOR3}$ | 656.00 | 188.37 | 511.37 | 43.75 |
| 1 | 1 | 0 | $OUT_{NOR3}$ | 192.12 | 182.35 | 47.49 | 37.73 |
| 0 | 0 | 1 | $OUT_{NOR3}$ | 1459.87 | 1064.06 | 1315.25 | 919.43 |
| 0 | 1 | 1 | $OUT_{NOR3}$ | 1605.09 | 902.61 | 1460.46 | 757.99 |
| 1 | 0 | 1 | $OUT_{NOR3}$ | 284.90 | 184.28 | 140.27 | 39.66 |
| 1 | 1 | 1 | $OUT_{NOR3}$ | 206.47 | 183.57 | 61.84 | 38.94 |
| 0 | 0 | 0 | $OUT_{NOT2}$ | 3156.88 | 3468.42 | 3012.25 | 3323.79 |
| 0 | 1 | 0 | $OUT_{NOT2}$ | 600.29 | 349.74 | 455.66 | 205.11 |
| 1 | 0 | 0 | $OUT_{NOT2}$ | 2811.95 | 3468.42 | 2667.32 | 3323.79 |
| 1 | 1 | 0 | $OUT_{NOT2}$ | 965.92 | 349.74 | 821.30 | 205.11 |
| 0 | 0 | 1 | $OUT_{NOT2}$ | 6733.03 | 4212.73 | 6588.41 | 4068.11 |
| 0 | 1 | 1 | $OUT_{NOT2}$ | 6381.32 | 4212.73 | 6236.70 | 4068.11 |
| 1 | 0 | 1 | $OUT_{NOT2}$ | 6374.93 | 4212.73 | 6230.30 | 4068.11 |
| 1 | 1 | 1 | $OUT_{NOT2}$ | 5937.29 | 4212.73 | 5792.67 | 4068.11 |
| 0 | 0 | 0 | $OUT_{NOR6*}$ | 206.24 | 156.34 | 61.61 | 11.71 |
| 0 | 1 | 0 | $OUT_{NOR6*}$ | 4215.95 | 1198.58 | 4071.32 | 1053.95 |
| 1 | 0 | 0 | $OUT_{NOR6*}$ | 285.37 | 156.34 | 140.74 | 11.71 |
| 1 | 1 | 0 | $OUT_{NOR6*}$ | 4547.17 | 1198.58 | 4402.54 | 1053.95 |
| 0 | 0 | 1 | $OUT_{NOR6*}$ | 189.26 | 154.98 | 44.63 | 10.35 |
| 0 | 1 | 1 | $OUT_{NOR6*}$ | 192.70 | 172.01 | 48.07 | 27.38 |
| 1 | 0 | 1 | $OUT_{NOR6*}$ | 221.82 | 154.98 | 77.20 | 10.35 |
| 1 | 1 | 1 | $OUT_{NOR6*}$ | 226.84 | 172.01 | 82.21 | 27.38 |

**Table S7. Measured and simulated AHL-DEMUX behavior.** Fluorescence signals were calculated as described for **Table S2**. Inducer concentrations: 0 (0); 3.5 nM AHL, 100 µM DAPG (1).

| AHL | DAPG | Output | Measured mean fluor. (MEFL) | Simulated mean fluor. (MEFL) | Measured mean sfGFP fluor. (MEFL) | Simulated mean sfGFP fluor. (MEFL) |
|---|---|---|---|---|---|---|
| 0 | 0 | $OUT_{AHL\ sensor}$ | 266.10 | 144.63 | 121.47 | 0.00 |
| 1 | 0 | $OUT_{AHL\ sensor}$ | 7104.58 | 1185.52 | 6959.95 | 1040.90 |
| 0 | 1 | $OUT_{AHL\ sensor}$ | 258.52 | 144.63 | 113.89 | 0.00 |
| 1 | 1 | $OUT_{AHL\ sensor}$ | 6233.05 | 1185.52 | 6088.42 | 1040.90 |
| 0 | 0 | $OUT_{DAPG\ sensor}$ | 401.75 | 144.63 | 257.12 | 0.00 |
| 1 | 0 | $OUT_{DAPG\ sensor}$ | 244.53 | 144.63 | 99.91 | 0.00 |
| 0 | 1 | $OUT_{DAPG\ sensor}$ | 3912.75 | 5076.89 | 3768.13 | 4932.27 |
| 1 | 1 | $OUT_{DAPG\ sensor}$ | 3286.28 | 5076.89 | 3141.66 | 4932.27 |
| 0 | 0 | $OUT_{NOT3}$ | 4586.74 | 3352.53 | 4442.11 | 3207.91 |
| 1 | 0 | $OUT_{NOT3}$ | 4395.24 | 3352.53 | 4250.61 | 3207.91 |
| 0 | 1 | $OUT_{NOT3}$ | 652.93 | 156.60 | 508.31 | 11.97 |
| 1 | 1 | $OUT_{NOT3}$ | 389.86 | 156.60 | 245.24 | 11.97 |
| 0 | 0 | $OUT_{NOT8}$ | 3806.48 | 2693.88 | 3661.86 | 2549.25 |
| 1 | 0 | $OUT_{NOT8}$ | 286.69 | 188.95 | 142.06 | 44.32 |
| 0 | 1 | $OUT_{NOT8}$ | 2968.11 | 2693.88 | 2823.49 | 2549.25 |
| 1 | 1 | $OUT_{NOT8}$ | 249.55 | 188.95 | 104.93 | 44.32 |
| 0 | 0 | $OUT_{NOR7}$ | 218.98 | 157.72 | 74.36 | 13.09 |
| 1 | 0 | $OUT_{NOR7}$ | 258.91 | 162.32 | 114.28 | 17.69 |
| 0 | 1 | $OUT_{NOR7}$ | 252.63 | 166.43 | 108.00 | 21.80 |
| 1 | 1 | $OUT_{NOR7}$ | 1864.01 | 2940.20 | 1719.38 | 2795.57 |
| 0 | 0 | $OUT_{NOT9}$ | 518.40 | 173.98 | 373.78 | 29.35 |
| 1 | 0 | $OUT_{NOT9}$ | 254.13 | 173.98 | 109.50 | 29.35 |
| 0 | 1 | $OUT_{NOT9}$ | 4985.35 | 4543.07 | 4840.72 | 4398.44 |
| 1 | 1 | $OUT_{NOT9}$ | 3250.55 | 4543.07 | 3105.92 | 4398.44 |
| 0 | 0 | $OUT_{NOR2}$ | 209.84 | 191.65 | 65.21 | 47.03 |
| 1 | 0 | $OUT_{NOR2}$ | 7345.26 | 1296.13 | 7200.64 | 1151.51 |
| 0 | 1 | $OUT_{NOR2}$ | 191.98 | 184.00 | 47.35 | 39.38 |
| 1 | 1 | $OUT_{NOR2}$ | 538.78 | 186.23 | 394.15 | 41.60 |

**Table S8. Measured and simulated CS behavior.** Simulated fluorescence signals were reproduced from individual SENSOR-MUX* and AHL-DEMUX characterizations (**Tables S6 and S7**). The $OUT_{NOR3}$ measurements from coculture set 2 are shown in **Fig. 4**, although both sets looked nearly identical. Inducer concentrations: 0 (0); 20ng/mL aTc, 0.3mM IPTG, and 100µM DAPG (1).

| aTc | IPTG | DAPG | Output | Coculture | Measured mean fluor. (MEFL) | Simulated mean fluor. (MEFL) | Measured mean sfGFP fluor. (MEFL) | Simulated mean sfGFP fluor. (MEFL) |
|---|---|---|---|---|---|---|---|---|
| 0 | 0 | 1 | $OUT_{NOR3}$ | 1 | 2528.38 | 1064.06 | 2383.75 | 919.43 |
| 0 | 1 | 1 | $OUT_{NOR3}$ | 1 | 2096.63 | 902.61 | 1952.00 | 757.99 |
| 1 | 0 | 1 | $OUT_{NOR3}$ | 1 | 442.37 | 184.28 | 297.75 | 39.66 |
| 1 | 1 | 1 | $OUT_{NOR3}$ | 1 | 244.87 | 183.57 | 100.24 | 38.94 |
| 0 | 0 | 0 | $OUT_{NOR3}$ | 1 | 2026.07 | 1086.01 | 1881.44 | 941.38 |
| 0 | 1 | 0 | $OUT_{NOR3}$ | 1 | 173.29 | 183.44 | 28.66 | 38.81 |
| 1 | 0 | 0 | $OUT_{NOR3}$ | 1 | 1586.82 | 188.37 | 1442.19 | 43.75 |
| 1 | 1 | 0 | $OUT_{NOR3}$ | 1 | 213.73 | 182.35 | 69.10 | 37.73 |
| 0 | 0 | 1 | $OUT_{NOR7}$ | 1 | 2419.71 | 2940.20 | 2275.08 | 2795.57 |
| 0 | 1 | 1 | $OUT_{NOR7}$ | 1 | 2147.22 | 2940.20 | 2002.59 | 2795.57 |
| 1 | 0 | 1 | $OUT_{NOR7}$ | 1 | 742.79 | 166.43 | 598.17 | 21.80 |
| 1 | 1 | 1 | $OUT_{NOR7}$ | 1 | 340.63 | 166.43 | 196.00 | 21.80 |
| 0 | 0 | 0 | $OUT_{NOR7}$ | 1 | 721.22 | 162.32 | 576.59 | 17.69 |
| 0 | 1 | 0 | $OUT_{NOR7}$ | 1 | 249.46 | 157.72 | 104.83 | 13.09 |
| 1 | 0 | 0 | $OUT_{NOR7}$ | 1 | 362.27 | 162.32 | 217.64 | 17.69 |
| 1 | 1 | 0 | $OUT_{NOR7}$ | 1 | 247.58 | 157.72 | 102.95 | 13.09 |
| 0 | 0 | 1 | $OUT_{NOR3}$ | 2 | 2488.58 | 1064.06 | 2343.95 | 919.43 |
| 0 | 1 | 1 | $OUT_{NOR3}$ | 2 | 2155.84 | 902.61 | 2011.21 | 757.99 |
| 1 | 0 | 1 | $OUT_{NOR3}$ | 2 | 423.62 | 184.28 | 278.99 | 39.66 |
| 1 | 1 | 1 | $OUT_{NOR3}$ | 2 | 248.62 | 183.57 | 104.00 | 38.94 |
| 0 | 0 | 0 | $OUT_{NOR3}$ | 2 | 1942.34 | 1086.01 | 1797.72 | 941.38 |
| 0 | 1 | 0 | $OUT_{NOR3}$ | 2 | 175.72 | 183.44 | 31.10 | 38.81 |
| 1 | 0 | 0 | $OUT_{NOR3}$ | 2 | 1485.66 | 188.37 | 1341.04 | 43.75 |
| 1 | 1 | 0 | $OUT_{NOR3}$ | 2 | 211.30 | 182.35 | 66.67 | 37.73 |
| 0 | 0 | 1 | $OUT_{NOR2}$ | 2 | 796.40 | 186.23 | 651.78 | 41.60 |
| 0 | 1 | 1 | $OUT_{NOR2}$ | 2 | 2047.01 | 186.23 | 1902.38 | 41.60 |
| 1 | 0 | 1 | $OUT_{NOR2}$ | 2 | 208.76 | 184.00 | 64.13 | 39.38 |
| 1 | 1 | 1 | $OUT_{NOR2}$ | 2 | 182.98 | 184.00 | 38.36 | 39.38 |
| 0 | 0 | 0 | $OUT_{NOR2}$ | 2 | 8980.22 | 1296.13 | 8835.59 | 1151.51 |
| 0 | 1 | 0 | $OUT_{NOR2}$ | 2 | 318.49 | 191.65 | 173.86 | 47.03 |
| 1 | 0 | 0 | $OUT_{NOR2}$ | 2 | 5669.10 | 1296.13 | 5524.47 | 1151.51 |
| 1 | 1 | 0 | $OUT_{NOR2}$ | 2 | 244.92 | 191.65 | 100.30 | 47.03 |

**Table S9. Total sgRNA expressed by probed circuits.** This table shows numerical values from **Fig. S12C**. Expression levels of the constitutive promoters driving $IN_1$, $IN_2$, and SELECT in the MUX and $IN_{DEMUX}$ and SELECT in the DEMUX were taken to be 0 or the sfGFP fluorescence produced by P1, P9, P4, $P_R$, and P3, respectively, in the absence of repressor, which is 292, 4398, 335, 5864, and 3208 MEFL.

| Circuit | Inputs | | Select | sgRNA (MEFL) | | | | | | | | | Total |
|---|---|---|---|---|---|---|---|---|---|---|---|---|---|
| | | | | S1 | S2 | S3 | S4 | S5 | S6/S6* | S7 | S8 | S9 | |
| MUX | $IN_1$ | $IN_2$ | SELECT | | | | | | | | | | |
| | 0 | 0 | 0 | 0 | 0 | 1046 | 0 | 0 | 4527 | 0 | 0 | 0 | 5573 |
| | 0 | 1 | 0 | 0 | 0 | 1382 | 0 | 0 | 8943 | 0 | 0 | 0 | 10325 |
| | 1 | 0 | 0 | 0 | 0 | 199 | 0 | 292 | 4319 | 0 | 0 | 0 | 4810 |
| | 1 | 1 | 0 | 0 | 0 | 201 | 0 | 292 | 8981 | 0 | 0 | 0 | 9474 |
| | 0 | 0 | 1 | 0 | 335 | 481 | 0 | 335 | 90 | 0 | 0 | 0 | 1241 |
| | 0 | 1 | 1 | 0 | 335 | 61 | 0 | 335 | 4467 | 0 | 0 | 0 | 5199 |
| | 1 | 0 | 1 | 0 | 335 | 409 | 0 | 627 | 59 | 0 | 0 | 0 | 1430 |
| | 1 | 1 | 1 | 0 | 335 | 41 | 0 | 627 | 4478 | 0 | 0 | 0 | 5481 |
| DEMUX | $IN_{DEMUX}$ | | SELECT | | | | | | | | | | |
| | 0 | | 0 | 0 | 7810 | 0 | 0 | 0 | 0 | 3139 | 0 | 0 | 10949 |
| | 1 | | 0 | 0 | 2942 | 0 | 0 | 0 | 0 | 98 | 5864 | 0 | 8904 |
| | 0 | | 1 | 0 | 4300 | 0 | 0 | 0 | 0 | 7169 | 0 | 3208 | 14677 |
| | 1 | | 1 | 0 | 267 | 0 | 0 | 0 | 0 | 3354 | 5864 | 3208 | 12693 |
| SENSOR-MUX* | aTc | IPTG | DAPG | | | | | | | | | | |
| | 0 | 0 | 0 | 17 | 52 | 123 | 2015 | 256 | 5619 | 0 | 0 | 99 | 8181 |
| | 0 | 1 | 0 | 11 | 252 | 4118 | 98 | 458 | 493 | 0 | 0 | 5344 | 10773 |
| | 1 | 0 | 0 | 1778 | 136 | 640 | 1236 | 152 | 5841 | 0 | 0 | 89 | 9872 |
| | 1 | 1 | 0 | 1745 | 310 | 4738 | 76 | 325 | 1032 | 0 | 0 | 5166 | 13391 |
| | 0 | 0 | 1 | 22 | 20 | 181 | 4740 | 271 | 9524 | 0 | 0 | 117 | 14875 |
| | 0 | 1 | 1 | 17 | 16 | 146 | 2752 | 231 | 6286 | 0 | 0 | 4793 | 14242 |
| | 1 | 0 | 1 | 1506 | 31 | 1260 | 3783 | 51 | 9110 | 0 | 0 | 74 | 15814 |
| | 1 | 1 | 1 | 1308 | 31 | 1150 | 2013 | 56 | 5947 | 0 | 0 | 3931 | 14436 |
| AHL-DEMUX | AHL (nM) | | DAPG | | | | | | | | | | |
| | 0 | | 1 | 0 | 7664 | 3768 | 0 | 0 | 0 | 3332 | 114 | 508 | 15386 |
| | 3.5 | | 1 | 0 | 3211 | 3142 | 0 | 0 | 0 | 350 | 6088 | 245 | 13036 |
| | 5 | | 1 | 0 | 3211 | 3142 | 0 | 0 | 0 | 350 | 9453 | 245 | 16401 |
| | 10 | | 1 | 0 | 3211 | 3142 | 0 | 0 | 0 | 350 | 20845 | 245 | 27793 |
| | 50 | | 1 | 0 | 3211 | 3142 | 0 | 0 | 0 | 350 | 42827 | 245 | 49775 |
| | 0 | | 0 | 0 | 4036 | 257 | 0 | 0 | 0 | 8104 | 121 | 4442 | 16960 |
| | 3.5 | | 0 | 0 | 252 | 100 | 0 | 0 | 0 | 4393 | 6960 | 4251 | 15955 |
| | 5 | | 0 | 0 | 252 | 100 | 0 | 0 | 0 | 4393 | 9881 | 4251 | 18876 |
| | 10 | | 0 | 0 | 252 | 100 | 0 | 0 | 0 | 4393 | 21632 | 4251 | 30627 |
| | 50 | | 0 | 0 | 252 | 100 | 0 | 0 | 0 | 4393 | 44216 | 4251 | 53211 |

**Table S10. Measured and simulated SENSOR-MUX*-AHL dynamics.** Measured mean sfGFP fluorescence was calculated by subtracting mean autofluorescence (238 MEFL, which differed from previous measurements due to small cytometer fluctuations) from mean cellular fluorescence, and simulated mean cellular fluorescence was similarly calculated by adding mean autofluorescence to simulated sfGFP fluorescence. Inducer concentrations: 0 (0); 20 ng/mL aTc, 0.3 mM IPTG, and 100 µM DAPG (1).

| aTc | IPTG | DAPG | Time (hours) | Output | Measured mean fluor. (MEFL) | Simulated mean fluor. (MEFL) | Measured mean sfGFP fluor. (MEFL) | Simulated mean sfGFP fluor. (MEFL) |
|---|---|---|---|---|---|---|---|---|
| 0 | 1 | 1 | 0 | $OUT_{DAPG\ sensor}$ | 305.71 | 237.90 | 67.81 | 0.00 |
| 0 | 1 | 1 | 0.5 | $OUT_{DAPG\ sensor}$ | 654.70 | 5170.17 | 416.80 | 4932.27 |
| 0 | 1 | 1 | 1 | $OUT_{DAPG\ sensor}$ | 993.36 | 5170.17 | 755.46 | 4932.27 |
| 0 | 1 | 1 | 1.5 | $OUT_{DAPG\ sensor}$ | 1129.32 | 5170.17 | 891.42 | 4932.27 |
| 0 | 1 | 1 | 2 | $OUT_{DAPG\ sensor}$ | 1342.30 | 5170.17 | 1104.40 | 4932.27 |
| 0 | 1 | 1 | 2.5 | $OUT_{DAPG\ sensor}$ | 1387.53 | 5170.17 | 1149.63 | 4932.27 |
| 0 | 1 | 1 | 3 | $OUT_{DAPG\ sensor}$ | 1537.55 | 5170.17 | 1299.65 | 4932.27 |
| 0 | 1 | 1 | 4 | $OUT_{DAPG\ sensor}$ | 1733.83 | 5170.17 | 1495.93 | 4932.27 |
| 0 | 1 | 1 | 5 | $OUT_{DAPG\ sensor}$ | 1903.48 | 5170.17 | 1665.58 | 4932.27 |
| 0 | 1 | 1 | 6 | $OUT_{DAPG\ sensor}$ | 2263.98 | 5170.17 | 2026.07 | 4932.27 |
| 0 | 1 | 1 | 7 | $OUT_{DAPG\ sensor}$ | 2498.97 | 5170.17 | 2261.07 | 4932.27 |
| 0 | 1 | 1 | 8 | $OUT_{DAPG\ sensor}$ | 2745.03 | 5170.17 | 2507.13 | 4932.27 |
| 0 | 1 | 1 | 9 | $OUT_{DAPG\ sensor}$ | 2809.13 | 5170.17 | 2571.23 | 4932.27 |
| 0 | 1 | 1 | 10 | $OUT_{DAPG\ sensor}$ | 2991.15 | 5170.17 | 2753.25 | 4932.27 |
| 0 | 1 | 1 | 0 | $OUT_{NOT4}$ | 535.54 | 572.97 | 297.64 | 335.07 |
| 0 | 1 | 1 | 0.5 | $OUT_{NOT4}$ | 501.82 | 254.71 | 263.92 | 16.81 |
| 0 | 1 | 1 | 1 | $OUT_{NOT4}$ | 401.48 | 251.43 | 163.58 | 13.53 |
| 0 | 1 | 1 | 1.5 | $OUT_{NOT4}$ | 351.59 | 250.47 | 113.69 | 12.57 |
| 0 | 1 | 1 | 2 | $OUT_{NOT4}$ | 321.15 | 250.05 | 83.25 | 12.15 |
| 0 | 1 | 1 | 2.5 | $OUT_{NOT4}$ | 292.65 | 249.83 | 54.75 | 11.93 |
| 0 | 1 | 1 | 3 | $OUT_{NOT4}$ | 281.11 | 249.70 | 43.21 | 11.80 |
| 0 | 1 | 1 | 4 | $OUT_{NOT4}$ | 265.02 | 249.58 | 27.12 | 11.68 |
| 0 | 1 | 1 | 5 | $OUT_{NOT4}$ | 260.79 | 249.54 | 22.89 | 11.64 |
| 0 | 1 | 1 | 6 | $OUT_{NOT4}$ | 262.70 | 249.52 | 24.80 | 11.62 |
| 0 | 1 | 1 | 7 | $OUT_{NOT4}$ | 261.22 | 249.51 | 23.32 | 11.61 |
| 0 | 1 | 1 | 8 | $OUT_{NOT4}$ | 259.02 | 249.51 | 21.12 | 11.61 |
| 0 | 1 | 1 | 9 | $OUT_{NOT4}$ | 256.67 | 249.51 | 18.77 | 11.61 |
| 0 | 1 | 1 | 10 | $OUT_{NOT4}$ | 258.73 | 249.51 | 20.83 | 11.60 |
| 0 | 1 | 1 | 0 | $OUT_{NOT2}$ | 957.88 | 443.01 | 719.98 | 205.11 |
| 0 | 1 | 1 | 0.5 | $OUT_{NOT2}$ | 874.45 | 547.23 | 636.55 | 309.32 |
| 0 | 1 | 1 | 1 | $OUT_{NOT2}$ | 822.46 | 735.83 | 584.56 | 497.93 |
| 0 | 1 | 1 | 1.5 | $OUT_{NOT2}$ | 1313.05 | 1024.88 | 1075.15 | 786.97 |
| 0 | 1 | 1 | 2 | $OUT_{NOT2}$ | 2248.32 | 1426.98 | 2010.42 | 1189.08 |
| 0 | 1 | 1 | 2.5 | $OUT_{NOT2}$ | 3334.93 | 1921.72 | 3097.03 | 1683.82 |
| 0 | 1 | 1 | 3 | $OUT_{NOT2}$ | 4535.55 | 2450.96 | 4297.65 | 2213.06 |
| 0 | 1 | 1 | 4 | $OUT_{NOT2}$ | 6315.98 | 3350.24 | 6078.08 | 3112.34 |
| 0 | 1 | 1 | 5 | $OUT_{NOT2}$ | 7171.15 | 3874.06 | 6933.25 | 3636.16 |
| 0 | 1 | 1 | 6 | $OUT_{NOT2}$ | 8074.13 | 4120.94 | 7836.22 | 3883.04 |
| 0 | 1 | 1 | 7 | $OUT_{NOT2}$ | 8413.45 | 4228.20 | 8175.54 | 3990.30 |
| 0 | 1 | 1 | 8 | $OUT_{NOT2}$ | 8650.51 | 4273.52 | 8412.61 | 4035.62 |
| 0 | 1 | 1 | 9 | $OUT_{NOT2}$ | 8737.67 | 4292.48 | 8499.77 | 4054.58 |
| 0 | 1 | 1 | 10 | $OUT_{NOT2}$ | 8229.04 | 4300.38 | 7991.14 | 4062.48 |
| 0 | 1 | 1 | 0 | $OUT_{NOR6*}$ | 4512.96 | 1288.51 | 4275.06 | 1050.61 |
| 0 | 1 | 1 | 0.5 | $OUT_{NOR6*}$ | 4288.31 | 1214.82 | 4050.41 | 976.92 |

| | | | | | | | | |
|---|---|---|---|---|---|---|---|---|
| 0 | 1 | 1 | 1 | $OUT_{NOR6*}$ | 3795.37 | 992.11 | 3557.47 | 754.21 |
| 0 | 1 | 1 | 1.5 | $OUT_{NOR6*}$ | 3039.23 | 740.97 | 2801.33 | 503.07 |
| 0 | 1 | 1 | 2 | $OUT_{NOR6*}$ | 2221.04 | 549.61 | 1983.14 | 311.71 |
| 0 | 1 | 1 | 2.5 | $OUT_{NOR6*}$ | 1493.18 | 430.26 | 1255.28 | 192.36 |
| 0 | 1 | 1 | 3 | $OUT_{NOR6*}$ | 988.09 | 362.00 | 750.19 | 124.09 |
| 0 | 1 | 1 | 4 | $OUT_{NOR6*}$ | 509.66 | 301.70 | 271.76 | 63.80 |
| 0 | 1 | 1 | 5 | $OUT_{NOR6*}$ | 362.13 | 280.72 | 124.23 | 42.82 |
| 0 | 1 | 1 | 6 | $OUT_{NOR6*}$ | 319.47 | 272.31 | 81.57 | 34.41 |
| 0 | 1 | 1 | 7 | $OUT_{NOR6*}$ | 301.53 | 268.59 | 63.63 | 30.69 |
| 0 | 1 | 1 | 8 | $OUT_{NOR6*}$ | 291.31 | 266.85 | 53.41 | 28.95 |
| 0 | 1 | 1 | 9 | $OUT_{NOR6*}$ | 300.80 | 266.03 | 62.90 | 28.13 |
| 0 | 1 | 1 | 10 | $OUT_{NOR6*}$ | 295.10 | 265.63 | 57.19 | 27.73 |
| 0 | 1 | 1 | 0 | $OUT_{NOR3}$ | 429.07 | 277.54 | 191.17 | 39.64 |
| 0 | 1 | 1 | 0.5 | $OUT_{NOR3}$ | 424.76 | 277.60 | 186.86 | 39.70 |
| 0 | 1 | 1 | 1 | $OUT_{NOR3}$ | 380.46 | 280.12 | 142.56 | 42.22 |
| 0 | 1 | 1 | 1.5 | $OUT_{NOR3}$ | 358.44 | 287.07 | 120.54 | 49.17 |
| 0 | 1 | 1 | 2 | $OUT_{NOR3}$ | 357.74 | 300.26 | 119.84 | 62.36 |
| 0 | 1 | 1 | 2.5 | $OUT_{NOR3}$ | 332.22 | 321.98 | 94.32 | 84.08 |
| 0 | 1 | 1 | 3 | $OUT_{NOR3}$ | 372.21 | 354.91 | 134.30 | 117.01 |
| 0 | 1 | 1 | 4 | $OUT_{NOR3}$ | 511.90 | 461.52 | 274.00 | 223.62 |
| 0 | 1 | 1 | 5 | $OUT_{NOR3}$ | 837.09 | 610.84 | 599.19 | 372.94 |
| 0 | 1 | 1 | 6 | $OUT_{NOR3}$ | 1272.77 | 757.57 | 1034.86 | 519.67 |
| 0 | 1 | 1 | 7 | $OUT_{NOR3}$ | 1538.82 | 864.50 | 1300.91 | 626.59 |
| 0 | 1 | 1 | 8 | $OUT_{NOR3}$ | 1783.15 | 928.35 | 1545.25 | 690.45 |
| 0 | 1 | 1 | 9 | $OUT_{NOR3}$ | 1842.91 | 962.46 | 1605.01 | 724.56 |
| 0 | 1 | 1 | 10 | $OUT_{NOR3}$ | 1940.29 | 979.66 | 1702.39 | 741.76 |

**Table S11. Measured and simulated AHL-DEMUX dynamics.** Fluorescence signals were calculated as described for **Table S10**. Inducer concentrations: 0 (0); 3.5 nM AHL, 100 µM DAPG (1).

| AHL | DAPG | Time (hours) | Output | Measured mean fluor. (MEFL) | Simulated mean fluor. (MEFL) | Measured mean sfGFP fluor. (MEFL) | Simulated mean sfGFP fluor. (MEFL) |
|---|---|---|---|---|---|---|---|
| 1 | 1 | 0 | $OUT_{AHL\ sensor}$ | 309.53 | 237.90 | 71.63 | 0.00 |
| 1 | 1 | 0.5 | $OUT_{AHL\ sensor}$ | 3153.66 | 1278.80 | 2915.76 | 1040.90 |
| 1 | 1 | 1 | $OUT_{AHL\ sensor}$ | 4566.63 | 1278.80 | 4328.73 | 1040.90 |
| 1 | 1 | 1.5 | $OUT_{AHL\ sensor}$ | 5512.31 | 1278.80 | 5274.41 | 1040.90 |
| 1 | 1 | 2 | $OUT_{AHL\ sensor}$ | 5796.93 | 1278.80 | 5559.03 | 1040.90 |
| 1 | 1 | 2.5 | $OUT_{AHL\ sensor}$ | 6710.26 | 1278.80 | 6472.36 | 1040.90 |
| 1 | 1 | 3 | $OUT_{AHL\ sensor}$ | 6670.47 | 1278.80 | 6432.57 | 1040.90 |
| 1 | 1 | 4 | $OUT_{AHL\ sensor}$ | 6625.50 | 1278.80 | 6387.60 | 1040.90 |
| 1 | 1 | 5 | $OUT_{AHL\ sensor}$ | 6633.93 | 1278.80 | 6396.03 | 1040.90 |
| 1 | 1 | 6 | $OUT_{AHL\ sensor}$ | 6763.30 | 1278.80 | 6525.40 | 1040.90 |
| 1 | 1 | 7 | $OUT_{AHL\ sensor}$ | 6815.89 | 1278.80 | 6577.99 | 1040.90 |
| 1 | 1 | 8 | $OUT_{AHL\ sensor}$ | 6842.15 | 1278.80 | 6604.25 | 1040.90 |
| 1 | 1 | 9 | $OUT_{AHL\ sensor}$ | 6765.62 | 1278.80 | 6527.72 | 1040.90 |
| 1 | 1 | 10 | $OUT_{AHL\ sensor}$ | 7223.53 | 1278.80 | 6985.63 | 1040.90 |
| 1 | 1 | 0 | $OUT_{NOT8}$ | 2918.89 | 2787.15 | 2680.99 | 2549.25 |
| 1 | 1 | 0.5 | $OUT_{NOT8}$ | 2438.55 | 795.41 | 2200.64 | 557.51 |
| 1 | 1 | 1 | $OUT_{NOT8}$ | 1751.90 | 393.98 | 1514.00 | 156.08 |
| 1 | 1 | 1.5 | $OUT_{NOT8}$ | 1300.56 | 326.13 | 1062.66 | 88.23 |
| 1 | 1 | 2 | $OUT_{NOT8}$ | 969.34 | 304.02 | 731.44 | 66.12 |
| 1 | 1 | 2.5 | $OUT_{NOT8}$ | 742.61 | 294.31 | 504.71 | 56.41 |
| 1 | 1 | 3 | $OUT_{NOT8}$ | 599.54 | 289.34 | 361.64 | 51.44 |
| 1 | 1 | 4 | $OUT_{NOT8}$ | 409.34 | 284.91 | 171.44 | 47.01 |
| 1 | 1 | 5 | $OUT_{NOT8}$ | 351.04 | 283.30 | 113.14 | 45.40 |
| 1 | 1 | 6 | $OUT_{NOT8}$ | 328.78 | 282.66 | 90.88 | 44.76 |
| 1 | 1 | 7 | $OUT_{NOT8}$ | 323.18 | 282.40 | 85.28 | 44.50 |
| 1 | 1 | 8 | $OUT_{NOT8}$ | 319.72 | 282.30 | 81.82 | 44.40 |
| 1 | 1 | 9 | $OUT_{NOT8}$ | 328.93 | 282.25 | 91.03 | 44.35 |
| 1 | 1 | 10 | $OUT_{NOT8}$ | 345.56 | 282.23 | 107.66 | 44.33 |
| 1 | 1 | 0 | $OUT_{NOR7}$ | 325.55 | 259.70 | 87.65 | 21.80 |
| 1 | 1 | 0.5 | $OUT_{NOR7}$ | 350.43 | 263.34 | 112.53 | 25.44 |
| 1 | 1 | 1 | $OUT_{NOR7}$ | 343.14 | 279.01 | 105.24 | 41.11 |
| 1 | 1 | 1.5 | $OUT_{NOR7}$ | 376.71 | 314.54 | 138.81 | 76.64 |
| 1 | 1 | 2 | $OUT_{NOR7}$ | 444.31 | 389.62 | 206.41 | 151.72 |
| 1 | 1 | 2.5 | $OUT_{NOR7}$ | 554.33 | 536.83 | 316.42 | 298.93 |
| 1 | 1 | 3 | $OUT_{NOR7}$ | 740.52 | 793.42 | 502.62 | 555.52 |
| 1 | 1 | 4 | $OUT_{NOR7}$ | 1207.06 | 1614.85 | 969.16 | 1376.95 |
| 1 | 1 | 5 | $OUT_{NOR7}$ | 1485.37 | 2370.68 | 1247.47 | 2132.77 |
| 1 | 1 | 6 | $OUT_{NOR7}$ | 1667.97 | 2759.38 | 1430.06 | 2521.48 |
| 1 | 1 | 7 | $OUT_{NOR7}$ | 1694.64 | 2921.28 | 1456.74 | 2683.38 |
| 1 | 1 | 8 | $OUT_{NOR7}$ | 1673.79 | 2987.04 | 1435.89 | 2749.13 |
| 1 | 1 | 9 | $OUT_{NOR7}$ | 1646.26 | 3014.09 | 1408.36 | 2776.19 |
| 1 | 1 | 10 | $OUT_{NOR7}$ | 1773.31 | 3025.35 | 1535.41 | 2787.45 |

**Table S12. Constitutive promoters used to drive *phlF* in the DAPG sensor**. We initially used an insulated BBa_J23101 (http://parts.igem.org/) to mimic a previously published design (*40*) (**Methods**). We found, however, that the $P_{PhlF}$ promoter remained repressed in the presence of DAPG. To address this, we replaced BBa_J23101 with the weaker BBa_J23114 (same insulator), which recovered DAPG-dependent $P_{PhlF}$ activation.

| Promoter | Sequence |
| --- | --- |
| Insulated BBa_J23114 (functional) | TGTAGAGTTATCCGCCTACGGCGCCGTCGTATCGG TAATCCGTACGGGAATCGAAACGACGTCTACGAGC TTTATGGCTAGCTCAGTCCTAGGTACAATGCTAGC |
| Insulated BBa_J23101 (nonfunctional) | TGTAGAGTTATCCGCCTACGGCGCCGTCGTATCGG TAATCCGTACGGGAATCGAAACGACGTCTACGAGC TTTACAGCTAGCTCAGTCCTAGGTATTATGCTAGC |

**Table S13. SENSOR-MUX*-AHL connectivity matrix, $M_{SENSOR\text{-}MUX*\text{-}AHL}$.** To simulate dynamical circuit behavior, a connectivity matrix was used to specify connections between gates and to reporter genes (**Supplementary Text**). Here, the matrix for SENSOR-MUX*-AHL is shown. Abbreviations: A, aTc sensor; I, IPTG sensor; Q, AHL sensor; D, DAPG sensor.

| | C | | | | | | | | | | G | | | | | | | | | | | | | |
|---|---|---|---|---|---|---|---|---|---|---|---|---|---|---|---|---|---|---|---|---|---|---|---|---|
| | 1 | 2 | 3 | 4 | 5 | 6 | 7 | 8 | 9 | 6* | A | I | Q | D | 1 | 2 | 3 | 4 | 5 | 6 | 7 | 8 | 9 | 6* |
| $OUT_{aTc\ sensor}$ | 1 | 0 | 0 | 0 | 0 | 0 | 0 | 0 | 0 | 0 | 1 | 0 | 0 | 0 | 0 | 0 | 0 | 0 | 0 | 0 | 0 | 0 | 0 | 0 |
| $OUT_{IPTG\ sensor}$ | 0 | 0 | 0 | 0 | 0 | 0 | 0 | 0 | 1 | 0 | 0 | 1 | 0 | 0 | 0 | 0 | 0 | 0 | 0 | 0 | 0 | 0 | 0 | 0 |
| $OUT_{AHL\ sensor}$ | 0 | 0 | 0 | 0 | 0 | 0 | 0 | 0 | 0 | 0 | 0 | 0 | 0 | 0 | 0 | 0 | 0 | 0 | 0 | 0 | 0 | 0 | 0 | 0 |
| $OUT_{DAPG\ sensor}$ | 0 | 0 | 0 | 1 | 0 | 0 | 0 | 0 | 0 | 0 | 0 | 0 | 0 | 1 | 0 | 0 | 0 | 0 | 0 | 0 | 0 | 0 | 0 | 0 |
| $OUT_{GATE1}$ | 0 | 0 | 0 | 0 | 1 | 0 | 0 | 0 | 0 | 0 | 0 | 0 | 0 | 0 | 1 | 0 | 0 | 0 | 0 | 0 | 0 | 0 | 0 | 0 |
| $OUT_{GATE2}$ | 0 | 0 | 0 | 0 | 0 | 0 | 0 | 0 | 0 | 1 | 0 | 0 | 0 | 0 | 0 | 1 | 0 | 0 | 0 | 0 | 0 | 0 | 0 | 0 |
| $OUT_{GATE3}$ | 0 | 0 | 0 | 0 | 0 | 0 | 0 | 0 | 0 | 0 | 0 | 0 | 0 | 0 | 0 | 0 | 1 | 0 | 0 | 0 | 0 | 0 | 0 | 0 |
| $OUT_{GATE4}$ | 0 | 1 | 0 | 0 | 1 | 0 | 0 | 0 | 0 | 0 | 0 | 0 | 0 | 0 | 0 | 0 | 0 | 1 | 0 | 0 | 0 | 0 | 0 | 0 |
| $OUT_{GATE5}$ | 0 | 0 | 1 | 0 | 0 | 0 | 0 | 0 | 0 | 0 | 0 | 0 | 0 | 0 | 0 | 0 | 0 | 0 | 1 | 0 | 0 | 0 | 0 | 0 |
| $OUT_{GATE6}$ | 0 | 0 | 0 | 0 | 0 | 0 | 0 | 0 | 0 | 0 | 0 | 0 | 0 | 0 | 0 | 0 | 0 | 0 | 0 | 0 | 0 | 0 | 0 | 0 |
| $OUT_{GATE7}$ | 0 | 0 | 0 | 0 | 0 | 0 | 0 | 0 | 0 | 0 | 0 | 0 | 0 | 0 | 0 | 0 | 0 | 0 | 0 | 0 | 0 | 0 | 0 | 0 |
| $OUT_{GATE8}$ | 0 | 0 | 0 | 0 | 0 | 0 | 0 | 0 | 0 | 0 | 0 | 0 | 0 | 0 | 0 | 0 | 0 | 0 | 0 | 0 | 0 | 0 | 0 | 0 |
| $OUT_{GATE9}$ | 0 | 0 | 0 | 0 | 0 | 0 | 0 | 0 | 0 | 1 | 0 | 0 | 0 | 0 | 0 | 0 | 0 | 0 | 0 | 0 | 0 | 0 | 1 | 0 |
| $OUT_{GATE6*}$ | 0 | 0 | 1 | 0 | 0 | 0 | 0 | 0 | 0 | 0 | 0 | 0 | 0 | 0 | 0 | 0 | 0 | 0 | 0 | 0 | 0 | 0 | 0 | 1 |

**Table S14. AHL-DEMUX connectivity matrix, $M_{AHL\text{-}DEMUX}$.** To simulate dynamical circuit behavior, a connectivity matrix was used to specify connections between gates and to reporter genes (**Supplementary Text**). Here, the matrix for the AHL-DEMUX is shown. Abbreviations: A, aTc sensor; I, IPTG sensor; Q, AHL sensor; D, DAPG sensor.

| | C | | | | | | | | | | G | | | | | | | | | | | | | |
|---|---|---|---|---|---|---|---|---|---|---|---|---|---|---|---|---|---|---|---|---|---|---|---|---|
| | 1 | 2 | 3 | 4 | 5 | 6 | 7 | 8 | 9 | 6* | A | I | Q | D | 1 | 2 | 3 | 4 | 5 | 6 | 7 | 8 | 9 | 6* |
| $\mathrm{OUT}_{\mathrm{aTc\ sensor}}$ | 0 | 0 | 0 | 0 | 0 | 0 | 0 | 0 | 0 | 0 | 0 | 0 | 0 | 0 | 0 | 0 | 0 | 0 | 0 | 0 | 0 | 0 | 0 | 0 |
| $\mathrm{OUT}_{\mathrm{IPTG\ sensor}}$ | 0 | 0 | 0 | 0 | 0 | 0 | 0 | 0 | 0 | 0 | 0 | 0 | 0 | 0 | 0 | 0 | 0 | 0 | 0 | 0 | 0 | 0 | 0 | 0 |
| $\mathrm{OUT}_{\mathrm{AHL\ sensor}}$ | 0 | 0 | 0 | 0 | 0 | 0 | 0 | 1 | 0 | 0 | 0 | 0 | 1 | 0 | 0 | 0 | 0 | 0 | 0 | 0 | 0 | 0 | 0 | 0 |
| $\mathrm{OUT}_{\mathrm{DAPG\ sensor}}$ | 0 | 0 | 1 | 0 | 0 | 0 | 0 | 0 | 0 | 0 | 0 | 0 | 0 | 1 | 0 | 0 | 0 | 0 | 0 | 0 | 0 | 0 | 0 | 0 |
| $\mathrm{OUT}_{\mathrm{GATE1}}$ | 0 | 0 | 0 | 0 | 0 | 0 | 0 | 0 | 0 | 0 | 0 | 0 | 0 | 0 | 0 | 0 | 0 | 0 | 0 | 0 | 0 | 0 | 0 | 0 |
| $\mathrm{OUT}_{\mathrm{GATE2}}$ | 0 | 0 | 0 | 0 | 0 | 0 | 0 | 0 | 0 | 0 | 0 | 0 | 0 | 0 | 0 | 1 | 0 | 0 | 0 | 0 | 0 | 0 | 0 | 0 |
| $\mathrm{OUT}_{\mathrm{GATE3}}$ | 0 | 0 | 0 | 0 | 0 | 0 | 1 | 0 | 1 | 0 | 0 | 0 | 0 | 0 | 0 | 0 | 1 | 0 | 0 | 0 | 0 | 0 | 0 | 0 |
| $\mathrm{OUT}_{\mathrm{GATE4}}$ | 0 | 0 | 0 | 0 | 0 | 0 | 0 | 0 | 0 | 0 | 0 | 0 | 0 | 0 | 0 | 0 | 0 | 0 | 0 | 0 | 0 | 0 | 0 | 0 |
| $\mathrm{OUT}_{\mathrm{GATE5}}$ | 0 | 0 | 0 | 0 | 0 | 0 | 0 | 0 | 0 | 0 | 0 | 0 | 0 | 0 | 0 | 0 | 0 | 0 | 0 | 0 | 0 | 0 | 0 | 0 |
| $\mathrm{OUT}_{\mathrm{GATE6}}$ | 0 | 0 | 0 | 0 | 0 | 0 | 0 | 0 | 0 | 0 | 0 | 0 | 0 | 0 | 0 | 0 | 0 | 0 | 0 | 0 | 0 | 0 | 0 | 0 |
| $\mathrm{OUT}_{\mathrm{GATE7}}$ | 0 | 0 | 0 | 0 | 0 | 0 | 0 | 0 | 0 | 0 | 0 | 0 | 0 | 0 | 0 | 0 | 0 | 0 | 0 | 0 | 1 | 0 | 0 | 0 |
| $\mathrm{OUT}_{\mathrm{GATE8}}$ | 0 | 1 | 0 | 0 | 0 | 0 | 1 | 0 | 0 | 0 | 0 | 0 | 0 | 0 | 0 | 0 | 0 | 0 | 0 | 0 | 0 | 1 | 0 | 0 |
| $\mathrm{OUT}_{\mathrm{GATE9}}$ | 0 | 1 | 0 | 0 | 0 | 0 | 0 | 0 | 0 | 0 | 0 | 0 | 0 | 0 | 0 | 0 | 0 | 0 | 0 | 0 | 0 | 0 | 1 | 0 |
| $\mathrm{OUT}_{\mathrm{GATE6*}}$ | 0 | 0 | 0 | 0 | 0 | 0 | 0 | 0 | 0 | 0 | 0 | 0 | 0 | 0 | 0 | 0 | 0 | 0 | 0 | 0 | 0 | 0 | 0 | 0 |

**Table S15. Sequences of parts used in this study**. Terminator sequences can be found in (*41*). For compound parts, promoter = green, RBS = purple, gene = blue, terminator = red.

| Label | Type | Sequence | Reference |
|---|---|---|---|
| P1 | promoter | ACGTAGGGTAAGGAAGCGTAGTCGGGTCCGTTAAGGTTCGGTACGGACGGACCGCGTCGGTAGAGACTTCGACAATCGATCATGCGATTGGTATAATAGATTCAT | This work |
| P2 | promoter | CGTTTTCGGGACGGATAAGGATTTCCTCCCGCGTAACCGTTTAATAACGGCGACCGTTACGCGAAGACTCGACAACAACGTGACAACACGGTATAATAGATTCAT | This work |
| P3 | promoter | GTTACCTTCCCGGAGGTAGCCGCGTTCCGCCCGAGTCGGACGAGAACCGGGAGTCTTCGAAGGCTCTATCGACAATGTTGTGTTACGTTGGTATAATAGATTCAT | This work |
| P4 | promoter | GGTTCCCTTTTCCTCGTAAGGGCCGGGAACCGATTACGTCTCGGAGGGCCTCCGAGTCCTCGCGGTCTTCGACATCGATAATGACACGCGGTATAATAGATTCAT | This work |
| P5 | promoter | GTTACGCGAAGGTAGGGAGAAGCGGTTTCCGCTTTAACCCGCCGGGACGTCTCGTAAACGTCGCGGGTTCGACATACGACACACGCAATGGTATAATAGATTCAT | This work |
| P6 | promoter | TATCGCTTCCGCGTTCCGTCTACTCCCTATTACGCTCCGGAATCTCCGACGCCTACGGCCGACGTCGGATTGACTATTGATTACAACGTGTCGGATAATGGTTGC | This work |
| P7 | promoter | CTCCTTCGTAGATATCGTCCGGGTCCGCTTAAGCTCTTACGTTCGTAGGAGTAAGACGACCCTCGGTATCGACATTGTGCGCGATCGACGGTATAATAGATTCAT | This work |
| P8 | promoter | CCTCTCCTTCGTCCCGAGAGTCCCGTTCGCGAAACGCCTCGATAACGAAAACGGAGCGAGGACTTCCCTCGACACGTGCGTTCACGTGTGGTATAATAGATTCAT | This work |
| P9 | promoter | TTTATTTATCGTCCGCCGGCCGACGGACTTCGTATTTCGGAGCTACTCGGGCGTATACGTATCGGGCCTCGACATGTCGTGCGTGTATTGGTATAATAGATTCAT | This work |
| P6* | promoter | TCGCTTCCGCGTTCCGTCTACTCCCTATTACGCTCCGGAATCTCCGACGCCTACGGCCGACGTCGGAAAAAAATTTATTTGCTTTGATTACAACGTGTCTGGTATAATGTGTGGAT | This work |
| $P_R$ | promoter | TAACACCGTGCGTGTTGACTATTTTACCTCTGGCGGTGATAATGGTTGC | (*48*) |
| $P_{tet}$ | promoter | GGTAGTAGCTCGGGAGTCCTTTTCCGGATTTCTATCCGGGCCTTCGACGGATCCCTATCAGTGATAGAGATTGACATCCCTATCAGTGATAGAGATACTGAGCAC | Modified from (*49*) |
| $P_{tac}$ | promoter | TTTCGATAAACTTCGGGACGGGATAGCTCTCCTACCTTTCTCTTAAGCTCGAATACCTCGAGGCCCGCCCTTGACAATTAATCATCGGCTCGTATAATGTGTGGAATTGTGAGCGGATAACAATTTCACACA | Modified from (*50*) |
| $P_{PhlF}$ | promoter | CGACGTACGGTGGAATCTGATTCGTTACCAATTGACATGATACGAAACGTACCGTATCGTTAAGGT | (*40*) |
| $P_{lux}$ | promoter | TACAATTGTTTAACATAAGTACCTGTAGGATCGTACAGGTTTACTATTTTACCTCTGGCGGTGATAATTCTTGCAACAAACAATAGGTAAGGCGTTACCCAAC | D49 from (*51*) |
| S1 | sgRNA | TCGACAATCGATCATGCGATGTTTTAGAGCTAGAAATAGCAAGTTAAAATAAGGCTAGTCCGTTATCAACTTGAAAAAGTGGCACCGAGTCGGTGCTTTTTT | This work |
| S2 | sgRNA | TCGACAACAACGTGACAACAGTTTTAGAGCTAGAAATAGCAAGTTAAAATAAGGCTAGTCCGTTATCAACTTGAAAAAGTGGCACCGAGTCGGTGCTTTTTT | This work |
| S3 | sgRNA | TCGACAATGTTGTGTTACGTGTTTTAGAGCTAGAAATAGCAAGTTAAAATAAGGCTAGTCCGTTATCAACTTGAAAAAGTGGCACCGAGTCGGTGCTTTTTT | This work |

| | | | |
|---|---|---|---|
| S4 | sgRNA | TCGACATCGATAATGACACGGTTTTAGAGCTAGAAATAGCAAGTT<br>AAAATAAGGCTAGTCCGTTATCAACTTGAAAAAGTGGCACCGAGT<br>CGGTGCTTTTTT | This work |
| S5 | sgRNA | TCGACATACGACACACGCAAGTTTTAGAGCTAGAAATAGCAAGTT<br>AAAATAAGGCTAGTCCGTTATCAACTTGAAAAAGTGGCACCGAGT<br>CGGTGCTTTTTT | This work |
| S6 | sgRNA | GACTATTGATTACAACGTGTGTTTTAGAGCTAGAAATAGCAAGTT<br>AAAATAAGGCTAGTCCGTTATCAACTTGAAAAAGTGGCACCGAGT<br>CGGTGCTTTTTT | This work |
| S7 | sgRNA | TCGACATTGTGCGCGATCGAGTTTTAGAGCTAGAAATAGCAAGTT<br>AAAATAAGGCTAGTCCGTTATCAACTTGAAAAAGTGGCACCGAGT<br>CGGTGCTTTTTT | This work |
| S8 | sgRNA | TCGACACGTGCGTTCACGTGGTTTTAGAGCTAGAAATAGCAAGTT<br>AAAATAAGGCTAGTCCGTTATCAACTTGAAAAAGTGGCACCGAGT<br>CGGTGCTTTTTT | This work |
| S9 | sgRNA | TCGACATGTCGTGCGTGTATGTTTTAGAGCTAGAAATAGCAAGTT<br>AAAATAAGGCTAGTCCGTTATCAACTTGAAAAAGTGGCACCGAGT<br>CGGTGCTTTTTT | This work |
| S6* | sgRNA | TGCTTTGATTACAACGTGTCGTTTTAGAGCTAGAAATAGCAAGTT<br>AAAATAAGGCTAGTCCGTTATCAACTTGAAAAAGTGGCACCGAGT<br>CGGTGCTTTTTT | This work |
| *tetR* | transcription unit | GGTACCCGGGGATCCTCTAGAGTCGACCTGCAGGCATGCAAGCTT<br>TTTACGGCTAGCTCAGTCCTAGGTATAGTGCTAGCCCAGCCAGAG<br>AAACACTCTTTAACAGGGGGTAGTATGATGTCTCGTTTAGATAAA<br>AGTAAAGTGATTAACAGCGCATTAGAGCTGCTTAATGAGGTCGGA<br>ATCGAAGGTTTAACAACCCGTAAACTCGCCCAGAAGCTAGGTGTA<br>GAGCAGCCTACATTGTATTGGCATGTAAAAAATAAGCGGGCTTTG<br>CTCGACGCCTTAGCCATTGAGATGTTAGATAGGCACCATACTCAC<br>TTTTGCCCTTTAGAAGGGGAAAGCTGGCAAGATTTTTTACGTAAT<br>AACGCTAAAAGTTTTAGATGTGCTTTACTAAGTCATCGCGATGGA<br>GCAAAAGTACATTTAGGTACACGGCCTACAGAAAAACAGTATGAA<br>ACTCTCGAAAATCAATTAGCCTTTTTATGCCAACAAGGTTTTTCA<br>CTAGAGAATGCATTATATGCACTCAGCGCAGTGGGGCATTTTACT<br>TTAGGTTGCGTATTGGAAGATCAAGAGCATCAAGTCGCTAAAGAA<br>GAAAGGGAAACACCTACTACTGATAGTATGCCGCCATTATTACGA<br>CAAGCTATCGAATTATTTGATCACCAAGGTGCAGAGCCAGCCTTC<br>TTATTCGGCCTTGAATTGATCATATGCGGATTAGAAAAACAACTT<br>AAATGTGAAAGTGGGTCTTAAGGCATCAAATAAAACGAAAGGCTC<br>AGTCGAAAGACTGGGCCTTTCGTTTTATCTGTTGTTTGTCGGTGA<br>ACGCTCTCCTGAGTAGGACAAATCCGCCGCCCTAGA | (*39*) |
| *lacI* | transcription unit | CGAAGCGGCATGCATTTACGTTGACACCATCGAATGGTGCAAAAC<br>CTTTCGCGGTATGGCATGATAGCGCCCGGAAGAGAGTCAATTCAG<br>GGTGGTGAATGTGAAACCAGTAACGTTATACGATGTCGCAGAGTA<br>TGCCGGTGTCTCTTATCAGACCGTTTCCCGCGTGGTGAACCAGGC<br>CAGCCACGTTTCTGCGAAAACGCGGGAAAAAGTGGAAGCGGCGAT<br>GGCGGAGCTGAATTACATTCCCAACCGCGTGGCACAACAACTGGC<br>GGGCAAACAGTCGTTGCTGATTGGCGTTGCCACCTCCAGTCTGGC<br>CCTGCACGCGCCGTCGCAAATTGTCGCGGCGATTAAATCTCGCGC<br>CGATCAACTGGGTGCCAGCGTGGTGGTGTCGATGGTAGAACGAAG<br>CGGCGTCGAAGCCTGTAAAGCGGCGGTGCACAATCTTCTCGCGCA<br>ACGCGTCAGTGGGCTGATCATTAACTATCCGCTGGATGACCAGGA<br>TGCCATTGCTGTGGAAGCTGCCTGCACTAATGTTCCGGCGTTATT<br>TCTTGATGTCTCTGACCAGACACCCATCAACAGTATTATTTTCTC<br>CCATGAAGACGGTACGCGACTGGGCGTGGAGCATCTGGTCGCATT<br>GGGTCACCAGCAAATCGCGCTGTTAGCGGGCCCATTAAGTTCTGT<br>CTCGGCGCGTCTGCGTCTGGCTGGCTGGCATAAATATCTCACTCG<br>CAATCAAATTCAGCCGATAGCGGAACGGGAAGGCGACTGGAGTGC | Modified from (*39*) |

| | | | |
|---|---|---|---|
| | | CATGTCCGGTTTTCAACAAACCATGCAAATGCTGAATGAGGGCAT<br>CGTTCCCACTGCGATGCTGGTTGCCAACGATCAGATGGCGCTGGG<br>CGCAATGCGCGCCATTACCGAGTCCGGGCTGCGCGTTGGTGCGGA<br>TATCTCGGTAGTGGGATACGACGATACCGAAGACAGCTCATGTTA<br>TATCCCGCCGTTAACCACCATCAAACAGGATTTTCGCCTGCTGGG<br>GCAAACCAGCGTGGACCGCTTGCTGCAACTCTCTCAGGGCCAGGC<br>GGTGAAGGGCAATCAGCTGTTGCCCGTCTCACTGGTGAAAAGAAA<br>AACCACCCTGGCGCCCAATACGCAAACCGCCTCTCCCCGCGCGTT<br>GGCCGATTCATTAATGCAGCTGGCACGACAGGTTTCCCGACTGGA<br>AAGCGGGCAGTGACTCGGTACCAAATTCCAGAAAAGACACCCGAA<br>AGGGTGTTTTTTCGTTTTGGTCC | |
| *phlF* | transcription unit | TGTAGAGTTATCCGCCTACGGCGCCGTCGTATCGGTAATCCGTAC<br>GGGAATCGAAACGACGTCTACGAGCTTTATGGCTAGCTCAGTCCT<br>AGGTACAATGCTAGCCTGAAGTACGTCTGAGCGTGATACCCGCTC<br>ACTGAAGATGGCCCGGTAGGGCCGAAACGTACCTCTACAAATAAT<br>TTTGTTTAACTATGGACTATGTTTGAAAGGGAGAAATACTAGATG<br>GCACGTACCCCGAGCCGTAGCAGCATTGGTAGCCTGCGTAGTCCG<br>CATACCCATAAAGCAATTCTGACCAGCACCATTGAAATCCTGAAA<br>GAATGTGGTTATAGCGGTCTGAGCATTGAAAGCGTTGCACGTCGT<br>GCCGGTGCAAGCAAACCGACCATTTATCGTTGGTGGACCAATAAA<br>GCAGCACTGATTGCCGAAGTGTATGAAAATGAAAGCGAACAGGTG<br>CGTAAATTTCCGGATCTGGGTAGCTTTAAAGCCGATCTGGATTTT<br>CTGCTGCGTAATCTGTGGAAAGTTTGGCGTGAAACCATTTGTGGT<br>GAAGCATTTCGTTGTGTTATTGCAGAAGCACAGCTGGACCCTGCA<br>ACCCTGACCCAGCTGAAAGATCAGTTTATGGAACGTCGTCGTGAG<br>ATGCCGAAAAAACTGGTTGAAAATGCCATTAGCAATGGTGAACTG<br>CCGAAAGATACCAATCGTGAACTGCTGCTGGATATGATTTTTGGT<br>TTTTGTTGGTATCGCCTGCTGACCGAACAGCTGACCGTTGAACAG<br>GATATTGAAGAATTTACCTTCCTGCTGATTAATGGTGTTTGTCCG<br>GGTACACAGCGTTAATAAGGTTGAAAAATAAAAACGGCGCTAAAA<br>AGCGCCGTTTTTTTTGACGGTGGTA | Modified from (*40*) |
| *luxI* | transcription unit | GTTACCTTCCCGGAGGTAGCCGCGTTCCGCCCGAGTCGGACGAGA<br>ACCGGGAGTCTTCGAAGGCTCTATCGACAATGTTGTGTTACGTTG<br>GTATAATAGATTCATCGCTGATAGTGCTAGTGTAGATCGCTACTA<br>GAGTCACACAGGAAACCTACTAGATGACTATAATGATAAAAAAAT<br>CGGATTTTTTGGCAATTCCATCGGAGGAGTATAAAGGTATTCTAA<br>GTCTTCGTTATCAAGTGTTTAAGCAAAGACTTGAGTGGGACTTAG<br>TTGTAGAAAATAACCTTGAATCAGATGAGTATGATAACTCAAATG<br>CAGAATATATTTATGCTTGTGATGATACTGAAAATGTAAGTGGAT<br>GCTGGCGTTTATTACCTACAACAGGTGATTATATGCTGAAAAGTG<br>TTTTTCCTGAATTGCTTGGTCAACAGAGTGCTCCCAAAGATCCTA<br>ATATAGTCGAATTAAGTCGTTTTGCTGTAGGTAAAAATAGCTCAA<br>AGATAAATAACTCTGCTAGTGAAATTACAATGAAACTATTTGAAG<br>CTATATATAAACACGCTGTTAGTCAAGGTATTACAGAATATGTAA<br>CAGTAACATCAACAGCAATAGAGCGATTTTTAAAGCGTATTAAAG<br>TTCCTTGTCATCGTATTGGAGACAAAGAAATTCATGTATTAGGTG<br>ACACTAAATCGGTTGTATTGTCTATGCCTATTAATGAACAGTTTA<br>AAAAAGCAGTCTTAAATGCTGCAAACGACGAAAACTACGCTTTAG<br>TAGCTTAATAAAACGCATGAGAAAGCCCCCGGAAGATCACCTTCC<br>GGGGGCTTTTTTATTGCGC | Modified from (*6*) |
| *luxR* | transcription unit | TGCCCGCTCGCGAGCGCGTCCTCTATAGATTCCTCGAGGAGCGGA<br>TACTTCGTAGGGTAGACTCGGGTCCTTTATAGCTAGCTCAGCCCT<br>TGGTACAATGCTAGCTACTAGAGAAAGAGGAGAAATACTAGATGA<br>AAAACATAAATGCCGACGACACATACAGAATAATTAATAAAATTA<br>AAGCTTGTAGAAGCAATAATGATATTAATCAATGCTTATCTGATA<br>TGACTAAAATGGTACATTGTGAATATTATTTACTCGCGATCATTT<br>ATCCTCATTCTATGGTTAAATCTGATATTTCAATCCTAGATAATT | Modified from (*6*) |

| | | | |
|---|---|---|---|
| *sfgfp* | insulated gene | ACCCTAAAAAATGGAGGCAATATTATGATGACGCTAATTTAATAA<br>AATATGATCCTATAGTAGATTATTCTAACTCCAATCATTCACCAA<br>TTAATTGGAATATATTTGAAAACAATGCTGTAAATAAAAAATCTC<br>CAAATGTAATTAAAGAAGCGAAAACATCAGGTCTTATCACTGGGT<br>TTAGTTTCCCTATTCATACGGCTAACAATGGCTTCGGAATGCTTA<br>GTTTTGCACATTCAGAAAAAGACAACTATATAGATAGTTTATTTT<br>TACATGCGTGTATGAACATACCATTAATTGTTCCTTCTCTAGTTG<br>ATAATTATCGAAAAATAAATATAGCAAATAATAAATCAAACAACG<br>ATTTAACCAAAAGAGAAAAAGAATGTTTAGCGTGGGCATGCGAAG<br>GAAAAAGCTCTTGGGATATTTCAAAAATATTAGGTTGCAGTGAGC<br>GTACTGTCACTTTCCATTTAACCAATGCGCAAATGAAACTCAATA<br>CAACAAACCGCTGCCAAAGTATTTCTAAAGCAATTTTAACAGGAG<br>CAATTGATTGCCCATACTTTAAAAATTAATAAAACGCATGAGAAA<br>GCCCCCGGAAGATCACCTTCCGGGGGCTTTTTTATTGCGC<br>AGCTGTCACCGGATGTGCTTTCCGGTCTGATGAGTCCGTGAGGAC<br>GAAACAGCCTCTACAAATAATTTTGTTTAATAAGTATCCTCTAAC<br>CCTAAAGGGGCACAAAATCATGCGTAAAGGCGAAGAGCTGTTCAC<br>TGGTGTCGTCCCTATTCTGGTGGAACTGGATGGTGATGTCAACGG<br>TCATAAGTTTTCCGTGCGTGGCGAGGGTGAAGGTGACGCAACTAA<br>TGGTAAACTGACGCTGAAGTTCATCTGTACTACTGGTAAACTGCC<br>GGTACCTTGGCCGACTCTGGTAACGACGCTGACTTATGGTGTTCA<br>GTGCTTTGCTCGTTATCCGGACCATATGAAGCAGCATGACTTCTT<br>CAAGTCCGCCATGCCGGAAGGCTATGTGCAGGAACGCACGATTTC<br>CTTTAAGGATGACGGCACGTACAAAACGCGTGCGGAAGTGAAATT<br>TGAAGGCGATACCCTGGTAAACCGCATTGAGCTGAAAGGCATTGA<br>CTTTAAAGAAGACGGCAATATCCTGGGCCATAAGCTGGAATACAA<br>TTTTAACAGCCACAATGTTTACATCACCGCCGATAAACAAAAAAA<br>TGGCATTAAAGCGAATTTTAAAATTCGCCACAACGTGGAGGATGG<br>CAGCGTGCAGCTGGCTGATCACTACCAGCAAAACACTCCAATCGG<br>TGATGGTCCTGTTCTGCTGCCAGACAATCACTATCTGAGCACGCA<br>AAGCGTTCTGTCTAAAGATCCGAACGAGAAACGCGATCATATGGT<br>TCTGCTGGAGTTCGTAACCGCAGCGGGCATCACGCATGGTATGGA<br>TGAACTGTACAAATGATGACCAGGCATCAAATAAAACGAAAGGCT<br>CAGTCGAAAGACTGGGCCTTTCGTTTTATCTGTTGTTTGTCGGTG<br>AACGCTCTCTACTAGAGTCACACTGGCTCACCTTCGGGTGGGCCT<br>TTCTGCGTTTATA | (*52*, *42*) |

**Table S16. Plasmids used in this study**. CmR = chloramphenicol, SpecR = spectinomycin, AmpR = ampicillin. Superscript u ($^u$) designates uninsulated promoters.

| Label | Type | Description | Origin of replication | Resistance cassette |
|---|---|---|---|---|
| pJS0115 | circuit | $P_{LtetO-1}{}^u$:S1 | ColE1 | CmR |
| pJS0120 | circuit | $P_{LtetO-1}{}^u$:S2 | ColE1 | CmR |
| pJS0114 | circuit | $P_{LtetO-1}{}^u$:S3 | ColE1 | CmR |
| pJS0118 | circuit | $P_{LtetO-1}{}^u$:S4 | ColE1 | CmR |
| pJS0119 | circuit | $P_{LtetO-1}{}^u$:S5 | ColE1 | CmR |
| pJS0107 | circuit | $P_{LtetO-1}{}^u$:S6 | ColE1 | CmR |
| pJS0116 | circuit | $P_{LtetO-1}{}^u$:S7 | ColE1 | CmR |
| pJS0113 | circuit | $P_{LtetO-1}{}^u$:S8 | ColE1 | CmR |
| pJS0117 | circuit | $P_{LtetO-1}{}^u$:S9 | ColE1 | CmR |
| pJS0101 | probe | P1$^u$:*sfgfp* | p15A | SpecR |
| pJS0106 | probe | P2$^u$:*sfgfp* | p15A | SpecR |
| pJS0100 | probe | P3$^u$:*sfgfp* | p15A | SpecR |
| pJS0104 | probe | P4$^u$:*sfgfp* | p15A | SpecR |
| pJS0105 | probe | P5$^u$:*sfgfp* | p15A | SpecR |
| pJS0087 | probe | P6$^u$:*sfgfp* | p15A | SpecR |
| pJS0102 | probe | P7$^u$:*sfgfp* | p15A | SpecR |
| pJS0099 | probe | P8$^u$:*sfgfp* | p15A | SpecR |
| pJS0103 | probe | P9$^u$:*sfgfp* | p15A | SpecR |
| pSC31_1 | dCas9 | Weak constitutive dCas9 expression | pSC101* | AmpR |
| pJS0344 | circuit | aTc-NOT1 | ColE1 | CmR |
| pJS0349 | circuit | aTc-NOT2 | ColE1 | CmR |
| pJS0343 | circuit | aTc-NOT3 | ColE1 | CmR |
| pJS0347 | circuit | aTc-NOT4 | ColE1 | CmR |
| pJS0348 | circuit | aTc-NOT5 | ColE1 | CmR |
| pJS0341 | circuit | aTc-NOT6 | ColE1 | CmR |
| pJS0345 | circuit | aTc-NOT7 | ColE1 | CmR |
| pJS0342 | circuit | aTc-NOT8 | ColE1 | CmR |
| pJS0346 | circuit | aTc-NOT9 | ColE1 | CmR |
| pJS0275 | probe | $P_{tet}$:*sfgfp* | p15A | SpecR |
| pJS0307 | probe | P1:*sfgfp* | p15A | SpecR |
| pJS0312 | probe | P2:*sfgfp* | p15A | SpecR |
| pJS0306 | probe | P3:*sfgfp* | p15A | SpecR |
| pJS0310 | probe | P4:*sfgfp* | p15A | SpecR |
| pJS0311 | probe | P5:*sfgfp* | p15A | SpecR |
| pJS0337 | probe | P6:*sfgfp* | p15A | SpecR |
| pJS0308 | probe | P7:*sfgfp* | p15A | SpecR |
| pJS0305 | probe | P8:*sfgfp* | p15A | SpecR |
| pJS0309 | probe | P9:*sfgfp* | p15A | SpecR |
| pSC31_3 | dCas9 | Strong constitutive dCas9 expression | pSC101* | AmpR |
| pJS0143 | circuit | Empty circuit plasmid | ColE1 | CmR |
| pJS0130 | probe | Empty probe plasmid | p15A | SpecR |
| pJS0122 | circuit | MUX (P1=0, P9=0, P4=0) | ColE1 | CmR |
| pJS0123 | circuit | MUX (P1=0, P9=1, P4=0) | ColE1 | CmR |
| pJS0156 | circuit | MUX (P1=1, P9=0, P4=0) | ColE1 | CmR |

| | | | | |
|---|---|---|---|---|
| pJS0157 | circuit | MUX (P1=1, P9=1, P4=0) | ColE1 | CmR |
| pJS0126 | circuit | MUX (P1=0, P9=0, P4=1) | ColE1 | CmR |
| pJS0127 | circuit | MUX (P1=0, P9=1, P4=1) | ColE1 | CmR |
| pJS0158 | circuit | MUX (P1=1, P9=0, P4=1) | ColE1 | CmR |
| pJS0155 | circuit | MUX (P1=1, P9=1, P4=1) | ColE1 | CmR |
| pJS0162 | circuit | DEMUX ($P_R$=0, P3=0) | ColE1 | CmR |
| pJS0133 | circuit | DEMUX ($P_R$=1, P3=0) | ColE1 | CmR |
| pJS0164 | circuit | DEMUX ($P_R$=0, P3=1) | ColE1 | CmR |
| pJS0134 | circuit | DEMUX ($P_R$=1, P3=1) | ColE1 | CmR |
| pJS0002 | probe | $P_R$:*sfgfp* | p15A | SpecR |
| pJS0338 | circuit | aTc sensor | ColE1 | CmR |
| pJS0339 | circuit | IPTG sensor | ColE1 | CmR |
| pJS0340 | circuit | DAPG sensor | ColE1 | CmR |
| pJS0260 | probe | $P_{tac}$:*sfgfp* | p15A | SpecR |
| pJS0304 | probe | $P_{PhlF}$:*sfgfp* | p15A | SpecR |
| pJS0277 | circuit | SENSOR-MUX | ColE1 | CmR |
| pJS0350 | circuit | aTc-NOT6* | ColE1 | CmR |
| pJS0200 | probe | P6*:*sfgfp* | p15A | SpecR |
| pJS0301 | circuit | SENSOR-MUX* | ColE1 | CmR |
| pJS0318 | circuit | $SENDER_{BCD22}$ | ColE1 | CmR |
| pJS0317 | circuit | $SENDER_{B0031}$ | ColE1 | CmR |
| pJS0315 | circuit | $SENDER_{B0030}$ | ColE1 | CmR |
| pJS0286 | circuit | $SENDER_{B0034}$ | ColE1 | CmR |
| pJS0285 | circuit | $RECEIVER_{J23117}$ | ColE1 | CmR |
| pJS0284 | circuit | $RECEIVER_{J23115}$ | ColE1 | CmR |
| pJS0283 | circuit | $RECEIVER_{J23105}$ | ColE1 | CmR |
| pJS0282 | circuit | $RECEIVER_{J23107}$ | ColE1 | CmR |
| pJS0205 | marker | P2:*mCherry* | p15A | SpecR |
| pJS0281 | probe | $P_{lux}$:*sfgfp* | p15A | SpecR |
| pJS0314 | circuit | AHL-DEMUX | ColE1 | CmR |
| pJS0321 | circuit | SENSOR-MUX*-AHL | ColE1 | CmR |

**Table S17. Bacterial strains used in this study.** All strains were derived from *E. coli* K-12 MG1655. Superscript u ($^u$) designates uninsulated promoters.

| Label | DNA content† | Description | Figure(s) |
|---|---|---|---|
| sJS0329 | pJS0115, pJS0101, pSC31_1 | $P_{LtetO-1}^u$:S1, P1$^u$:*sfgfp* | 2C |
| sJS0334 | pJS0120, pJS0101, pSC31_1 | $P_{LtetO-1}^u$:S2, P1$^u$:*sfgfp* | 2C |
| sJS0328 | pJS0114, pJS0101, pSC31_1 | $P_{LtetO-1}^u$:S3, P1$^u$:*sfgfp* | 2C |
| sJS0332 | pJS0118, pJS0101, pSC31_1 | $P_{LtetO-1}^u$:S4, P1$^u$:*sfgfp* | 2C |
| sJS0333 | pJS0119, pJS0101, pSC31_1 | $P_{LtetO-1}^u$:S5, P1$^u$:*sfgfp* | 2C |
| sJS0321 | pJS0107, pJS0101, pSC31_1 | $P_{LtetO-1}^u$:S6, P1$^u$:*sfgfp* | 2C |
| sJS0330 | pJS0116, pJS0101, pSC31_1 | $P_{LtetO-1}^u$:S7, P1$^u$:*sfgfp* | 2C |
| sJS0327 | pJS0113, pJS0101, pSC31_1 | $P_{LtetO-1}^u$:S8, P1$^u$:*sfgfp* | 2C |
| sJS0331 | pJS0117, pJS0101, pSC31_1 | $P_{LtetO-1}^u$:S9, P1$^u$:*sfgfp* | 2C |
| sJS0399 | pJS0115, pJS0106, pSC31_1 | $P_{LtetO-1}^u$:S1, P2$^u$:*sfgfp* | 2C |
| sJS0404 | pJS0120, pJS0106, pSC31_1 | $P_{LtetO-1}^u$:S2, P2$^u$:*sfgfp* | 2C |
| sJS0398 | pJS0114, pJS0106, pSC31_1 | $P_{LtetO-1}^u$:S3, P2$^u$:*sfgfp* | 2C |
| sJS0402 | pJS0118, pJS0106, pSC31_1 | $P_{LtetO-1}^u$:S4, P2$^u$:*sfgfp* | 2C |
| sJS0403 | pJS0119, pJS0106, pSC31_1 | $P_{LtetO-1}^u$:S5, P2$^u$:*sfgfp* | 2C |
| sJS0391 | pJS0107, pJS0106, pSC31_1 | $P_{LtetO-1}^u$:S6, P2$^u$:*sfgfp* | 2C |
| sJS0400 | pJS0116, pJS0106, pSC31_1 | $P_{LtetO-1}^u$:S7, P2$^u$:*sfgfp* | 2C |
| sJS0397 | pJS0113, pJS0106, pSC31_1 | $P_{LtetO-1}^u$:S8, P2$^u$:*sfgfp* | 2C |
| sJS0401 | pJS0117, pJS0106, pSC31_1 | $P_{LtetO-1}^u$:S9, P2$^u$:*sfgfp* | 2C |
| sJS0315 | pJS0115, pJS0100, pSC31_1 | $P_{LtetO-1}^u$:S1, P3$^u$:*sfgfp* | 2C |
| sJS0320 | pJS0120, pJS0100, pSC31_1 | $P_{LtetO-1}^u$:S2, P3$^u$:*sfgfp* | 2C |
| sJS0314 | pJS0114, pJS0100, pSC31_1 | $P_{LtetO-1}^u$:S3, P3$^u$:*sfgfp* | 2C |
| sJS0318 | pJS0118, pJS0100, pSC31_1 | $P_{LtetO-1}^u$:S4, P3$^u$:*sfgfp* | 2C |
| sJS0319 | pJS0119, pJS0100, pSC31_1 | $P_{LtetO-1}^u$:S5, P3$^u$:*sfgfp* | 2C |
| sJS0307 | pJS0107, pJS0100, pSC31_1 | $P_{LtetO-1}^u$:S6, P3$^u$:*sfgfp* | 2C |
| sJS0316 | pJS0116, pJS0100, pSC31_1 | $P_{LtetO-1}^u$:S7, P3$^u$:*sfgfp* | 2C |
| sJS0313 | pJS0113, pJS0100, pSC31_1 | $P_{LtetO-1}^u$:S8, P3$^u$:*sfgfp* | 2C |
| sJS0317 | pJS0117, pJS0100, pSC31_1 | $P_{LtetO-1}^u$:S9, P3$^u$:*sfgfp* | 2C |
| sJS0371 | pJS0115, pJS0104, pSC31_1 | $P_{LtetO-1}^u$:S1, P4$^u$:*sfgfp* | 2C |
| sJS0376 | pJS0120, pJS0104, pSC31_1 | $P_{LtetO-1}^u$:S2, P4$^u$:*sfgfp* | 2C |
| sJS0370 | pJS0114, pJS0104, pSC31_1 | $P_{LtetO-1}^u$:S3, P4$^u$:*sfgfp* | 2C |
| sJS0374 | pJS0118, pJS0104, pSC31_1 | $P_{LtetO-1}^u$:S4, P4$^u$:*sfgfp* | 2C |
| sJS0375 | pJS0119, pJS0104, pSC31_1 | $P_{LtetO-1}^u$:S5, P4$^u$:*sfgfp* | 2C |
| sJS0363 | pJS0107, pJS0104, pSC31_1 | $P_{LtetO-1}^u$:S6, P4$^u$:*sfgfp* | 2C |
| sJS0372 | pJS0116, pJS0104, pSC31_1 | $P_{LtetO-1}^u$:S7, P4$^u$:*sfgfp* | 2C |
| sJS0369 | pJS0113, pJS0104, pSC31_1 | $P_{LtetO-1}^u$:S8, P4$^u$:*sfgfp* | 2C |
| sJS0373 | pJS0117, pJS0104, pSC31_1 | $P_{LtetO-1}^u$:S9, P4$^u$:*sfgfp* | 2C |
| sJS0385 | pJS0115, pJS0105, pSC31_1 | $P_{LtetO-1}^u$:S1, P5$^u$:*sfgfp* | 2C |
| sJS0390 | pJS0120, pJS0105, pSC31_1 | $P_{LtetO-1}^u$:S2, P5$^u$:*sfgfp* | 2C |
| sJS0384 | pJS0114, pJS0105, pSC31_1 | $P_{LtetO-1}^u$:S3, P5$^u$:*sfgfp* | 2C |
| sJS0388 | pJS0118, pJS0105, pSC31_1 | $P_{LtetO-1}^u$:S4, P5$^u$:*sfgfp* | 2C |
| sJS0389 | pJS0119, pJS0105, pSC31_1 | $P_{LtetO-1}^u$:S5, P5$^u$:*sfgfp* | 2C |
| sJS0377 | pJS0107, pJS0105, pSC31_1 | $P_{LtetO-1}^u$:S6, P5$^u$:*sfgfp* | 2C |
| sJS0386 | pJS0116, pJS0105, pSC31_1 | $P_{LtetO-1}^u$:S7, P5$^u$:*sfgfp* | 2C |
| sJS0383 | pJS0113, pJS0105, pSC31_1 | $P_{LtetO-1}^u$:S8, P5$^u$:*sfgfp* | 2C |
| sJS0387 | pJS0117, pJS0105, pSC31_1 | $P_{LtetO-1}^u$:S9, P5$^u$:*sfgfp* | 2C |
| sJS0217 | pJS0115, pJS0087, pSC31_1 | $P_{LtetO-1}^u$:S1, P6$^u$:*sfgfp* | 2C |
| sJS0222 | pJS0120, pJS0087, pSC31_1 | $P_{LtetO-1}^u$:S2, P6$^u$:*sfgfp* | 2C |
| sJS0216 | pJS0114, pJS0087, pSC31_1 | $P_{LtetO-1}^u$:S3, P6$^u$:*sfgfp* | 2C |
| sJS0220 | pJS0118, pJS0087, pSC31_1 | $P_{LtetO-1}^u$:S4, P6$^u$:*sfgfp* | 2C |
| sJS0221 | pJS0119, pJS0087, pSC31_1 | $P_{LtetO-1}^u$:S5, P6$^u$:*sfgfp* | 2C |
| sJS0209 | pJS0107, pJS0087, pSC31_1 | $P_{LtetO-1}^u$:S6, P6$^u$:*sfgfp* | 2C |

| | | | |
|---|---|---|---|
| sJS0218 | pJS0116, pJS0087, pSC31_1 | $P_{LtetO-1}^{u}$:S7, $P6^{u}$:*sfgfp* | 2C |
| sJS0215 | pJS0113, pJS0087, pSC31_1 | $P_{LtetO-1}^{u}$:S8, $P6^{u}$:*sfgfp* | 2C |
| sJS0219 | pJS0117, pJS0087, pSC31_1 | $P_{LtetO-1}^{u}$:S9, $P6^{u}$:*sfgfp* | 2C |
| sJS0343 | pJS0115, pJS0102, pSC31_1 | $P_{LtetO-1}^{u}$:S1, $P7^{u}$:*sfgfp* | 2C |
| sJS0348 | pJS0120, pJS0102, pSC31_1 | $P_{LtetO-1}^{u}$:S2, $P7^{u}$:*sfgfp* | 2C |
| sJS0342 | pJS0114, pJS0102, pSC31_1 | $P_{LtetO-1}^{u}$:S3, $P7^{u}$:*sfgfp* | 2C |
| sJS0346 | pJS0118, pJS0102, pSC31_1 | $P_{LtetO-1}^{u}$:S4, $P7^{u}$:*sfgfp* | 2C |
| sJS0347 | pJS0119, pJS0102, pSC31_1 | $P_{LtetO-1}^{u}$:S5, $P7^{u}$:*sfgfp* | 2C |
| sJS0335 | pJS0107, pJS0102, pSC31_1 | $P_{LtetO-1}^{u}$:S6, $P7^{u}$:*sfgfp* | 2C |
| sJS0344 | pJS0116, pJS0102, pSC31_1 | $P_{LtetO-1}^{u}$:S7, $P7^{u}$:*sfgfp* | 2C |
| sJS0341 | pJS0113, pJS0102, pSC31_1 | $P_{LtetO-1}^{u}$:S8, $P7^{u}$:*sfgfp* | 2C |
| sJS0345 | pJS0117, pJS0102, pSC31_1 | $P_{LtetO-1}^{u}$:S9, $P7^{u}$:*sfgfp* | 2C |
| sJS0301 | pJS0115, pJS0099, pSC31_1 | $P_{LtetO-1}^{u}$:S1, $P8^{u}$:*sfgfp* | 2C |
| sJS0306 | pJS0120, pJS0099, pSC31_1 | $P_{LtetO-1}^{u}$:S2, $P8^{u}$:*sfgfp* | 2C |
| sJS0300 | pJS0114, pJS0099, pSC31_1 | $P_{LtetO-1}^{u}$:S3, $P8^{u}$:*sfgfp* | 2C |
| sJS0304 | pJS0118, pJS0099, pSC31_1 | $P_{LtetO-1}^{u}$:S4, $P8^{u}$:*sfgfp* | 2C |
| sJS0305 | pJS0119, pJS0099, pSC31_1 | $P_{LtetO-1}^{u}$:S5, $P8^{u}$:*sfgfp* | 2C |
| sJS0293 | pJS0107, pJS0099, pSC31_1 | $P_{LtetO-1}^{u}$:S6, $P8^{u}$:*sfgfp* | 2C |
| sJS0302 | pJS0116, pJS0099, pSC31_1 | $P_{LtetO-1}^{u}$:S7, $P8^{u}$:*sfgfp* | 2C |
| sJS0299 | pJS0113, pJS0099, pSC31_1 | $P_{LtetO-1}^{u}$:S8, $P8^{u}$:*sfgfp* | 2C |
| sJS0303 | pJS0117, pJS0099, pSC31_1 | $P_{LtetO-1}^{u}$:S9, $P8^{u}$:*sfgfp* | 2C |
| sJS0357 | pJS0115, pJS0103, pSC31_1 | $P_{LtetO-1}^{u}$:S1, $P9^{u}$:*sfgfp* | 2C |
| sJS0362 | pJS0120, pJS0103, pSC31_1 | $P_{LtetO-1}^{u}$:S2, $P9^{u}$:*sfgfp* | 2C |
| sJS0356 | pJS0114, pJS0103, pSC31_1 | $P_{LtetO-1}^{u}$:S3, $P9^{u}$:*sfgfp* | 2C |
| sJS0360 | pJS0118, pJS0103, pSC31_1 | $P_{LtetO-1}^{u}$:S4, $P9^{u}$:*sfgfp* | 2C |
| sJS0361 | pJS0119, pJS0103, pSC31_1 | $P_{LtetO-1}^{u}$:S5, $P9^{u}$:*sfgfp* | 2C |
| sJS0349 | pJS0107, pJS0103, pSC31_1 | $P_{LtetO-1}^{u}$:S6, $P9^{u}$:*sfgfp* | 2C |
| sJS0358 | pJS0116, pJS0103, pSC31_1 | $P_{LtetO-1}^{u}$:S7, $P9^{u}$:*sfgfp* | 2C |
| sJS0355 | pJS0113, pJS0103, pSC31_1 | $P_{LtetO-1}^{u}$:S8, $P9^{u}$:*sfgfp* | 2C |
| sJS0359 | pJS0117, pJS0103, pSC31_1 | $P_{LtetO-1}^{u}$:S9, $P9^{u}$:*sfgfp* | 2C |
| sJS0061 | pSC31_1 | Autofluorescence control | 2C |
| sJS1213 | pJS0344, pJS0275, pSC31_3 | aTc-NOT1, $P_{tet}$:*sfgfp* | 2E |
| sJS1106 | pJS0344, pJS0307, pSC31_3 | aTc-NOT1, P1:*sfgfp* | 2E |
| sJS1218 | pJS0349, pJS0275, pSC31_3 | aTc-NOT2, $P_{tet}$:*sfgfp* | 2E |
| sJS1111 | pJS0349, pJS0312, pSC31_3 | aTc-NOT2, P2:*sfgfp* | 2E |
| sJS1212 | pJS0343, pJS0275, pSC31_3 | aTc-NOT3, $P_{tet}$:*sfgfp* | 2E |
| sJS1105 | pJS0343, pJS0306, pSC31_3 | aTc-NOT3, P3:*sfgfp* | 2E |
| sJS1216 | pJS0347, pJS0275, pSC31_3 | aTc-NOT4, $P_{tet}$:*sfgfp* | 2E |
| sJS1109 | pJS0347, pJS0310, pSC31_3 | aTc-NOT4, P4:*sfgfp* | 2E |
| sJS1217 | pJS0348, pJS0275, pSC31_3 | aTc-NOT5, $P_{tet}$:*sfgfp* | 2E |
| sJS1110 | pJS0348, pJS0311, pSC31_3 | aTc-NOT5, P5:*sfgfp* | 2E |
| sJS1210 | pJS0341, pJS0275, pSC31_3 | aTc-NOT6, $P_{tet}$:*sfgfp* | 2E |
| sJS1103 | pJS0341, pJS0337, pSC31_3 | aTc-NOT6, P6:*sfgfp* | 2E |
| sJS1214 | pJS0345, pJS0275, pSC31_3 | aTc-NOT7, $P_{tet}$:*sfgfp* | 2E |
| sJS1107 | pJS0345, pJS0308, pSC31_3 | aTc-NOT7, P7:*sfgfp* | 2E |
| sJS1211 | pJS0342, pJS0275, pSC31_3 | aTc-NOT8, $P_{tet}$:*sfgfp* | 2E |
| sJS1104 | pJS0342, pJS0305, pSC31_3 | aTc-NOT8, P8:*sfgfp* | 2E |
| sJS1215 | pJS0346, pJS0275, pSC31_3 | aTc-NOT9, $P_{tet}$:*sfgfp* | 2E |
| sJS1108 | pJS0346, pJS0309, pSC31_3 | aTc-NOT9, P9:*sfgfp* | 2E |
| sJS1015 | pJS0143, pJS0307, pSC31_3 | P1:*sfgfp* | 2E, S6, S8 |
| sJS1020 | pJS0143, pJS0312, pSC31_3 | P2:*sfgfp* | 2E, 3, S6, S8, S10, 4, S12, S13 |
| sJS1014 | pJS0143, pJS0306, pSC31_3 | P3:*sfgfp* | 2E, 3A, S6, S8, S10, 4, S13 |
| sJS1018 | pJS0143, pJS0310, pSC31_3 | P4:*sfgfp* | 2E, S6, S8, S13 |
| sJS1019 | pJS0143, pJS0311, pSC31_3 | P5:*sfgfp* | 2E, 3A, S6, S8 |

| | | | |
|---|---|---|---|
| sJS1092 | pJS0143, pJS0337, pSC31_3 | P6:*sfgfp* | 2E, 3A, S6 |
| sJS1016 | pJS0143, pJS0308, pSC31_3 | P7:*sfgfp* | 2E, 3B, S10, 4, S12, S13 |
| sJS1013 | pJS0143, pJS0305, pSC31_3 | P8:*sfgfp* | 2E, 3B, S10, S13 |
| sJS1017 | pJS0143, pJS0309, pSC31_3 | P9:*sfgfp* | 2E, 3B, S6, S8, S10 |
| sJS1007 | pJS0143, pJS0130, pSC31_3 | Autofluorescence control | 2E, 3, S5-S10, S12, S13, 4 |
| sJS1132 | pJS0122, pJS0311, pSC31_3 | MUX (P1=0, P9=0, P4=0), P5:*sfgfp* | 3A |
| sJS1133 | pJS0123, pJS0311, pSC31_3 | MUX (P1=0, P9=1, P4=0), P5:*sfgfp* | 3A |
| sJS1137 | pJS0156, pJS0311, pSC31_3 | MUX (P1=1, P9=0, P4=0), P5:*sfgfp* | 3A |
| sJS1138 | pJS0157, pJS0311, pSC31_3 | MUX (P1=1, P9=1, P4=0), P5:*sfgfp* | 3A |
| sJS1134 | pJS0126, pJS0311, pSC31_3 | MUX (P1=0, P9=0, P4=1), P5:*sfgfp* | 3A |
| sJS1135 | pJS0127, pJS0311, pSC31_3 | MUX (P1=0, P9=1, P4=1), P5:*sfgfp* | 3A |
| sJS1139 | pJS0158, pJS0311, pSC31_3 | MUX (P1=1, P9=0, P4=1), P5:*sfgfp* | 3A |
| sJS1136 | pJS0155, pJS0311, pSC31_3 | MUX (P1=1, P9=1, P4=1), P5:*sfgfp* | 3A |
| sJS1124 | pJS0122, pJS0306, pSC31_3 | MUX (P1=0, P9=0, P4=0), P3:*sfgfp* | 3A |
| sJS1125 | pJS0123, pJS0306, pSC31_3 | MUX (P1=0, P9=1, P4=0), P3:*sfgfp* | 3A |
| sJS1129 | pJS0156, pJS0306, pSC31_3 | MUX (P1=1, P9=0, P4=0), P3:*sfgfp* | 3A |
| sJS1130 | pJS0157, pJS0306, pSC31_3 | MUX (P1=1, P9=1, P4=0), P3:*sfgfp* | 3A |
| sJS1126 | pJS0126, pJS0306, pSC31_3 | MUX (P1=0, P9=0, P4=1), P3:*sfgfp* | 3A |
| sJS1127 | pJS0127, pJS0306, pSC31_3 | MUX (P1=0, P9=1, P4=1), P3:*sfgfp* | 3A |
| sJS1131 | pJS0158, pJS0306, pSC31_3 | MUX (P1=1, P9=0, P4=1), P3:*sfgfp* | 3A |
| sJS1128 | pJS0155, pJS0306, pSC31_3 | MUX (P1=1, P9=1, P4=1), P3:*sfgfp* | 3A |
| sJS1148 | pJS0122, pJS0312, pSC31_3 | MUX (P1=0, P9=0, P4=0), P2:*sfgfp* | 3A |
| sJS1149 | pJS0123, pJS0312, pSC31_3 | MUX (P1=0, P9=1, P4=0), P2:*sfgfp* | 3A |
| sJS1153 | pJS0156, pJS0312, pSC31_3 | MUX (P1=1, P9=0, P4=0), P2:*sfgfp* | 3A |
| sJS1154 | pJS0157, pJS0312, pSC31_3 | MUX (P1=1, P9=1, P4=0), P2:*sfgfp* | 3A |
| sJS1150 | pJS0126, pJS0312, pSC31_3 | MUX (P1=0, P9=0, P4=1), P2:*sfgfp* | 3A |
| sJS1151 | pJS0127, pJS0312, pSC31_3 | MUX (P1=0, P9=1, P4=1), P2:*sfgfp* | 3A |
| sJS1155 | pJS0158, pJS0312, pSC31_3 | MUX (P1=1, P9=0, P4=1), P2:*sfgfp* | 3A |
| sJS1152 | pJS0155, pJS0312, pSC31_3 | MUX (P1=1, P9=1, P4=1), P2:*sfgfp* | 3A |
| sJS1140 | pJS0122, pJS0337, pSC31_3 | MUX (P1=0, P9=0, P4=0), P6:*sfgfp* | 3A |
| sJS1141 | pJS0123, pJS0337, pSC31_3 | MUX (P1=0, P9=1, P4=0), P6:*sfgfp* | 3A |
| sJS1145 | pJS0156, pJS0337, pSC31_3 | MUX (P1=1, P9=0, P4=0), P6:*sfgfp* | 3A |
| sJS1146 | pJS0157, pJS0337, pSC31_3 | MUX (P1=1, P9=1, P4=0), P6:*sfgfp* | 3A |
| sJS1142 | pJS0126, pJS0337, pSC31_3 | MUX (P1=0, P9=0, P4=1), P6:*sfgfp* | 3A |
| sJS1143 | pJS0127, pJS0337, pSC31_3 | MUX (P1=0, P9=1, P4=1), P6:*sfgfp* | 3A |
| sJS1147 | pJS0158, pJS0337, pSC31_3 | MUX (P1=1, P9=0, P4=1), P6:*sfgfp* | 3A |
| sJS1144 | pJS0155, pJS0337, pSC31_3 | MUX (P1=1, P9=1, P4=1), P6:*sfgfp* | 3A |
| sJS1161 | pJS0162, pJS0305, pSC31_3 | DEMUX ($P_R$=0, P3=0), P8:*sfgfp* | 3B |
| sJS1176 | pJS0133, pJS0305, pSC31_3 | DEMUX ($P_R$=1, P3=0), P8:*sfgfp* | 3B |
| sJS1163 | pJS0164, pJS0305, pSC31_3 | DEMUX ($P_R$=0, P3=1), P8:*sfgfp* | 3B |
| sJS1177 | pJS0134, pJS0305, pSC31_3 | DEMUX ($P_R$=1, P3=1), P8:*sfgfp* | 3B |
| sJS1169 | pJS0162, pJS0308, pSC31_3 | DEMUX ($P_R$=0, P3=0), P7:*sfgfp* | 3B |
| sJS1180 | pJS0133, pJS0308, pSC31_3 | DEMUX ($P_R$=1, P3=0), P7:*sfgfp* | 3B |
| sJS1171 | pJS0164, pJS0308, pSC31_3 | DEMUX ($P_R$=0, P3=1), P7:*sfgfp* | 3B |
| sJS1181 | pJS0134, pJS0308, pSC31_3 | DEMUX ($P_R$=1, P3=1), P7:*sfgfp* | 3B |
| sJS1165 | pJS0162, pJS0309, pSC31_3 | DEMUX ($P_R$=0, P3=0), P9:*sfgfp* | 3B |
| sJS1178 | pJS0133, pJS0309, pSC31_3 | DEMUX ($P_R$=1, P3=0), P9:*sfgfp* | 3B |
| sJS1167 | pJS0164, pJS0309, pSC31_3 | DEMUX ($P_R$=0, P3=1), P9:*sfgfp* | 3B |
| sJS1179 | pJS0134, pJS0309, pSC31_3 | DEMUX ($P_R$=1, P3=1), P9:*sfgfp* | 3B |
| sJS1173 | pJS0162, pJS0312, pSC31_3 | DEMUX ($P_R$=0, P3=0), P2:*sfgfp* | 3B |
| sJS1182 | pJS0133, pJS0312, pSC31_3 | DEMUX ($P_R$=1, P3=0), P2:*sfgfp* | 3B |
| sJS1175 | pJS0164, pJS0312, pSC31_3 | DEMUX ($P_R$=0, P3=1), P2:*sfgfp* | 3B |
| sJS1183 | pJS0134, pJS0312, pSC31_3 | DEMUX ($P_R$=1, P3=1), P2:*sfgfp* | 3B |

| | | | |
|---|---|---|---|
| sJS0595 | pJS0002, pSC31_3 | $P_{R}$:*sfgfp* | 3B |
| sJS1094 | pJS0338, pJS0275, pSC31_3 | aTc sensor, $P_{tet}$:*sfgfp* | S5D |
| sJS1083 | pJS0339, pJS0260, pSC31_3 | IPTG sensor, $P_{tac}$:*sfgfp* | S5E |
| sJS1091 | pJS0340, pJS0304, pSC31_3 | DAPG sensor, $P_{PhlF}$:*sfgfp* | S5F |
| sJS1009 | pJS0143, pJS0275, pSC31_3 | $P_{tet}$:*sfgfp* | S5D, S6, S8 |
| sJS1010 | pJS0143, pJS0260, pSC31_3 | $P_{tac}$:*sfgfp* | S5E, S6, S8 |
| sJS1012 | pJS0143, pJS0304, pSC31_3 | $P_{PhlF}$:*sfgfp* | S5F, S6, S8, S10, S13 |
| sJS1201 | pJS0277, pJS0275, pSC31_3 | SENSOR-MUX, $P_{tet}$:*sfgfp* | S6 |
| sJS0830 | pJS0277, pJS0260, pSC31_3 | SENSOR-MUX, $P_{tac}$:*sfgfp* | S6 |
| sJS1202 | pJS0277, pJS0304, pSC31_3 | SENSOR-MUX, $P_{PhlF}$:*sfgfp* | S6 |
| sJS1203 | pJS0277, pJS0307, pSC31_3 | SENSOR-MUX, P1:*sfgfp* | S6 |
| sJS1204 | pJS0277, pJS0309, pSC31_3 | SENSOR-MUX, P9:*sfgfp* | S6 |
| sJS1205 | pJS0277, pJS0310, pSC31_3 | SENSOR-MUX, P4:*sfgfp* | S6 |
| sJS1208 | pJS0277, pJS0311, pSC31_3 | SENSOR-MUX, P5:*sfgfp* | S6 |
| sJS1209 | pJS0277, pJS0306, pSC31_3 | SENSOR-MUX, P3:*sfgfp* | S6 |
| sJS1206 | pJS0277, pJS0312, pSC31_3 | SENSOR-MUX, P2:*sfgfp* | S6 |
| sJS1207 | pJS0277, pJS0337, pSC31_3 | SENSOR-MUX, P6:*sfgfp* | S6 |
| sJS1219 | pJS0350, pJS0275, pSC31_3 | aTc-NOT6*, $P_{tet}$:*sfgfp* | S7 |
| sJS1112 | pJS0350, pJS0200, pSC31_3 | aTc-NOT6*, P6*:*sfgfp* | S7 |
| sJS1011 | pJS0143, pJS0200, pSC31_3 | P6*:*sfgfp* | S7, S8, S13 |
| sJS0986 | pJS0301, pJS0275, pSC31_3 | SENSOR-MUX*, $P_{tet}$:*sfgfp* | S8 |
| sJS0987 | pJS0301, pJS0260, pSC31_3 | SENSOR-MUX*, $P_{tac}$:*sfgfp* | S8 |
| sJS1194 | pJS0301, pJS0304, pSC31_3 | SENSOR-MUX*, $P_{PhlF}$:*sfgfp* | S8 |
| sJS1195 | pJS0301, pJS0307, pSC31_3 | SENSOR-MUX*, P1:*sfgfp* | S8 |
| sJS1196 | pJS0301, pJS0309, pSC31_3 | SENSOR-MUX*, P9:*sfgfp* | S8 |
| sJS1197 | pJS0301, pJS0310, pSC31_3 | SENSOR-MUX*, P4:*sfgfp* | S8 |
| sJS1199 | pJS0301, pJS0311, pSC31_3 | SENSOR-MUX*, P5:*sfgfp* | S8 |
| sJS1200 | pJS0301, pJS0306, pSC31_3 | SENSOR-MUX*, P3:*sfgfp* | S8 |
| sJS1198 | pJS0301, pJS0312, pSC31_3 | SENSOR-MUX*, P2:*sfgfp* | S8 |
| sJS0993 | pJS0301, pJS0200, pSC31_3 | SENSOR-MUX*, P6*:*sfgfp* | S8 |
| sJS1051 | pJS0318, pJS0205, pSC31_3 | SENDER$_{BCD22}$, P2:*mCherry* | S9 |
| sJS1050 | pJS0317, pJS0205, pSC31_3 | SENDER$_{B0031}$, P2:*mCherry* | S9 |
| sJS1049 | pJS0315, pJS0205, pSC31_3 | SENDER$_{B0030}$, P2:*mCherry* | S9 |
| sJS0866 | pJS0286, pJS0205, pSC31_3 | SENDER$_{B0034}$, P2:*mCherry* | S9 |
| sJS0865 | pJS0285, pJS0281, pSC31_3 | RECEIVER$_{J23117}$, $P_{lux}$:*sfgfp* | S9 |
| sJS0864 | pJS0284, pJS0281, pSC31_3 | RECEIVER$_{J23115}$, $P_{lux}$:*sfgfp* | S9, S10, S12, S13 |
| sJS0863 | pJS0283, pJS0281, pSC31_3 | RECEIVER$_{J23105}$, $P_{lux}$:*sfgfp* | S9 |
| sJS0862 | pJS0282, pJS0281, pSC31_3 | RECEIVER$_{J23107}$, $P_{lux}$:*sfgfp* | S9 |
| sJS1073 | pJS0314, pJS0281, pSC31_3 | AHL-DEMUX, $P_{lux}$:*sfgfp* | S10, S12, S13 |
| sJS1070 | pJS0314, pJS0304, pSC31_3 | AHL-DEMUX, $P_{PhlF}$:*sfgfp* | S10 |
| sJS1071 | pJS0314, pJS0306, pSC31_3 | AHL-DEMUX, P3:*sfgfp* | S10 |
| sJS1074 | pJS0314, pJS0305, pSC31_3 | AHL-DEMUX, P8:*sfgfp* | S10, S13 |
| sJS1037 | pJS0314, pJS0308, pSC31_3 | AHL-DEMUX, P7:*sfgfp* | S10, 4, S12, S13 |
| sJS1072 | pJS0314, pJS0309, pSC31_3 | AHL-DEMUX, P9:*sfgfp* | S10 |
| sJS1038 | pJS0314, pJS0312, pSC31_3 | AHL-DEMUX, P2:*sfgfp* | S10, 4, S12 |
| sJS1069 | pJS0321, pJS0306, pSC31_3 | SENSOR-MUX*-AHL, P3:*sfgfp* | 4, S13 |

| | | | |
|---|---|---|---|
| sJS1222 | pJS0321, pJS0304, pSC31_3 | SENSOR-MUX*-AHL, $P_{PhlF}$:*sfgfp* | S13 |
| sJS1223 | pJS0321, pJS0310, pSC31_3 | SENSOR-MUX*-AHL, P4:*sfgfp* | S13 |
| sJS1224 | pJS0321, pJS0312, pSC31_3 | SENSOR-MUX*-AHL, P2:*sfgfp* | S13 |
| sJS1225 | pJS0321, pJS0200, pSC31_3 | SENSOR-MUX*-AHL, P6*:*sfgfp* | S13 |
